# The N-terminal domain of MX1 proteins is essential for their antiviral activity against different families of RNA viruses

**DOI:** 10.1101/2022.05.10.491305

**Authors:** Joe McKellar, Mary Arnaud-Arnould, Laurent Chaloin, Marine Tauziet, Charlotte Arpin-André, Oriane Pourcelot, Mickael Blaise, Olivier Moncorgé, Caroline Goujon

## Abstract

Myxovirus resistance protein 1 (MX1) and MX2, are homologous, dynamin-like large GTPases, induced upon interferon (IFN) exposure. Human MX1 (HsMX1) is known to inhibit many viruses, including influenza A virus (IAV), by likely acting at various steps of their life cycles. Despite decades of studies, the mechanism(s) of action with which MX1 proteins manage to inhibit target viruses is not fully understood. MX1 proteins are mechano-enzymes and share a similar organization to dynamin, with an amino-terminal GTPase domain and a carboxy-terminal stalk domain, connected by a Bundle Signalling Element (BSE). These three elements are known to be essential for antiviral activity. HsMX1 has two unstructured regions, the L4 loop, also essential for antiviral activity, and a short amino (N)-terminal region, which greatly varies between MX1 proteins of different species. The role of this N-terminal domain in antiviral activity is not known. Herein, using mutagenesis, imaging and biochemical approaches, we demonstrate that the N-terminal domain of HsMX1 is essential for antiviral activity against IAV and Vesicular Stomatitis Virus (VSV), and for the ability to aggregate Orthobunyavirus nucleoproteins. Furthermore, we pinpoint a highly conserved leucine within this region, which is absolutely crucial for human, mouse and bat MX1 protein antiviral activity. Importantly, mutation of this leucine does not compromise GTPase activity or oligomerization capabilities, but does modify MX1 protein subcellular localisation. The discovery of this essential and highly conserved residue defines this region as key and may reveal insights as to the mechanism of action of MX1 proteins.

## Introduction

Influenza A virus (IAV) is a member of the *Orthomyxoviridae* family and the causative agent of the disease commonly known as the flu. Upon infection of target epithelial cells within the respiratory tract, IAV is sensed by pattern recognition receptors, including retinoic acid-inducible gene I (RIG-I), which induce a signalling cascade leading to the production and secretion of type 1 and type 3 interferons (IFNs). The IFNs act in a paracrine and autocrine manner and, through binding to their cognate receptors and activation of the Janus Kinase (JAK) / Signal transducer and activator of transcription (STAT) pathway, leads to the regulation of hundreds of IFN stimulated genes (ISGs). Among these ISGs, many antiviral restriction factors have been described (1). These factors establish a so-called antiviral state, powerfully limiting IAV replication (2). The study of the IFN response against IAV in mice led to the discovery of the Myxovirus resistance (MX) proteins, which were later identified in humans (3–6) . There are two homologous MX genes in humans, MX1 and MX2. Human MX1, also called MxA (and hereafter referred to as HsMX1), inhibits a wide range of RNA and DNA viruses, replicating either in the cytoplasm, such as the bunyavirus La Crosse Virus (LACV) and the rhabdovirus VSV, or in the nucleus, such as IAV (7). Human MX2 (HsMX2), or MxB, has notably been shown to potently inhibit Human Immunodeficiency Virus 1 (HIV-1) and Herpes viruses (8–13). Interestingly, mice also possess two Mx genes: MmMx1 and MmMx2. The latter is more closely related to HsMX1 than the former, but, interestingly, MmMx1 is a more potent inhibitor of IAV than HsMX1 (14). However, to our knowledge, MmMx1 only restricts orthomyxoviruses, whereas HsMX1 is broadly antiviral and MmMx2 inhibits VSV and Hantaan River Virus (HTNV) (7). This could be due partly to the fact that MmMx1 is mainly localized inside the nucleus (15, 16), contrary to HsMX1 which is cytosolic.

Dynamin-like GTPases all share a very similar general organization and among members of this family of proteins, can present almost superimposable three-dimensional (3D) crystal structures (17, 18). The 3D structure of HsMX1 has been partially solved (19, 20) (Fig. 1A). HsMX1 sports a globular head, which contains the GTPase module, and possesses a stalk domain, attached to the head by a tripartite Bundle Signalling Element (BSE), composed of three separate alpha-helices (α-helices) that allow the fold back of the stalk towards the GTPase domain, which is akin to dynamin (21) (Fig. 1A). The major difference between HsMX1 and dynamin is the lack in the former of the Pleckstrin Homology (PH) domain, essential for the PI(4,5)P2 binding capacity of dynamin (22). In place of the PH domain, HsMX1 has a flexible loop, termed loop L4, of which the structure is unresolved (Fig. 1A). Another unstructured loop, the L2 loop, is also found at the extremity of the stalk and in the vicinity of the L4 loop (Fig. 1A). In addition to these domains, HsMX1 and all MX proteins possess an amino-terminal (N-terminal) extension of unknown structure, which highly differs in length and sequence (Fig. 1A, 1B, Sup. Fig. 1). HsMX1 has been shown to homodimerize through the stalk domain and further oligomerizes through numerous other interfaces on the stalk and the GTPase domains (19, 20, 23). This may suggest the possible cohabitation of different types of HsMX1 oligomers within the cell. Of note, MmMx1 possesses a nuclear localization signal (NLS) localized in the third helix of the BSE (24) allowing transport into the nucleus as opposed to HsMX1 and MmMx2, which are uniquely cytoplasmic.

**Figure 1.**
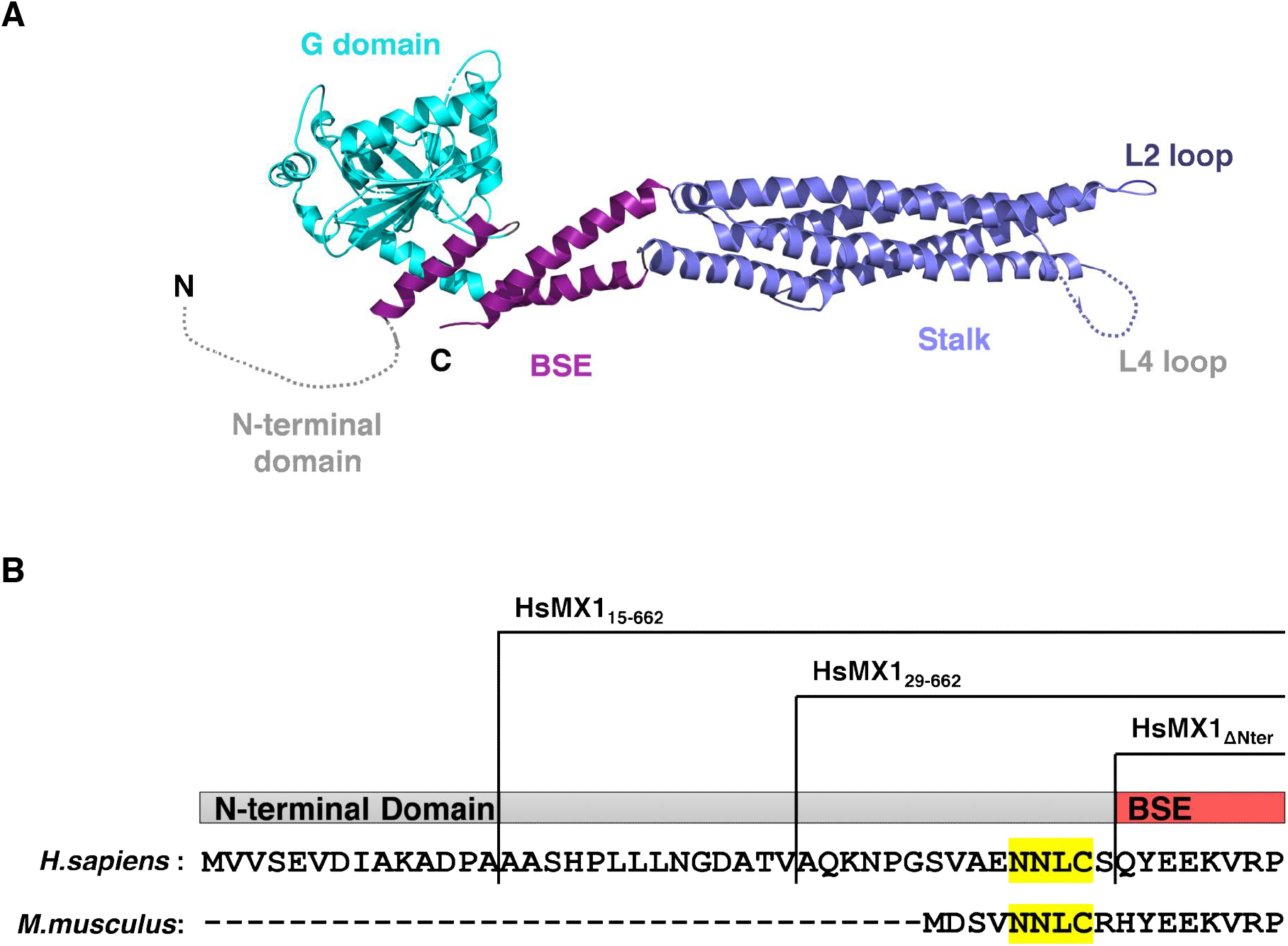
**A**. Crystal structure of HsMX1 from (20); PDB: 3SZR. The N-terminal domain and L4 loop (grey dotted lines) have unknown structures. **B**. Alignment of N-terminal domains of MX1 proteins of *Homo sapiens* (Human; P20591) and *Mus musculus* (Mouse; Q3UD61). HsMX1 N-terminal domain truncation mutations are represented and the conserved NNLC motif is highlighted in yellow.

The detailed antiviral mechanism(s) of action of MX1 proteins remain largely misunderstood, although certain intrinsic antiviral determinants have been well characterized. Four essential determinants have been identified to this day, the first being the binding/hydrolysis of GTP by the GTPase domain (25). Interestingly, HsMX2 does not require a functional GTPase domain for HIV-1 inhibition (8, 9, 26) and in the case of HsMX1, this might also be true for Hepatitis B Virus (HBV) inhibition (27). The second determinant is the presence of an intact BSE (28), the third being the possibility to oligomerize via the stalk domain (19), and the fourth requirement involves the extremity of the stalk, loops L2 (29–31) and L4 (32–34). Indeed, deletions or point mutations of the L4 loop of HsMX1 and MmMx1 abrogated antiviral activity against IAV and Thogoto virus (THOV) (32–34) and this loop was elegantly shown to have been under positive selection during evolution (35). Taken together, these studies showed that a number of intrinsic elements are needed for MX1 protein antiviral activity, however, to our knowledge, no study has so far addressed the importance of the N-terminal domain of MX1 proteins. The N-terminal domain of HsMX2, however, which is longer than that of HsMX1 (91 amino acids compared to 43 amino acids) has been shown to be a crucial determinant for HIV-1 restriction (26, 36, 37). Transferring the N-terminal domain of HsMX2 onto HsMX1 resulted in a chimeric protein able to inhibit HIV-1, without losing the anti-IAV activity (26). These data prompt that the N-terminal domain of MX proteins may generally be important for antiviral activity.

In this study, we examined the role of the N-terminal domain of several MX1 proteins, showing that this domain is essential for antiviral activity. Indeed, deletion of the N-terminal region abrogates antiviral activity against IAV and this effect was mapped to a single essential residue, leucine 41 (L41) in HsMX1. This residue was also essential for inhibition of rhabdoviruses and bunyaviruses. We further show that this residue is highly conserved between MX1 proteins of different origins, and we show that the corresponding leucines in MmMx1 (leucine 7, L7) and little yellow-shouldered bat, *Sturnira lilium*, MX1 (SlMX1) (leucine 39, L39) are also essential for anti-IAV activity. Finally, we demonstrate that mutation of this highly conserved leucine did not seem to impact lower- or higher-order oligomerization status of HsMX1 or MmMx1 in cells, the propensity to hydrolyse GTP *in vitro*, or the natural structure of the first BSE helix *in silico*. However, we show that this residue is essential for correct subcellular localization of MX1 proteins. This study therefore confirms the complex multimodular nature of MX1 proteins and defines their N-terminal region as another important antiviral module governed by an essential and conserved leucine.

## Materials and methods

### Plasmid constructs

FLAG-tagged HsMX1 and MmMx1 pCAGGS constructs were previously described (26). The *Sturnira lilium* MX1 encoding plasmid was a gift from Prof Georg Kochs (38). All FLAG-tagged HsMX1, MmMx1 and SlMX1 mutants were generated by overlapping PCR. Coding DNA sequences (CDS) of interest were subcloned into pCAGGS using NotI and XhoI cloning sites. The pRRL.sin.cPPT.SFFV/IRES-puro.WPRE lentiviral vector system has been described previously (39). All lentiviral vectors used here were obtained by subcloning of the CDSs of interest using NotI and either XhoI or SalI cloning sites. The pcDNA3-RFP-LACV-N construct was a gift from Prof Georg Kochs and generated by inserting the cDNA for mRFP1 (Addgene #14435) into pcDNA3 flanked by BamHI/NheI sites and then the LACV-N cDNA in frame with NheI/NotI. Bunyamwera N coding sequence (sequence ID: D00353.1) was synthetized (Eurofins) and cloned into pcDNA3-RFP in place of LACV-N, between NheI and NotI restriction sites, to generate pcDNA3-RFP-BUNV-N. pCAGGS-Renilla, pHSPOM1-Firefly, and pRRL.sin.cPPT.SFFV/FLAG-E2Crimson-IRES-puro have previously been described (26).

The FLAG-HsMX1, FLAG-HsMX1_T103A_, FLAG-HsMX1_L41A_, FLAG-MmMx1, FLAG-MmMx1_T69A_ and FLAG-MmMx1_L7A_ CDS were PCR-amplified from the aforementioned pRRL.sin.cPPT.SFFV/IRES-puro.WPRE lentiviral vector plasmids and cloned into pET-30 Ek/LIC expression vectors (Novagen) using KpnI and XhoI cloning sites. The different inserts are in frame with the S- and His-tags and a sequence encoding the Tobacco Etch Virus (TEV) protease cleavage site was inserted upstream of the FLAG-HsMX1, FLAG-MmMx1 and FLAG-mutant coding regions to enable tag removal during the protein purification process.

### Cells, cell culture, transduction and transfection

A549 (human lung adenocarcinoma; ATCC CCL-185), Human Embryonic Kidney 293T (HEK293T; ATCC CRL-3216) and Madin-Darby canine kidney (MDCK; ATCC CCL-34), were grown in Dulbecco’s modified Eagle medium (DMEM) supplemented with 10% fetal bovine serum (FBS) and 1% penicillin/streptomycin. Lentiviral vector (LV) stocks were produced as described previously (39). Transduction with LVs was performed by incubating cells for 8h prior to changing media for fresh media. Puromycin selection was performed 36h post-transduction and cells were used for experiments once confluency was attained. Transfection experiments were performed using Lipofectamine 2000 (Thermofisher Scientific) according to the manufacturer’s instructions.

### Immunoblotting analysis

Cells were washed in phosphate buffered saline (PBS) 1X and frozen dry at -80°C or lysed directly in lysis buffer [10 mM Tris-HCl pH7.6, 150 mM NaCl,1% Triton X100, 1 mM EDTA, 0,1% deoxycholate, 2% SDS, 5% Glycerol, 100 mM DTT, 0,02% bromophenol blue] and boiled. The lysates were resolved by SDS-PAGE and analysed by immunoblotting. Incubation with a primary Flag (mouse monoclonal M2, Sigma-Aldrich) antibody coupled to horseradish peroxidase (HRP), or Actin (Sigma-Aldrich) antibody followed by an HRP-conjugated secondary antibody was performed. Bioluminescence was then measured (Clarity ECL Western Blotting Substrate, Bio-Rad) using a ChemiDoc system (Bio-Rad).

### Immunofluorescence and Airyscan confocal microscopy

A549 cells stably expressing the FLAG-tagged MX1 proteins or mutants of interest were plated into 24-well plates on coverslips. The next day, cells were fixed using 2% paraformaldehyde (PFA) in PBS 1X for 10 minutes at room temperature (RT). Cells were permeabilized with 0.2% Triton X-100 in PBS 1X for 10 minutes followed by quenching and blocking in NGB buffer [50 mM NH_4_Cl, 2% Goat serum, 2% Bovine serum albumin, in PBS 1X] for 1h. Cells were incubated with primary antibodies for 1h, washed, then incubated for 1h with an Alexa Fluor-conjugated secondary antibody (Thermofisher). Cells were then incubated with Hoechst-33258 (Sigma-Aldrich) for 5 minutes and mounted on glass slides using ProLong™ Gold Antifade Mountant (Thermofisher). Slides were imaged on an LSM880 confocal microscope with an Airyscan module (Zeiss) using a 63x objective. Processing of the raw Airyscan images was performed on the ZEN Black software. Post-processing was performed using the FIJI software (40).

### Quantification of RFP-LACV/BUNV-N HsMX1 aggregates

HEK293T cells were plated in 96-well plates and co-transfected with of constructs of interest. 24h later, images of the transfected cells were acquired using an EVOS XL Core microscope (Thermofisher) and quantification of aggregated versus cytoplasmic RFP-LACV/BUNV-N phenotypes was performed manually on FIJI. Results were analysed using GraphPad PRISM. For representative images, HEK293T cells were plated in 24-well plates on coverslips pre-coated with Poly-L-lysine (Sigma-Aldrich) and co-transfected with constructs of interest. Cells were fixed with 2% PFA for 10 minutes, incubated with Hoechst-33258 and mounted as above, and images were acquired on an LSM880 confocal microscope with Airyscan module.

### VSV production and infection

G-pseudotyped VSVΔG-GFP-Firefly Luciferase virus was a gift from Dr Gert Zimmer and was produced as described in (41). 3.10^4^ HEK293T cells were plated in 96-well plates and co-transfected with 0.04 µg of constructs of interest together with 0.015 µg of a Renilla luciferase-encoding pCAGGS plasmid (pCAGGS-Renilla). The following day, G-pseudotyped VSVΔG-GFP-Firefly Luciferase virus stock was diluted in complete DMEM and cells were infected at a multiplicity of infection (MOI) of 0,5. 16h post-infection, cells were frozen dry at -80°C for 30 minutes and then lysed in Passive Lysis Buffer (Promega) for 30 minutes. Bioluminescence was measured using the Dual-Luciferase® Reporter Assay System (Promega) on a microplate reader (Tecan Infinite Lumi). Firefly signals were normalized by Renilla luciferase levels and results were analysed using GraphPad PRISM.

### Influenza virus production and infection

A/Victoria/3/75 (H3N2) containing the Nanoluciferase coding sequence in the PA segment (IAV-NLuc reporter virus) was described before (39). A/Victoria/3/75 (H3N2) wild-type (IAV-WT) and IAV-NLuc were produced as described previously (8). Briefly, the 8 Pol I plasmids (0.5 μg each) and 4 rescue plasmids (PB1, PB2, PA (0.32 μg each) and NP (0.64 μg)) were co-transfected into HEK293T cells in 6-well plates using Lipofectamine 3000 (Thermo Scientific). After 24h, the cells were detached and co-cultured with MDCK cells in 25 mL flasks. After 8h of co-culture in 10% serum, the medium was replaced with serum-free medium containing 0.5 μg/mL of TPCK-treated trypsin. Supernatants from day 5 post-transfection were used for virus amplification on MDCK cells. For IAV-NLuc reporter virus assays, A549 or HEK293T cells stably expressing the constructs of interest were infected at MOI 0.1 for 16h. Levels of infection were measured as described above using the Nano-Glo® Luciferase Assay System (Promega).

For the influenza minigenome infection reporter assay, 3.10^4^ HEK293T cells were co-transfected in 96-well plates with 0.015 µg of pCAGGS-Renilla and 0.04 µg of pCAGGS expressing FLAG-tagged MX1, mutants or controls and 0.03 µg of the minigenome plasmid pHSPOM1-Firefly. The negative sense, Firefly-coding minigenome transcribed from pHSPOM1-Firefly by the cellular RNA Pol I contains mutations 3-5-8 in the 3’ end of the promoter to increase replication and transcription efficiency following IAV infection (42). 24h later, the cells were infected with wild-type A/Victoria/3/75 (H3N2) at MOI 0.1. 16h post infection, cells were frozen dry at -80°C for 30 minutes and lysed in Passive Lysis Buffer (Promega). Firefly and Renilla activities were then measured using the Dual-Luciferase® Reporter Assay System (Promega). Firefly signals were normalized by Renilla luciferase levels. Results were analysed using GraphPad PRISM.

### Cross-link experiments

Crosslinking experiments were performed as described in (43). Briefly, cells expressing the FLAG-tagged MX1 constructs of interest were lysed in 0.5% Triton X-100-PBS 1X buffer in the presence of complete protease inhibitor cocktail (Roche) for 20 minutes on ice. Cells were water bath sonicated and centrifuged at 1,500 x *g* for 10 minutes at 4°C. Disuccinimidyl suberate (DSS, ThermoFisher) was added to the cells at a final concentration of 100 µg/ml (or an equivalent volume of Dimethyl sulfoxide (DMSO) was added to the control conditions) and incubated for 1h at RT. Laemmli was added at 1X final and samples were resolved by SDS-PAGE without boiling on a 5% acrylamide gel. Immunoblotting was performed as mentioned above.

### Protein expression and purification

The recombinant plasmids pET-30 Ek/LIC expressing FLAG-HsMX1, FLAG-MmMx1 and FLAG-mutants were transformed in an *Escherichia coli* BL21 (DE3) strain resistant to Phage T1 (New England Biolabs) carrying pRARE2. One colony was used to inoculate an overnight culture of 125mL Lysogeny broth (LB) medium supplemented with kanamycin (50 μg/mL) and chloramphenicol (30 μg/mL). This culture was diluted in 2.5L of LB medium supplemented with the two antibiotics. The cells were grown at 16°C to an optical density at 600 nm of 0.8, then protein expression was induced with 1 mM Isopropyl-β-D-thiogalactoside (IPTG) and the culture was grown overnight at 16°C. The cells were harvested by centrifugation at 8 200*g* for 20 minutes and resuspended in 30 mL of buffer A (50 mM Tris–HCl pH 8, 400 mM NaCl, 5 mM MgCl_2_, 7 mM β-mercaptoethanol, 40 mM imidazole, 10% glycerol and 1 mM benzamidine). The cells were disrupted by sonication and cell debris were removed by centrifugation at 28 000*g* for 60 minutes. The supernatant was loaded at 4°C on Ni–NTA agarose beads previously equilibrated with buffer A. The beads were washed once with buffer A and twice with buffer B (50 mM Tris–HCl pH 8, 1 M NaCl, 5 mM MgCl_2_, 7 mM β-mercaptoethanol, 40 mM imidazole and 10% glycerol) and elution was performed with buffer E (50 mM Tris–HCl pH 8, 200 mM NaCl, 5 mM MgCl_2_, 7 mM β-mercaptoethanol, 500 mM imidazole, 10% glycerol). The eluted protein was incubated with His-tagged TEV protease purified in our laboratory in a 1:100 (w:w) ratio; the cleavage reaction was performed during dialysis (dialysis-bag cutoff 12–15 kDa) against 1L dialysis buffer D (50 mM Tris–HCl pH 8, 200 mM NaCl, 5 mM MgCl_2_, 5 mM β-mercaptoethanol) overnight at 4°C. After dialysis, the proteins were centrifuged for 20 minutes at 28 000*g* and the supernatant was loaded again at 4°C on Ni–NTA agarose beads equilibrated with buffer D. The proteins without the His-tag were collected in the flow-through, concentrated to 5 mg/mL using a Vivaspin® column (50 kDa cutoff, Sartorius), loaded onto a size-exclusion chromatography column (Superose 6 Increase 10/300 GL, GE Healthcare) and eluted with buffer F (50 mM Tris–HCl pH 8, 500 mM NaCl, 5 mM MgCl_2_, 0,5 mM β-mercaptoethanol, 10% glycerol). Aliquots of purified FLAG-tagged proteins were snap frozen in liquid nitrogen and stored at -80 °C.

### GTPase activity assays

Recombinant FLAG-tagged, wild-type and mutant HsMX1 or MmMx1 proteins (0.2 mg/mL) were incubated at 37°C in GTPase assay buffer [50 mM Tris–HCl pH 8, 500 mM NaCl, 5 mM MgCl_2_, 0,5 mM β-mercaptoethanol, 10% glycerol, 50 µM GTP, 26 nM [α-^32^P]-GTP], as described previously (26). The reactions were stopped at the indicated time by addition of [2 mM EDTA, 0,5% SDS]. The reaction products were resolved by thin-layer chromatography (TLC, Merck Millipore) using TLC buffer [1 M LiCl, 1 M formic acid] and detected using phosphor screen autoradiography (Amersham Typhoon apparatus).

### Molecular modelling and dynamic simulations of wild-type HsMX1 and HsMX1_L41A_

HsMX1 WT (including the complete N-terminal domain and L4 loop) as well as HsMX1_L41A_ were modelled using AlphaFold (44) through the ColabFold (45) platform. Predicted 3D models were compared to crystal (PDB: 3SZR) or low-resolution EM (PDB: 3ZYS) structures for examination of the overall folding of the three main domains (GTPase, BSE and stalk) and also compared to 3D models based on these templates and built with Modeller v9.19 (46). Structural alignments of AlphaFold models and X-ray structure gave RMSD values (for all backbone atoms) of 0.4, 0.2 and 1.4 Å for GTPase, BSE and Stalk domains, respectively. Molecular dynamics (MD) simulations were performed using NAMD3 (47) and CHARMM36m force field (48). Briefly, each protein was immersed in an explicit solvent box (TIP3P water model) using 10 Å of edge in each direction (in respect to protein dimensions), then neutralized with NaCl ions at a physiological concentration (0.154 M) and energy minimized for 50 ps using the conjugate gradients method. After a gradual heating from -273°C to 37°C, each system was further equilibrated for 300 ps using periodic boundary conditions to replicate the system in each direction. A trajectory of 500 ns was then produced (1 frame saved every 20 ps) for analysis in the isobaric-isothermal ensemble to keep constant temperature (37°C) and pression (1 atm) using Langevin dynamics and Langevin piston methods. The Newton’s equation of motions was integrated using a timestep of 2 fs using the r-RESPA algorithm (49) with the short-range Lennard-Jones potential smoothly truncated from 10 to 12 Å and the PME (Particle Mesh Ewald) approach (50) used for calculating long-range electrostatics with a grid spacing of 1 Å. Trajectories analysis was performed using VMD (51), its Timeline plugin for per-residue Phi and Psi dihedral angles calculation, 3D-graphs (dihedral angles plots) were made with GnuPlot-5.4 and MD snapshots were illustrated with the PyMol Molecular Graphics System (v2.5, Schrödinger, LLC).

### Data availability statement

Datasets are currently being deposited on a public repository and the URL will be provided.

### Statistical analyses

Statistical analyses were performed with the GraphPad Prism software. The analysis types performed are indicated in the figure legends. Comparisons are relative to the indicated condition. P-values are indicated as the following: ns = not significant, p<0.05 = *, p<0.01 = **, p<0.001 = *** and p<0.0001 = ****.

## Results

### Leucine 41 from the N-terminal region of human MX1 is essential for antiviral activity against IAV

The loop L4 and all 43 amino acids of the N-terminal domain are absent from the published crystal structures of HsMX1 (19, 20) (Fig. 1A). While the L4 loop has been extensively studied in the past (33, 35), to date, the importance of the N-terminal region for the antiviral activity of HsMX1 has not been evaluated. Therefore, to address this, we generated a series of N-terminal truncation mutants for HsMX1, with mutants missing either the first 14 (HsMX1_15-662_) or 28 (HsMX1_29-662_) amino acids, or the entire N-terminal region (HsMX1_44-662_, named hereafter HsMX1_ΔNter_) (Fig. 1B). In parallel to two negative controls (E2 Crimson fluorescent protein, termed CTRL, and HsMX1 inactive GTPase mutant, HsMX1_T103A_), the wild-type (WT) protein and mutants were ectopically expressed in HEK293T cells, and an IAV minigenome infection reporter assay was performed, as reported previously (8). In this assay, a negative sense minigenome coding for the Firefly luciferase is recognized, replicated and transcribed by IAV polymerase in IAV infected cells. Thus, Firefly activity is used to monitor replication efficiency. Cells were also co-transfected with a Renilla luciferase coding plasmid for normalization. We observed that the deletion of the first 14 or 28 amino acids had no effect on HsMX1 antiviral activity (Fig. 2A). In contrast, the deletion of the entire N-terminal region totally abrogated antiviral activity in a comparable manner to the control inactive GTPase mutant HsMX1_T103A_ (Fig. 2A). This suggested the presence of essential residues located between positions 29 and 43 of the N-terminal domain of HsMX1. *Mus musculus* Mx1 (MmMx1) is another well studied IAV-inhibiting MX1 protein. Alignment of MmMx1 and HsMX1 proteins revealed a conserved motif of four amino acids located between positions 29 and 43 of HsMX1: 39-NNLC-42 (or 5-NNLC-8 in the case of MmMx1) (Fig. 1B). Upon replacement in HsMX1 of these four amino acids with four alanines (HsMX1_NNLC39-42A_), we saw a complete loss of antiviral activity comparable to that of HsMX1_ΔNter_ and HsMX1_T103A_ (Fig. 2A, top panel). Alanine point mutations of these four residues (HsMX1_N39A_, HsMX1_N40A_, HsMX1_L41A_ and HsMX1_C42A_), revealed that leucine 41 (L41) was responsible for the loss of function phenotype observed with the MX1_NNLC39-42A_ mutant, whereas the other three mutants completely retained their anti-IAV activity (Fig. 2A, top panel). While the HsMX1_29-662_ and HsMX1_ΔNter_ truncation mutants showed decreased expression levels compared to the WT protein, as assessed by immunoblotting, the loss of antiviral activity of the HsMX1_NNLC39-42A_ and HsMX1_L41A_ mutants could not be attributed to a decrease in expression levels (Fig. 2A, bottom panel).

**Figure 2.**
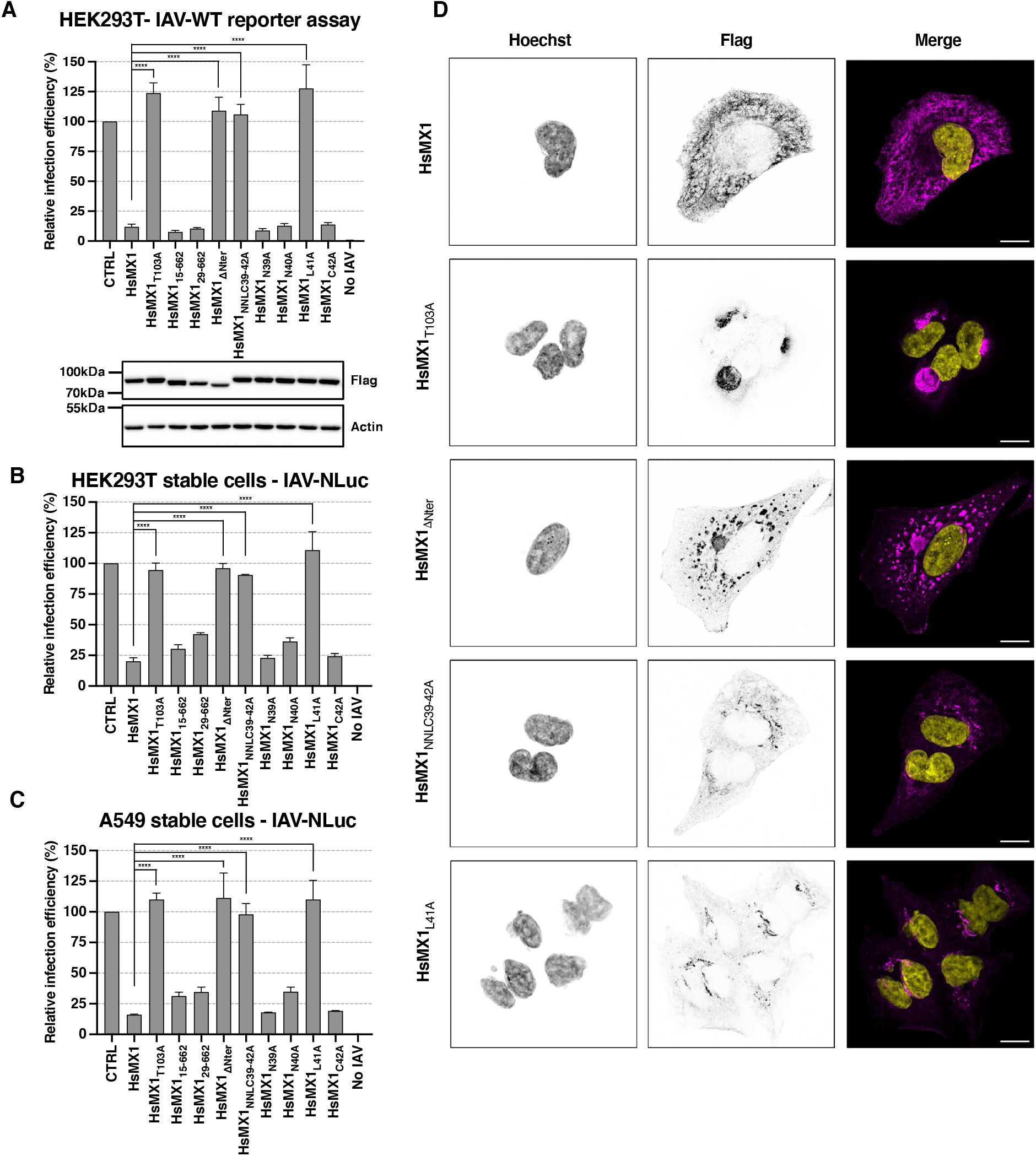
The N-terminal domain of HsMX1 is essential for IAV restriction and subcellular localization. **A**. Relative infection efficiency using an influenza minigenome infection reporter assay in transiently transfected HEK293T cells with FLAG-tagged WT HsMX1, mutants or E2 Crimson control (CTRL) (top), with a representative immunoblot (bottom; Actin served as a loading control). **B**. Relative infection efficiency of IAV-NLuc in HEK293T cells stably expressing FLAG-tagged proteins of interest. **C**. Relative infection efficiency of IAV-NLuc in A549 cells stably expressing the FLAG-tagged proteins of interest. **D**. Representative Airyscan immunofluorescence images of A549 cells stably expressing the FLAG-tagged proteins of interest, stained for anti-FLAG (magenta) and nuclei (yellow). Single channels are shown in inverted grey; scale bar: 10µm. For the infection assays, results were normalized to 100% for the CTRL condition, and the mean and standard deviations of 3 independent experiments are shown. Ordinary One-way ANOVA multiple comparison with HsMX1 was performed; **** = p<0.0001

The observed phenotypes were then confirmed using a Nanoluciferase reporter-expressing version of A/Victoria/3/75 (IAV-NLuc) (39) as a second infection readout, both in HEK293T (Fig. 2B) and in lung-derived A549 cells (Fig. 2C), which stably expressed the control and mutant proteins. Similarly to what was observed in the minigenome infection reporter assay (Fig. 2A), HsMX1_ΔNter_, HsMX1_NNLC39-42A_ and HsMX1_L41A_ were completely inactive against IAV, comparable to the inactive HsMX1_T103A_ mutant, whereas the other mutants retained antiviral activity (Fig. 2B-C).

Next, we investigated the subcellular localization of the N-terminal HsMX1 mutants using super-resolution Airyscan microscopy. As reported previously, HsMX1 showed a honeycomb-like, punctate cytoplasmic staining and HsMX1_T103A_ presented a juxtanuclear accumulation (Fig. 2D) (26, 52, 53). In contrast, the HsMX1_ΔNter_ mutant formed cytoplasmic aggregate-like structures that varied in size and were dispersed throughout the cytoplasm (Fig. 2D). Unlike this mutant, HsMX1_NNLC39-42A_ and HsMX1_L41A_ accumulated at the perinuclear region into small spherical structures that were phenotypically different from those of HsMX1_T103A_ (Fig. 2D). This attests to a potential difference between the L41 point and N-terminal truncation mutants, which might be linked to differences in protein synthesis or stability as seen by immunoblot (Fig. 2A).

Taken together, this data showed that the N-terminal region, and more precisely, leucine 41, was essential for anti-IAV activity and correct subcellular localization of HsMX1.

### L41 is essential for HsMX1 restriction of other families of RNA viruses

To further understand the importance of this newly discovered essential residue for the antiviral activity of HsMX1, we tested the restriction abilities of aforementioned HsMX1 mutants against other RNA viruses known to be inhibited by HsMX1. HEK293T cells were co-transfected with either a control or HsMX1 WT or mutant expression constructs along with a Renilla luciferase-coding plasmid and the infection levels of Firefly-expressing, Vesicular Stomatitis Virus (VSV) rhabdovirus were measured (Fig. 3A). Reminiscent of what observed for IAV (Fig. 2A), the HsMX1_ΔNter_, HsMX1_NNLC39-42A_ and HsMX1_L41A_ mutants totally lost their ability to restrict VSV (Fig. 3A). This therefore demonstrates the essential role of L41 for the restriction of VSV by HsMX1.

**Figure 3.**
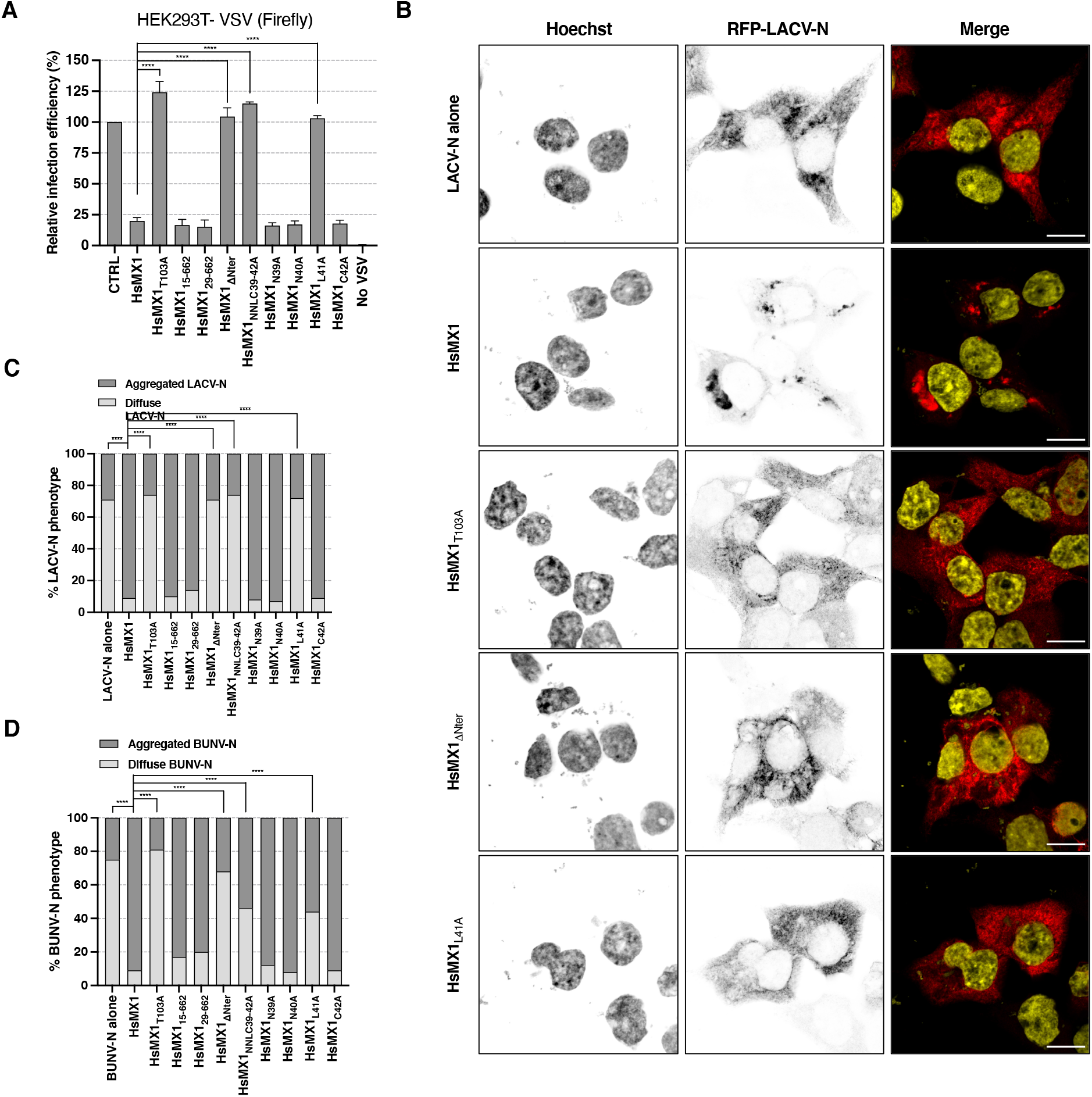
L41 is essential for restriction of other RNA viruses. **A**. Relative VSV-Firefly infection efficiency of HEK293T cells transfected with the indicated FLAG-tagged HsMX1 constructs or E2-Crimson (CTRL). **B**. Representative Airyscan images of the phenotype of RFP-LACV-N in transfected HEK293T cells in the presence of different FLAG-tagged HsMX1 constructs. Single channels are shown in inverted grey and on the merge images, RFP-LACV-N is shown in red and nuclei in yellow. Scale bar: 10µm. **C**. Quantification of the percentage of RFP-LACV-N and WT or mutant FLAG-tagged HsMX1 transfected HEK293T cells presenting a diffuse or aggregated RFP-LACV-N staining. **D**. Similar to C, with RFP-BUNV-N. The graphs show the mean and standard deviations of 3 (**A, C**) or 2 (**D**) independent replicates. A total of 1500-2000 cells (**C**) and 950-1250 cells (**D**) were counted for each condition, respectively. Ordinary One-way ANOVA multiple comparison with HsMX1 was performed for diffuse LACV-N or BUNV-N values; **** = p<0.0001

As HsMX1 is well-known to inhibit the Orthobunyaviruses La Crosse Virus (LACV) and Bunyamwera Virus (BUNV) (54), we tested the importance of the N-terminal domain against these viruses. The inhibition phenotype of HsMX1 towards LACV and BUNV is characterized by the aggregation of their Nucleoprotein (N) into perinuclear aggregates (54), a phenotype that can be recapitulated by co-expressing HsMX1 and the N protein in cells and easily visualised by using a fusion of N with the red fluorescent protein (RFP) (54). Hence, HEK293T cells were co-transfected with plasmids expressing HsMX1 or the mutants of interest, together with an RFP-LACV-N expression plasmid. As reported previously (54), HsMX1 induced LACV-N aggregates and this ability was lost by HsMX1_T103A_, as shown on the representative images (Fig. 3B). To quantify HsMX1 protein activity here, the percentage of cells showing a diffuse or aggregated RFP-LACV-N staining was determined. In control conditions (RFP-LACV-N alone), around 70% of the cells displayed a diffuse cytoplasmic pattern (Fig. 3C), with 30% of cells showing limited aggregation, possibly due to overexpression levels. In the presence of HsMX1, only around 10% of cells showed a diffuse cytoplasmic staining, with around 90% of cells showing an RFP-LACV-N aggregation at the perinuclear region (Fig. 3C, 3B). In the presence of HsMX1_ΔNter_, HsMX1_NNLC39-42A_ and HsMX1_L41A_, there was a complete loss of RFP-LACV-N aggregation phenotype, with distributions comparable to the RFP-LACV-N alone or HsMX1_T103A_ conditions (Fig. 3C, 3B). The same experiment was performed for the RFP-tagged Nucleoprotein of BUNV (RFP-BUNV-N) (Fig. 3D). Similar results to the ones obtained with LACV-N were observed, although with a somewhat reduced, but nevertheless significant, impact of HsMX1_L41A_ and HsMX1_NNLC39-42A_ mutations on the ability of HsMX1 to induce RFP-BUNV-N aggregation. These data show that the N-terminal domain, and in particular L41, is important for LACV and BUNV restriction as well as for VSV and IAV.

### Conserved importance of the N-terminal domain for the anti-IAV activity of MX1 proteins of other mammals

MX1 proteins from various mammalian species are known to harbour anti-IAV activity (7, 55). Therefore, we sought to explore the importance of the conserved leucine we identified in the N-terminal region in the anti-IAV activity of MX1 proteins from other mammals. Mouse Mx1 (MmMx1) strongly inhibits IAV, and, interestingly, the N-terminal domain of MmMx1 is highly different to that of HsMX1, consisting of only 9 amino acids versus 43 for HsMX1 (Fig. 1B). Nevertheless, MmMx1 contains a leucine at position 7, placed exactly 3 amino acids before the start of the first α-helix of the BSE domain, similarly to leucine 41 of HsMX1 (Fig. 1B). To determine whether the N-terminal domain, and notably L7, were also essential for MmMx1 anti-IAV activity, we performed a truncation of the N-terminal domain (MmMx1_ΔNter_) and an alanine point mutation of L7 (MmMx1_L7A_). We stably expressed these constructs, as well as a GTPase inactive mutant (MmMx1_T69A_), in A549 cells and infected them with the IAV-NLuc reporter virus (Fig. 4A, top panel). Whereas MmMx1 potently inhibited IAV replication, MmMx1_ΔNter_ and MmMx1_L7A_ showed no antiviral activity, similar to the GTPase inactive mutant MmMx1_T69A_. It is noteworthy that, similarly to HsMX1_ΔNter_ (Fig 2A), MmMx1_ΔNter_ had reduced expression levels (Fig. 4A, bottom panel). However, MmMx1_L7A_ showed similar expression levels as compared to WT MmMx1 (Fig. 4A, bottom panel). Using super-resolution Airyscan microscopy, WT MmMx1 was located into classical nuclear bodies as well as being present to a lesser extent in the cytoplasm (Fig. 4B). MmMx1_ΔNter_ was localized in nuclear rod-like structures, and MmMx1_L7A_ was found as nuclear rod-like structures that associated into star-shaped superstructures, as well as in perinuclear, aggresome-like structures (Fig. 4B). Taking all these data together, we show that the N-terminal domain, and more specifically leucine 7, is essential for MmMx1 anti-IAV activity and subcellular localization.

**Figure 4.**
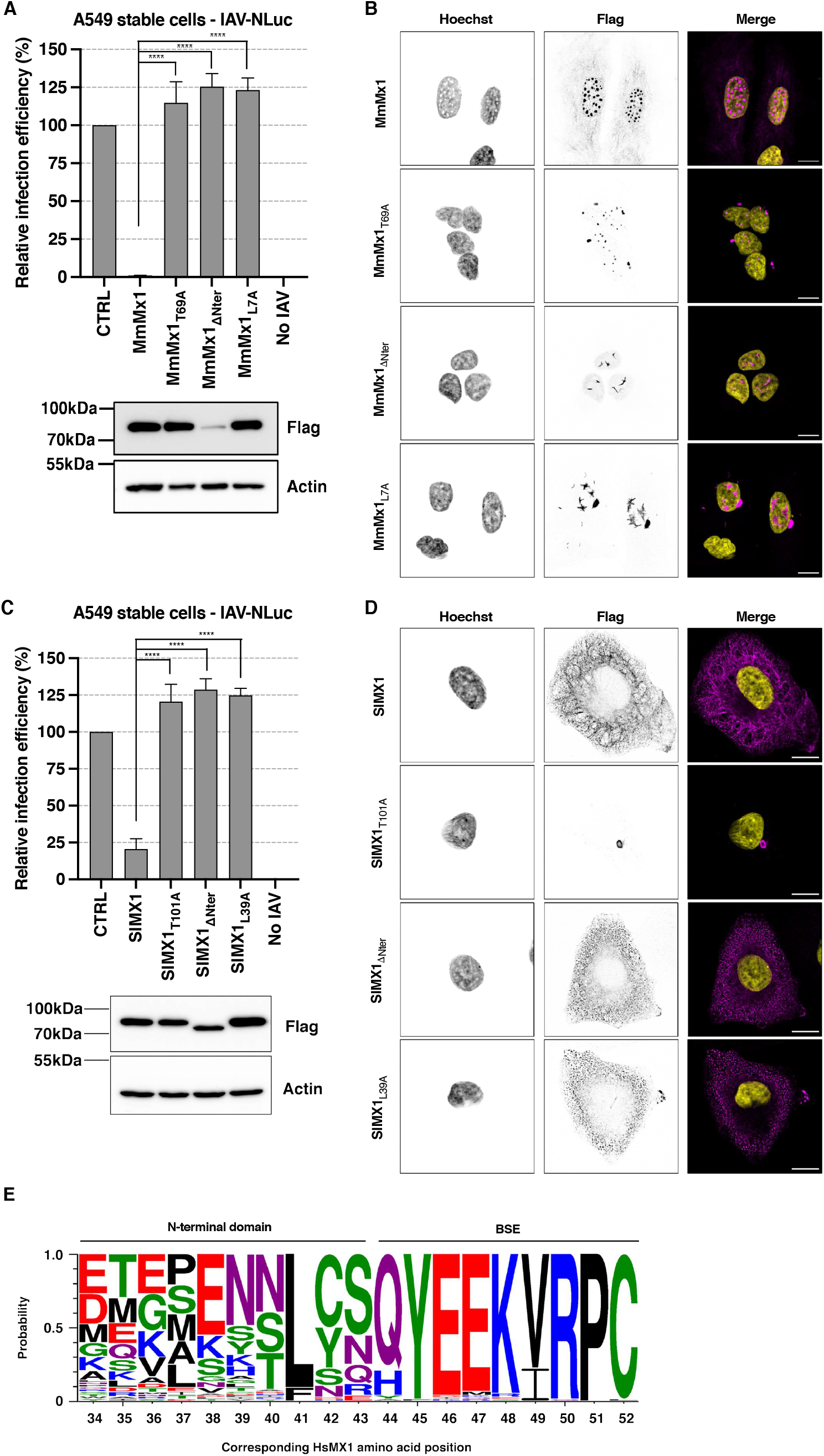
Importance of the conserved leucine of the N-terminal domains of MX1 proteins of other species. **A**. Top panel, relative IAV-NLuc infection efficiency of A549 cells stably expressing FLAG-tagged E2 Crimson (CTRL), WT or mutant MmMx1 proteins. Bottom panel, a representative immunoblot is shown. **B**. Representative Airyscan images of A549 cells stably expressing FLAG-tagged MmMx1 or mutants. Single channels are shown in inverted grey, FLAG proteins in magenta and nuclei in yellow; scale bar: 10µm. **C**. Top panel, relative IAV-NLuc infection efficiency of stable A549 cells stably expressing FLAG-tagged Renilla luciferase (CTRL), SlMX1 or mutants. Bottom panel, a representative immunoblot is shown. **D**. Representative Airyscan images of A549 cells stably expressing FLAG-tagged SlMX1 or mutants, (colour code as in **B**). **E**. WebLogo3 analysis of MX1 proteins from 76 different species. Amino acids of all protein sequences aligned correspond to those of HsMX1 from position 34-52. Alignment was performed using the align tool from Uniprot (www.uniprot.org/align/). Full alignment can be found in Sup. Fig. 1. Colors represent amino acid chemistry groups. **A** and **C**. The graphs show the mean and standard deviation of 3 independent experiments. Ordinary One-way ANOVA multiple comparison with WT MmMx1 or SlMX1 was performed; **** = p<0.0001

We next decided to look at a more distant relative of humans and therefore chose to study a bat MX1 protein, *Sturnira lilium* (little yellow-shouldered bat) MX1 (SlMX1), that also possesses anti-IAV activity (38). SlMX1 has a similar sized N-terminal domain (41 amino acids in length) to that of HsMX1, with, again, a leucine located 3 amino acids before the start of the BSE, leucine 39 (L39) (Sup. Fig. 1). As previously, we generated a truncation of the N-terminal domain (SlMX1_ΔNter_) and an alanine point mutation of L39 (SlMX1_L39A_). We produced A549 cells stably expressing WT SlMX1 and mutants and infected them with the IAV-NLuc reporter virus (Fig. 4C). As for HsMX1 and MmMx1, SlMX1_ΔNter_ and SlMX1_L39A_ mutants lost their antiviral activity against IAV, similar to the GTPase inactive mutant SlMX1_T101A_ (Fig. 4C, top panel). In opposition to HsMX1_ΔNter_ and MmMx1_ΔNter_, however, SlMX1_ΔNter_ did not show profoundly decreased expression levels and the SlMX1_L39A_ mutant seemed to show slightly higher expression levels than the WT SlMX1 (Fig. 4C, bottom panel). In terms of subcellular localization, SlMX1 was found in cytoplasmic puncta forming honeycomb-like subcellular structures, reminiscent of HsMX1 (Fig. 4D, 2D). In contrast, SlMX1_ΔNter_ and SlMX1_L39A_ were present in small puncta in the cytoplasm but lost their honeycomb-like network (Fig. 4D). This confirmed that the conserved leucine present in the N-terminal domain of mammalian MX1 proteins is a crucial residue for their antiviral activity and subcellular localization.

Considering that this leucine is conserved for human, mouse and bat MX1 proteins, we aligned the amino acid sequence of MX1 from 76 different species, comprising mammals, birds, amphibians and fish (Sup. Fig. 1). An analysis using the WebLogo3 tool (weblogo.threeplusone.com) (56, 57) of amino acids corresponding to positions 34-52 of HsMX1 from all 76 MX1 protein sequences shows the extreme conservation of the leucine corresponding to L41 throughout all MX1 proteins aligned (Fig. 4E), with the rest of the N-terminal domain before and after this residue being much less conserved than the leucine residue. Indeed, the leucine residue is conserved to a level similar to amino-acids from the BSE domain (Fig. 4E). Of note, only a few species of fish, including zebrafish (*Danio rerio*) and goldfish (*Carassius auratus*) did not show conservation of this leucine, with a phenylalanine (F) found in place of this leucine (Fig. 4E and Sup. Fig. 1).

Altogether these results show that similarly to HsMX1 L41, L7 and L39 from MmMx1 and SlMX1 respectively, are essential for antiviral activity against IAV and that this leucine is the most conserved residue of the N-terminal domain across many species.

### Loss of antiviral activity of N-terminal Leucine mutants is neither due to an oligomerization defect nor the inability to hydrolyse GTP

Correct oligomerization and GTPase activity being both essential for HsMX1 antiviral activity (19, 25), we wondered whether mutating L41 in HsMX1 could impair these properties. First, we performed crosslinking experiments on WT HsMX1, HsMX1_T103A_, HsMX1_M527D_ (a monomeric mutant (19)) and HsMX1_L41A_. HEK293T cells stably expressing these proteins showed the expected phenotypes with respect to IAV-NLuc infection, with a loss of antiviral activity for HsMX1_M527D_ comparable to that of HsMX1_T103A_ and HsMX1_L41A_ (Fig. 5A). Disuccinimidyl suberate (DSS) crosslinking followed by immunoblotting experiments showed that, contrary to WT HsMX1, HsMX1_M527D_ did not assemble into lower- or higher-order oligomers and was only detected as a monomer (Fig. 5B). In contrast, HsMX1_L41A_ showed a similar capacity to oligomerize into lower- and higher-order oligomers than WT HsMX1 or the GTPase inactive mutant HsMX1_T103A_ (Fig. 5B). We can therefore conclude that the absence of antiviral activity of this mutant is not due to an oligomerization defect. Next, the same experiments were performed with MmMx1 and mutants stably expressed in HEK293T cells (Fig. 5C, 5D). Of note, MmMx1_M493D_, the potential monomeric mutant corresponding to HsMX1_M527D_, could not be used as expression levels were undetectable (data not shown). MmMx1 stable overexpression in HEK293T cells strongly inhibited IAV replication, although somewhat less efficiently compared to stable overexpression in A549 cells (compare Fig. 5C with Fig. 4A). Both MmMx1_T69A_ and MmMx1_L7A_ mutants totally lost antiviral activity (Fig. 5C). In a similar fashion to HsMX1_L41A_, MmMx1_L7A_ was able to oligomerize into lower- and higher-order oligomers (Fig. 5D), showing that the inability of MmMx1_L7A_ to inhibit IAV was not due to an oligomerization defect.

**Figure 5.**
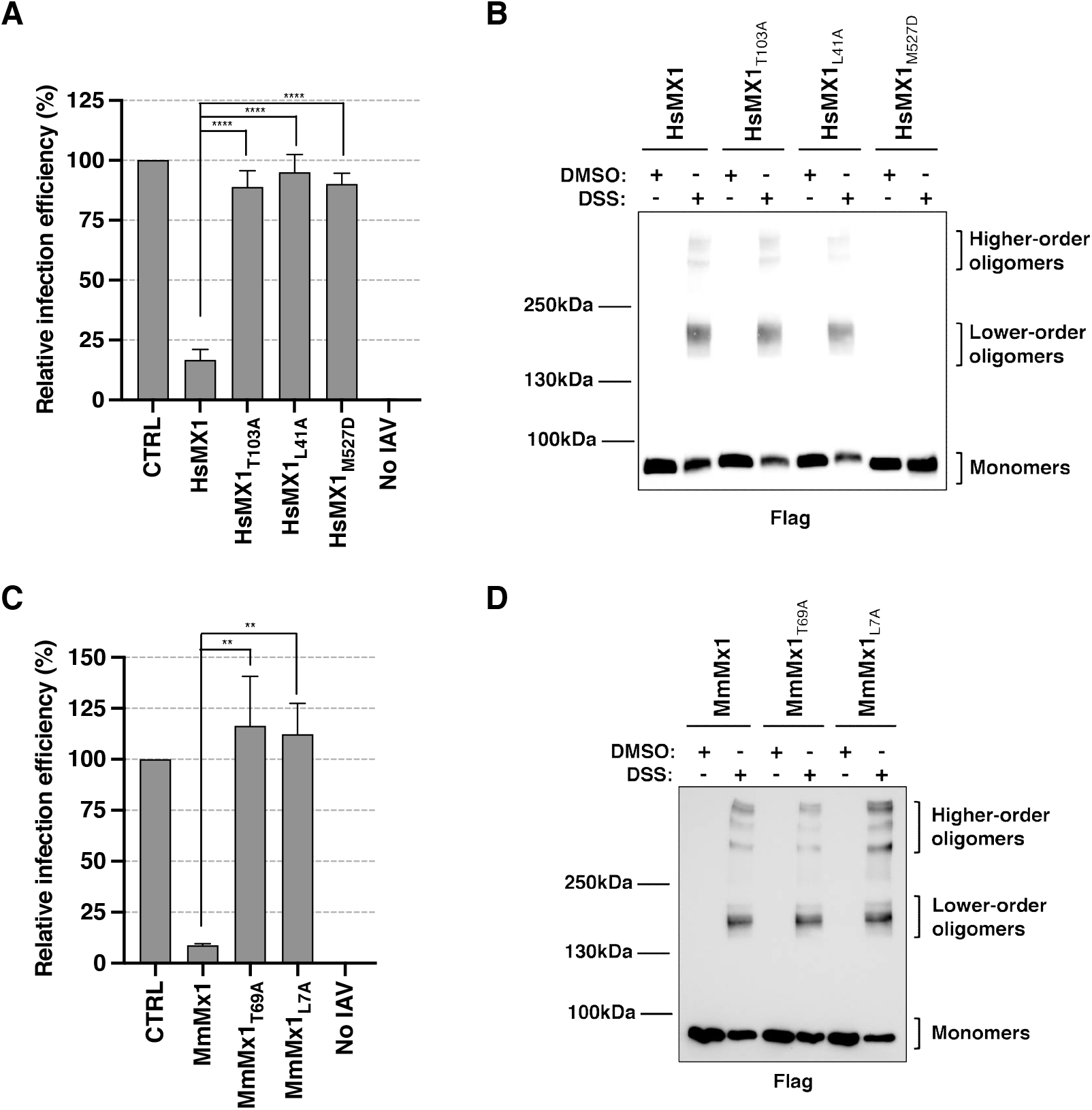
Chemical crosslinking indicates that the lack of antiviral activity of HsMX1_L41A_ and MmMx1_L7A_ is not due to an oligomerisation defect. **A, C**. HEK293T cells were stably transduced to express FLAG-tagged HsMX1/MmMx1 proteins and mutants or E2-Crimson (CTRL), challenged with IAV-NLuc and relative infection efficiency was analysed. The mean and standard deviation of 3 independent experiments is shown. Ordinary One-way ANOVA multiple comparison with WT HsMX1 or MmMx1 was performed; ** = p<0.01, **** = p<0.0001. **B, D**. The same cells were lysed on ice, sonicated and DSS (100µg/ml) (or DMSO) treated for 60 minutes. After quenching, the lysates were resolved by SDS-PAGE and an immunoblot was performed to detect the FLAG-tagged MX1 proteins. Representative immunoblots are shown.

To assess the potential impact of leucine mutations on GTPase activity, recombinant WT HsMX1, MmMx1 and mutant proteins HsMX1_T103A_, HsMX1_L41A_, MmMx1_T69A_ and MmMx1_L7A_ were produced in *E. coli* and purified (Fig. 6A). A GTPase activity assay was performed to assess the ability of these proteins to convert GTP into GDP (Fig. 6B-D). While the WT HsMX1 and the GTPase inactive mutant HsMX1_T103A_ were respectively able and unable to hydrolyse GTP, as expected (58) (Fig. 6B), the HsMX1_L41A_ mutant showed comparable GTPase activity to WT HsMX1, indicating no defect in enzymatic activity. In the case of MmMx1, WT MmMx1 could hydrolyse GTP and MmMx1_T69A_ was unable to hydrolyse it (Fig. 6D), as expected (34). However, while MmMx1_L7A_ was still able to hydrolyse GTP to some extent (Fig. 6D), in comparison to WT MmMx1, MmMx1_L7A_ showed ∼60% decrease in enzymatic activity (Fig. 6D). Time course experiments showed that GTPase activity efficiency of HsMX1_L41A_ was identical to that of WT HsMX1 regardless of the duration of the assay (Fig. 6C). In contrast, MmMx1_L7A_ appeared less active than WT MmMx1 and showed a 30% to 50% decrease in GTPase activity at the different time points (Fig. 6E). Nevertheless, these experiments showed that the loss of antiviral activity of HsMX1_L41A_ and MmMx1_L7A_ mutants could not be attributed to a complete inability to hydrolyse GTP.

**Figure 6.**
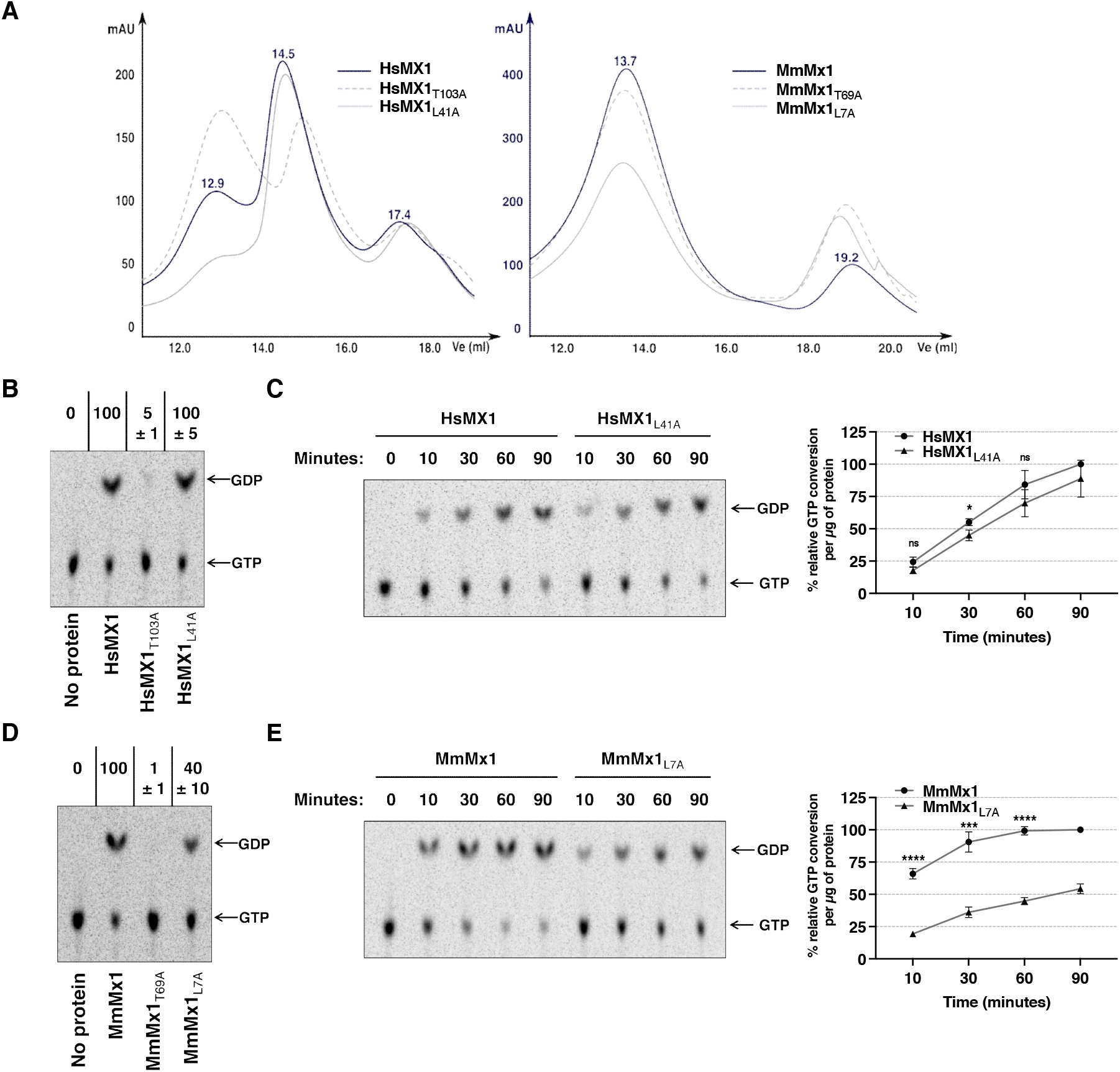
GTPase assays reveal that HsMX1_L41A_ is catalytically active and that MmMx1_L7A_ has a slightly impaired GTPase activity. **A**. Recombinant WT and mutant FLAG-HsMX1 and FLAG-MmMx1 proteins were produced in *E. coli* and purified. The elution profiles on Superose 6 increase 10/300GL columns are shown for WT FLAG-HsMX1 and FLAG-MmMx1 proteins (black lines) in parallel to the indicated FLAG-tagged mutants (grey continuous and dashed lines). **B-E**. GTPase assays were performed with the recombinant proteins and GDP and GTP levels were resolved using thin-layer chromatography (TLC) and visualized by autoradiography. **B**. Representative TLC image for a 60-minute GTPase assay with WT and mutant HsMX1 proteins. Numbers above the TLC represent the average of normalized signal intensity of the GDP spots from 3 independent experiments. **C**. Representative TLC image of a kinetic experiment performed on WT and L41 mutant HsMX1 (the GTPase assay was stopped at the indicated time points, i.e. 10, 30, 60, 90 minutes) (Left). The experiment was performed 3 times independently and the signals quantified and normalized to the 90-minute time point for the WT protein (Right). **D**. Similar to B, with WT and mutant MmMx1 proteins and a 30-minute GTPase assay. **E**. Similar to C, with WT and mutant MmMx1 proteins. **C**. and **E**. right panels, the mean and standard deviation of 3 independent experiments are shown. Paired Student t-test was performed for the same timepoints between WT and mutant proteins; ns: non-significant, *: p<0.05, ***: p<0.001, ****: p<0.0001

### Mutating L41 of HsMX1 does not seem to impact BSE structural properties

As mentioned above, the structure of the N-terminal domain of HsMX1 has not been solved (19, 20) (Fig. 1A). The AlphaFold tool (44, 59) may accurately predict structures of proteins (or regions of proteins) that have not been experimentally determined by structural approaches. Concerning HsMX1, the N-terminal domain can be modelled starting from serine 35 (S35) (alphafold.ebi.ac.uk/entry/P20591), although the confidence levels are not high and even lower for the long N-terminal unstructured tail. Nevertheless, AlphaFold predicts that the first α-helix of the BSE starts at S35, which would entail a redefinition of BSE region and would make L41 actually part of the BSE (Fig. 7A). Hence, mutation of L41 into an alanine could entail a restructuration of the BSE helix, explaining the loss of antiviral activity of the HsMX1_L41A_ mutant. However, performing molecular dynamics (MD) simulations with WT HsMX1 or the L41A mutant (Fig. 7B, 7C, Sup. Movie 1, Sup. Movie 2) showed that the structural behaviour of this α-helix did not change when comparing both proteins. Indeed, except of a hinge or a turn-like structure occurring quite rapidly around residues E46-E47 but observed in both cases, the α-helicoidal secondary structure is maintained all throughout the simulations (0.5 µs) (Fig. 7B, 7C, Sup. Movie 1, Sup. Movie 2). To confirm this point, two dihedral angles (Phi and Psi) from part of the N-terminal helix (residues ranging from 33 to 60) were measured over time leading to very similar values for both angles between both WT and the L41A mutant (Sup. Fig. 2). Angle values were characteristic of α-helix structure with average values centred at -60° and - 50° for Phi and Psi, respectively. This hinge or partial helix brake seems slightly more pronounced for WT HsMX1 than for the mutant when examining thoroughly angles values, however, this unique exception may not be sufficient to explain the difference observed in antiviral activity between the two constructs.

**Figure 7.**
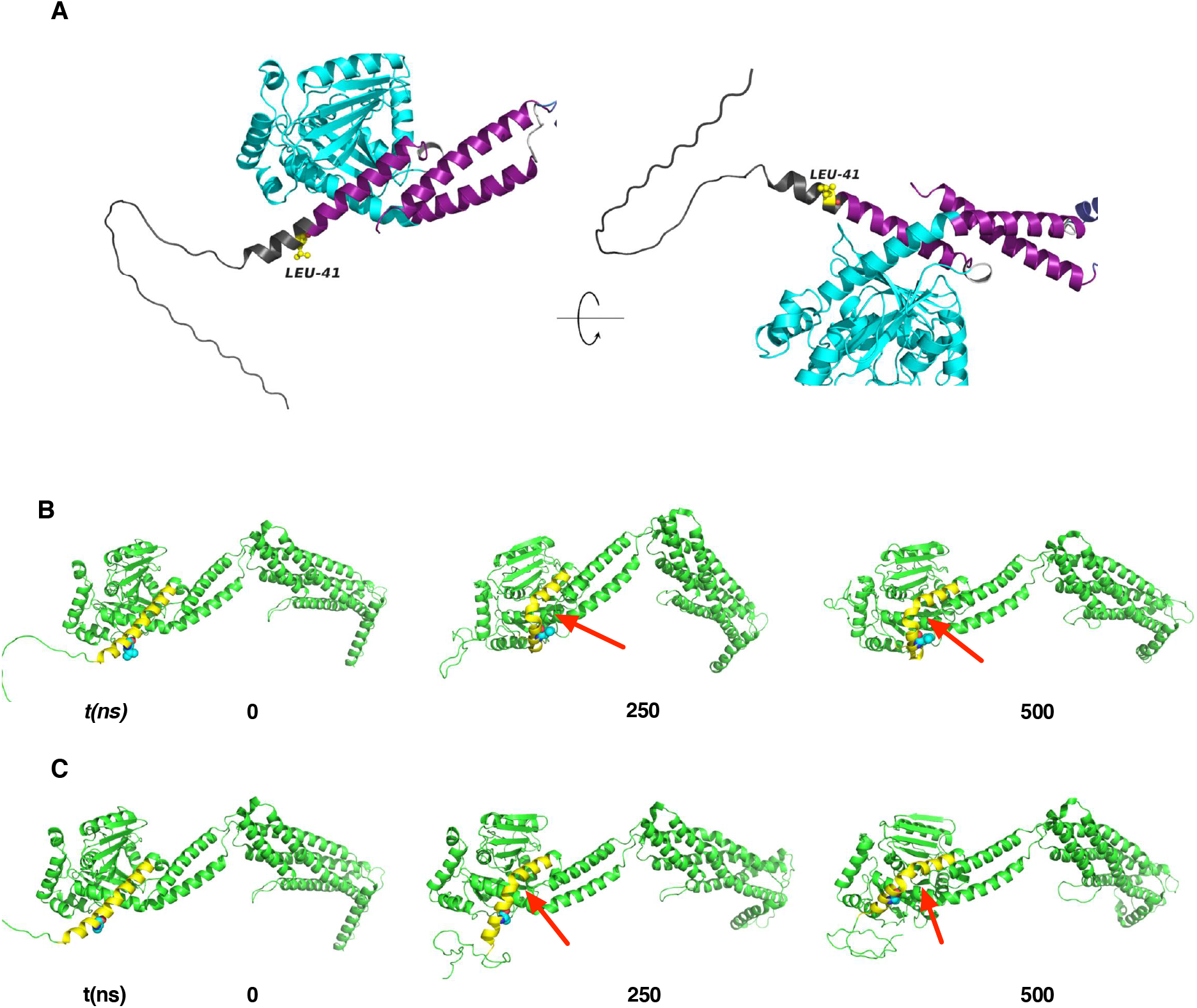
Mutation of L41 of HsMX1, which is predicted to be in an extended portion of the first BSE helix, does not affect BSE structural properties. **A**. Alphafold prediction of HsMX1 N-terminal domain with the position of L41 highlighted in yellow. BSE helices in magenta, G domain in cyan and modelized N-terminal region in black. **B-C**. Representative snapshots from MD simulations for WT HsMX1 (**B**) and HsMX1_L41A_ (**C**) at three different time points (0, 250 and 500 ns). The N-terminal α-helix is highlighted in yellow and leucine 41 is depicted as cyan spheres. The secondary structure brake or turn occurring within this helix for both WT HsMX1 and HsMX1_L41A_ is pointed by red arrows.

## Discussion

Several HsMX1 intrinsic antiviral determinants have been described in the past, namely an active GTPase domain, the capability to oligomerize via the stalk, and intact BSE, L2 and L4 loops, but the importance of the N-terminal domain had never been assessed. In the case of HsMX2, the N-terminal domain was found to be one of the two main determinants, together with the oligomerization capacity, for anti-HIV activity (36, 37). Therefore, it was plausible that HsMX1 might also need this domain for antiviral activity despite a high variability between orthologous MX proteins (Fig. 1B, 4E, Sup. Fig. 1). Herein, we show that the N-terminal domain, more specifically the highly conserved leucine situated 3 amino acids before the start of the first BSE helix, from human, mouse and bat (*Sturnira lilium*) MX1 proteins, are essential for their anti-IAV activity. The importance of this leucine is highlighted by the fact that its mutation induces a complete loss of anti-IAV activity for all three proteins without being correlated with a decrease in protein expression levels, or, for HsMX1 and MmMx1, without being correlated to a defect of oligomerization or of total inability to hydrolyse GTP. In addition, according to MD simulations based on the AlphaFold prediction, the mutation of L41 in HsMX1 into an alanine does not seem to affect the structure of the first BSE α-helix. Nevertheless, this residue was shown to be essential for subcellular localization of these three MX1 proteins. Finally, we show that this leucine is not only essential for HsMX1 restriction of IAV but also VSV, as well as LACV or BUNV nucleoprotein aggregation capacity.

The N-terminal domain is not present in the crystal structure of HsMX1 (Fig. 1A), the non-structured and highly flexible nature of this extension likely explains the lack of structural information (Fig. 1A, 7A, 7B, Sup. Movie. 1). Furthermore, MmMx1 and SlMX1 3D structures have not yet been determined. As we now show that this conserved leucine is essential for the antiviral activity of these proteins, it would therefore be interesting to be able to decipher experimentally the structure of an MX1 protein with an intact N-terminal domain. This would allow us to understand the three-dimensional architecture of this part of the protein and confirm the AlphaFold predictions. In terms of the role of this residue for MX1 function, we show above that it is not essential for the oligomerization of both HsMX1 and MmMx1. Of note, the elution profiles of the recombinant proteins are very similar for the WT and mutant proteins, confirming our crosslink data and the fact that the leucine point mutants do not seem to affect oligomerization status (Fig. 6A, 5B, 5D). AlphaFold predictions nevertheless suggest that L41 is part of the α-helix of the first BSE (Fig. 7A), which, if confirmed experimentally, should lead to a redefinition of the boundaries of MX1 N-terminal and BSE domains. A plausible effect of these leucine mutants could therefore be a partial hindrance to the correct folding or function of the first helix of the BSE, but MD simulations in this region did not support this hypothesis as similar structural behaviour was observed in both cases (Fig. 7B, 7C, Sup. Movie 1-2). The AlphaFold prediction models the conserved leucine as pointing away from the core of the protein (Fig. 7A). This could potentially suggest that L41 is a critical residue important for interaction with a cellular cofactor or a viral element, although the latter possibility is more difficult to imagine due to the wide breadth of viral families inhibited by MX1 proteins (7, 55). In support of the former idea, the subcellular localization of HsMX1_L41A_, MmMx1_L7A_, and SlMX1_L39A_ were all displaced compared to their WT counterparts (Fig. 1D, 4B, 4D), which might be explained by the loss of interaction with such a cellular cofactor, which would be essential for correct subcellular localization. Because of this, these single point leucine mutants therefore present themselves as ideal candidates for differential interactomic approaches to identify such putative cellular cofactors. To date, no cofactor that would be essential for MX1 effector antiviral activity has been identified. The discovery of such cofactor(s), if they exist, would probably be a major turning point to understand the mechanism of action of MX1 proteins.

Allelic variation studies have been performed for HsMX1 (60, 61), with only one variant being discovered in the N-terminal domain where a premature stop codon was found at amino acid position 30 (Q30*) causing a truncated, antivirally inactive protein (61). Using the gnomAD browser (https://gnomad.broadinstitute.org), a single variant can be found in a single individual implicating L41 of HsMX1 where a missense deletion of 1 base occurred (21-42807777-AC-A) inducing a frameshift. With no reported naturally occurring allelic variation occurring at this position so far, it would be interesting to find such allelic variants to see if these individuals would be more susceptible to viral infections due to a non-functional HsMX1 protein.

The amino acid placed exactly three residues before the (previously defined) beginning of the first BSE α-helix is consistently a leucine for almost all species, with the exception of some fish species where it is replaced by a phenylalanine (Fig. 4E, Sup. Fig. 1). The difference of the genetic code between leucine and phenylalanine is minimal (UUA or UUG for the former and UUU or UUC for the latter). A single point mutation would have therefore been necessary to induce this exchange of amino acids. Another interesting observation is the fact that when the N-terminal domain of HsMX2 was transferred onto HsMX1, this chimeric protein retained its anti-IAV activity while gaining anti-HIV activity (26). Despite being much longer than the N-terminal domain of HsMX1, that of HsMX2 also contains a leucine, 3 amino acids before start of the BSE, which was also the case for the chimeric protein (26). This information, coupled with the fact that the rest of the N-terminal domain is extremely divergent between species, attests to the high evolutionary selective pressure placed on this residue, further reinforcing the importance of this leucine for MX1 protein antiviral activity.

In conclusion, we show herein that the N-terminal domain of MX1 proteins, more specifically a highly conserved leucine residue, is important for their antiviral activity against multiple RNA viruses and governs correct subcellular localization. Further characterizing the importance of this residue could pave the way to a better understanding of the molecular mechanisms at play in the antiviral activity of these long-studied, yet still not well understood restriction factors.

## Acknowledgements

We are thankful to Prof. Georg Kochs (Freiburg University, Germany) for the pcDNA3-RFP-LACV-N and SlMX1 plasmids; to Prof. Gert Zimmer (Institute of Virology and Immunology, Switzerland) for the G-pseudotyped VSVΔG-GFP-Firefly Luciferase virus. This work was supported by the European Research Council (ERC) under the European Union’s Horizon 2020 research and innovation programme (grant agreement 759226, ANTIViR, to CG), the ATIP-Avenir program (to CG), the Institut National de la Santé et de la Recherche Médicale (INSERM) (to CG), institutional funds from the Centre National de la Recherche Scientifique (CNRS) and Montpellier University, a 3-year PhD studentship from the Ministry of Higher Education and Research (to JM) and a 4th year PhD funding from by the Fondation pour la Recherche Médicale, grant number [FDT202106013175] (to JM), and institutional funds from IRIM (inter-team project, to MAA and MB). We acknowledge the imaging facility MRI, member of the national infrastructure France-BioImaging supported by the French National Research Agency (ANR-10-INBS-04). JM, OM and CG acknowledge support from the French research network on influenza viruses (ResaFlu; GDR2073) financed by the CNRS.

## Author contributions

J.M., O.M. and C.G. conceived the study, designed the experiments and interpreted the data; J.M. performed the experiments; M.A-A performed immunoblots, produced the recombinant proteins and did the GTPase assays; M.B. provided expertise and supervision on recombinant protein production and biochemical assays; M.T., C.A-A., and O.P. provided technical help; L.C. performed and analysed the MD studies. J.M., O.M. and C.G. wrote the manuscript with input from all authors.

## Footnotes

3D: Three-dimensional

BSE: Bundle Signalling Element

BUNV: Bunyamwera Virus

CDS: Coding DNA Sequence

DMEM: Dulbecco’s Modified Eagle Medium

DMSO: DiMethyl SulfOxide

DNA: DeoxyriboNucleic Acid

DSS: DiSuccinimidyl Suberate

EDTA: EthyleneDiamineTetraacetic Acid

EM: Electron Microscopy

FBS: Fetal Bovine Serum

GDP: Guanosine DiPhosphate

GFP: Green Fluorescent Protein

GTP: Guanosine TriPhosphate

GTPase: Guanosine TriPhosphatase

HBV: Hepatitis B Virus

HEK293T: Human Embryonic Kidney 293T

HIV-1: Human Immunodeficiency Virus 1

HRP: HorseRadish Peroxidase

HsMX1/2: Human Myxovirus resistance protein 1/2

HTNV: Hantaan River Virus

IAV: Influenza A virus

IFN: Interferon

ISG: Interferon Stimulated Genes

IPTG: IsoPropyl-β-D-ThioGalactoside

JAK: Janus Kinase

L7/39/41: Leucine 7/39/41

LACV: La Crosse Virus

LACV/BUNV-N: LACV/BUNV Nucleoprotein

LB: Lysogeny Broth

LV: Lentiviral Vector

MD: Molecular Dynamics

MDCK: Madin-Darby Canine Kidney

MmMx1/2: Mouse Myxovirus resistance protein 1/2

MOI: Multiplicity of Infection

MX1/2: Myxovirus resistance protein 1/2

MxA/B: Myxovirus resistance protein

A/B N-terminal: Amino-terminal

NLS: Nuclear Localization Signal

NLuc: Nanoluciferase

NP: NucleoProtein

PA: Polymerase Acidic

PB1/2: Polymerase Basic 1/2

PBS: Phosphate Buffered Saline

PFA: ParaFormAldehyde

PH: Pleckstrin Homology

PI(4,5)P2: PhosphatidylInositol 4,5-bisPhosphate

RIG-I: Retinoic acid-Inducible Gene I

RFP: Red Fluorescent Protein

RMSD: Root Mean Square Deviation

RNA: RiboNucleic Acid

RT: Room Temperature S35: Serine 35

SDS-PAGE: Sodium Dodecyl Sulfate–Polyacrylamide Gel Electrophoresis

SlMX1: Sturnira lilium Myxovirus resistance protein 1

STAT: Signal Transducer and Activator of Transcription

TEV: Tobacco Etch Virus

THOV: Thogoto Virus

TPCK: N-Tosyl-L-Phenylalanine Chloromethyl Ketone

VSV: Vesicular Stomatitis Virus

WT: Wild-Type

## References

1. Schoggins, J. W. (2019) Interferon-Stimulated Genes: What Do They All Do? Annu Rev Virol. 6, 567–584

2. McKellar, J., Rebendenne, A., Wencker, M., Moncorgé, O., and Goujon, C. (2021) Mammalian and Avian Host Cell Influenza A Restriction Factors. Viruses. 13, 522

3. Lindenmann, J. (1962) Resistance of mice to mouse-adapted influenza A virus. Virology. 16, 203–204

4. Horisberger, M. A., Staeheli, P., and Haller, O. (1983) Interferon induces a unique protein in mouse cells bearing a gene for resistance to influenza virus. Proceedings of the National Academy of Sciences. 80, 1910–1914

5. Staeheli, P., Haller, O., Boll, W., Lindenmann, J., and Weissmann, C. (1986) Mx protein: Constitutive expression in 3T3 cells transformed with cloned Mx cDNA confers selective resistance to influenza virus. Cell. 44, 147–158

6. Haller, O., Arnheiter, H., Pavlovic, J., and Staeheli, P. (2018) The Discovery of the Antiviral Resistance Gene Mx: A Story of Great Ideas, Great Failures, and Some Success. Annu Rev Virol. 5, 33–51

7. Haller, O., Staeheli, P., Schwemmle, M., and Kochs, G. (2015) Mx GTPases: dynamin-like antiviral machines of innate immunity. Trends Microbiol. 23, 154–163

8. Goujon, C., Moncorgé, O., Bauby, H., Doyle, T., Ward, C. C., Schaller, T., Hué, S., Barclay, W. S., Schulz, R., and Malim, M. H. (2013) Human MX2 is an interferon-induced post-entry inhibitor of HIV-1 infection. Nature. 502, 559–562

9. Kane, M., Yadav, S. S., Bitzegeio, J., Kutluay, S. B., Zang, T., Wilson, S. J., Schoggins, J. W., Rice, C. M., Yamashita, M., Hatziioannou, T., and Bieniasz, P. D. (2013) MX2 is an interferon-induced inhibitor of HIV-1 infection. Nature. 502, 563–566

10. Liu, Z., Pan, Q., Ding, S., Qian, J., Xu, F., Zhou, J., Cen, S., Guo, F., and Liang, C. (2013) The interferon-inducible MxB protein inhibits HIV-1 infection. Cell Host Microbe. 14, 398–410

11. Schilling, M., Bulli, L., Weigang, S., Graf, L., Naumann, S., Patzina, C., Wagner, V., Bauersfeld, L., Goujon, C., Hengel, H., Halenius, A., Ruzsics, Z., Schaller, T., and Kochs, G. (2018) Human MxB Protein Is a Pan-herpesvirus Restriction Factor. J Virol. 10.1128/JVI.01056-18

12. Crameri, M., Bauer, M., Caduff, N., Walker, R., Steiner, F., Franzoso, F. D., Gujer, C., Boucke, K., Kucera, T., Zbinden, A., Münz, C., Fraefel, C., Greber, U. F., and Pavlovic, J. (2018) MxB is an interferon-induced restriction factor of human herpesviruses. Nat Commun. 9, 1980

13. Serrero, M. C., Girault, V., Weigang, S., Greco, T. M., Ramos Nascimento, A., Anderson, F., Piras, A., Hickford Martinez, A., Hertzog, J., Binz, A., Pohlmann, A., Prank, U., Rehwinkel, J., Bauerfeind, R., Cristea, I. M., Pichlmair, A., Kochs, G., and Sodeik, B. (2022) The interferon-inducible GTPase MxB promotes capsid disassembly and genome release of herpesviruses. Elife. 11, e76804

14. Zimmermann, P., Mänz, B., Haller, O., Schwemmle, M., and Kochs, G. (2011) The viral nucleoprotein determines Mx sensitivity of influenza A viruses. J Virol. 85, 8133–8140

15. Engelhardt, O. G., Ullrich, E., Kochs, G., and Haller, O. (2001) Interferon-Induced Antiviral Mx1 GTPase Is Associated with Components of the SUMO-1 System and Promyelocytic Leukemia Protein Nuclear Bodies. Experimental Cell Research. 271, 286–295

16. Engelhardt, O. G., Sirma, H., Pandolfi, P.-P., and Haller, O. (2004) Mx1 GTPase accumulates in distinct nuclear domains and inhibits influenza A virus in cells that lack promyelocytic leukaemia protein nuclear bodies. Journal of General Virology. 85, 2315–2326

17. Cao, Y.-L., Meng, S., Chen, Y., Feng, J.-X., Gu, D.-D., Yu, B., Li, Y.-J., Yang, J.-Y., Liao, S., Chan, D. C., and Gao, S. (2017) MFN1 structures reveal nucleotide-triggered dimerization critical for mitochondrial fusion. Nature. 542, 372–376

18. Ramachandran, R., and Schmid, S. L. (2018) The dynamin superfamily. Curr Biol. 28, R411–R416

19. Gao, S., von der Malsburg, A., Paeschke, S., Behlke, J., Haller, O., Kochs, G., and Daumke, O. (2010) Structural basis of oligomerization in the stalk region of dynamin-like MxA. Nature. 465, 502–506

20. Gao, S., von der Malsburg, A., Dick, A., Faelber, K., Schröder, G. F., Haller, O., Kochs, G., and Daumke, O. (2011) Structure of myxovirus resistance protein a reveals intra-and intermolecular domain interactions required for the antiviral function. Immunity. 35, 514–525

21. Chen, Y., Zhang, L., Graf, L., Yu, B., Liu, Y., Kochs, G., Zhao, Y., and Gao, S. (2017) Conformational dynamics of dynamin-like MxA revealed by single-molecule FRET. Nat Commun. 8, 15744

22. Zheng, J., Cahill, S. M., Lemmon, M. A., Fushman, D., Schlessinger, J., and Cowburn, D. (1996) Identification of the binding site for acidic phospholipids on the pH domain of dynamin: implications for stimulation of GTPase activity. J Mol Biol. 255, 14–21

23. Kochs, G., Haener, M., Aebi, U., and Haller, O. (2002) Self-assembly of Human MxA GTPase into Highly Ordered Dynamin-like Oligomers. Journal of Biological Chemistry. 277, 14172–14176

24. Zürcher, T., Pavlovic, J., and Staeheli, P. (1992) Nuclear localization of mouse Mx1 protein is necessary for inhibition of influenza virus. J Virol. 66, 5059–5066

25. Pitossi, F., Blank, A., Schröder, A., Schwarz, A., Hüssi, P., Schwemmle, M., Pavlovic, J., and Staeheli, P. (1993) A functional GTP-binding motif is necessary for antiviral activity of Mx proteins. J Virol. 67, 6726–6732

26. Goujon, C., Moncorgé, O., Bauby, H., Doyle, T., Barclay, W. S., and Malim, M. H. (2014) Transfer of the amino-terminal nuclear envelope targeting domain of human MX2 converts MX1 into an HIV-1 resistance factor. J Virol. 88, 9017–9026

27. Yu, Z., Wang, Z., Chen, J., Li, H., Lin, Z., Zhang, F., Zhou, Y., and Hou, J. (2008) GTPase activity is not essential for the interferon-inducible MxA protein to inhibit the replication of hepatitis B virus. Arch Virol. 153, 1677–1684

28. Ponten, A., Sick, C., Weeber, M., Haller, O., and Kochs, G. (1997) Dominant-negative mutants of human MxA protein: domains in the carboxy-terminal moiety are important for oligomerization and antiviral activity. J Virol. 71, 2591–2599

29. Arnheiter, H., and Haller, O. (1988) Antiviral state against influenza virus neutralized by microinjection of antibodies to interferon-induced Mx proteins. EMBO J. 7, 1315–1320

30. Flohr, F., Schneider-Schaulies, S., Haller, O., and Kochs, G. (1999) The central interactive region of human MxA GTPase is involved in GTPase activation and interaction with viral target structures. FEBS Letters. 463, 24–28

31. Kochs, G., and Haller, O. (1999) Interferon-induced human MxA GTPase blocks nuclear import of Thogoto virus nucleocapsids. Proc Natl Acad Sci U S A. 96, 2082–2086

32. Garber, E. A., Hreniuk, D. L., Scheidel, L. M., and van der Ploeg, L. H. (1993) Mutations in murine Mx1: effects on localization and antiviral activity. Virology. 194, 715–723

33. Patzina, C., Haller, O., and Kochs, G. (2014) Structural requirements for the antiviral activity of the human MxA protein against Thogoto and influenza A virus. J Biol Chem. 289, 6020–6027

34. Verhelst, J., Spitaels, J., Nürnberger, C., De Vlieger, D., Ysenbaert, T., Staeheli, P., Fiers, W., and Saelens, X. (2015) Functional Comparison of Mx1 from Two Different Mouse Species Reveals the Involvement of Loop L4 in the Antiviral Activity against Influenza A Viruses. J Virol. 89, 10879–10890

35. Mitchell, P. S., Patzina, C., Emerman, M., Haller, O., Malik, H. S., and Kochs, G. (2012) Evolution-guided identification of antiviral specificity determinants in the broadly acting interferon-induced innate immunity factor MxA. Cell Host Microbe. 12, 598–604

36. Goujon, C., Greenbury, R. A., Papaioannou, S., Doyle, T., and Malim, M. H. (2015) A triple-arginine motif in the amino-terminal domain and oligomerization are required for HIV-1 inhibition by human MX2. J Virol. 89, 4676–4680

37. Busnadiego, I., Kane, M., Rihn, S. J., Preugschas, H. F., Hughes, J., Blanco-Melo, D., Strouvelle, V. P., Zang, T. M., Willett, B. J., Boutell, C., Bieniasz, P. D., and Wilson, S. J. (2014) Host and viral determinants of Mx2 antiretroviral activity. J Virol. 88, 7738–7752

38. Fuchs, J., Hölzer, M., Schilling, M., Patzina, C., Schoen, A., Hoenen, T., Zimmer, G., Marz, M., Weber, F., Müller, M. A., and Kochs, G. (2017) Evolution and Antiviral Specificities of Interferon-Induced Mx Proteins of Bats against Ebola, Influenza, and Other RNA Viruses. J Virol. 10.1128/JVI.00361-17

39. Doyle, T., Moncorgé, O., Bonaventure, B., Pollpeter, D., Lussignol, M., Tauziet, M., Apolonia, L., Catanese, M.-T., Goujon, C., and Malim, M. H. (2018) The interferon-inducible isoform of NCOA7 inhibits endosome-mediated viral entry. Nat Microbiol. 3, 1369–1376

40. Schindelin, J., Arganda-Carreras, I., Frise, E., Kaynig, V., Longair, M., Pietzsch, T., Preibisch, S., Rueden, C., Saalfeld, S., Schmid, B., Tinevez, J.-Y., White, D. J., Hartenstein, V., Eliceiri, K., Tomancak, P., and Cardona, A. (2012) Fiji: an open-source platform for biological-image analysis. Nat Methods. 9, 676–682

41. Berger Rentsch, M., and Zimmer, G. (2011) A Vesicular Stomatitis Virus Replicon-Based Bioassay for the Rapid and Sensitive Determination of Multi-Species Type I Interferon. PLoS ONE. 6, e25858

42. Neumann, G., and Hobom, G. (1995) Mutational analysis of influenza virus promoter elements in vivo. J Gen Virol. 76 (Pt 7), 1709–1717

43. Dicks, M. D. J., Goujon, C., Pollpeter, D., Betancor, G., Apolonia, L., Bergeron, J. R. C., and Malim, M. H. (2016) Oligomerization Requirements for MX2-Mediated Suppression of HIV-1 Infection. J. Virol. 90, 22–32

44. Jumper, J., Evans, R., Pritzel, A., Green, T., Figurnov, M., Ronneberger, O., Tunyasuvunakool, K., Bates, R., Žídek, A., Potapenko, A., Bridgland, A., Meyer, C., Kohl, S. A., Ballard, A. J., Cowie, A., Romera-Paredes, B., Nikolov, S., Jain, R., Adler, J., Back, T., Petersen, S., Reiman, D., Clancy, E., Zielinski, M., Steinegger, M., Pacholska, M., Berghammer, T., Bodenstein, S., Silver, D., Vinyals, O., Senior, A. W., Kavukcuoglu, K., Kohli, P., and Hassabis, D. (2021) Highly accurate protein structure prediction with AlphaFold. Nature. 596, 583–589

45. Mirdita, M., Schütze, K., Moriwaki, Y., Heo, L., Ovchinnikov, S., and Steinegger, M. (2021) ColabFold -Making protein folding accessible to all, Bioinformatics, 10.1101/2021.08.15.456425

46. Šali, A., and Blundell, T. L. (1993) Comparative Protein Modelling by Satisfaction of Spatial Restraints. Journal of Molecular Biology. 234, 779–815

47. Phillips, J. C., Hardy, D. J., Maia, J. D. C., Stone, J. E., Ribeiro, J. V., Bernardi, R. C., Buch, R., Fiorin, G., Hénin, J., Jiang, W., McGreevy, R., Melo, M. C. R., Radak, B. K., Skeel, R. D., Singharoy, A., Wang, Y., Roux, B., Aksimentiev, A., Luthey-Schulten, Z., Kalé, L. V., Schulten, K., Chipot, C., and Tajkhorshid, E. (2020) Scalable molecular dynamics on CPU and GPU architectures with NAMD. J. Chem. Phys. 153, 044130

48. Huang, J., Rauscher, S., Nawrocki, G., Ran, T., Feig, M., de Groot, B. L., Grubmüller, H., and MacKerell, A. D. (2017) CHARMM36m: an improved force field for folded and intrinsically disordered proteins. Nat Methods. 14, 71–73

49. Tuckerman, M., Berne, B. J., and Martyna, G. J. (1992) Reversible multiple time scale molecular dynamics. The Journal of Chemical Physics. 97, 1990–2001

50. Essmann, U., Perera, L., Berkowitz, M. L., Darden, T., Lee, H., and Pedersen, L. G. (1995) A smooth particle mesh Ewald method. The Journal of Chemical Physics. 103, 8577–8593

51. Humphrey, W., Dalke, A., and Schulten, K. (1996) VMD: Visual molecular dynamics. Journal of Molecular Graphics. 14, 33–38

52. Nigg, P. E., and Pavlovic, J. (2015) Oligomerization and GTP-binding Requirements of MxA for Viral Target Recognition and Antiviral Activity against Influenza A Virus. Journal of Biological Chemistry. 290, 29893–29906

53. Dick, A., Graf, L., Olal, D., von der Malsburg, A., Gao, S., Kochs, G., and Daumke, O. (2015) Role of Nucleotide Binding and GTPase Domain Dimerization in Dynamin-like Myxovirus Resistance Protein A for GTPase Activation and Antiviral Activity. Journal of Biological Chemistry. 290, 12779–12792

54. Kochs, G., Janzen, C., Hohenberg, H., and Haller, O. (2002) Antivirally active MxA protein sequesters La Crosse virus nucleocapsid protein into perinuclear complexes. Proc Natl Acad Sci U S A. 99, 3153–3158

55. Verhelst, J., Hulpiau, P., and Saelens, X. (2013) Mx Proteins: Antiviral Gatekeepers That Restrain the Uninvited. Microbiol Mol Biol Rev. 77, 551–566

56. Schneider, T. D., and Stephens, R. M. (1990) Sequence logos: a new way to display consensus sequences. Nucleic Acids Res. 18, 6097–6100

57. Crooks, G. E., Hon, G., Chandonia, J.-M., and Brenner, S. E. (2004) WebLogo: a sequence logo generator. Genome Res. 14, 1188–1190

58. Janzen, C., Kochs, G., and Haller, O. (2000) A Monomeric GTPase-Negative MxA Mutant with Antiviral Activity. J Virol. 74, 8202–8206

59. Varadi, M., Anyango, S., Deshpande, M., Nair, S., Natassia, C., Yordanova, G., Yuan, D., Stroe, O., Wood, G., Laydon, A., Žídek, A., Green, T., Tunyasuvunakool, K., Petersen, S., Jumper, J., Clancy, E., Green, R., Vora, A., Lutfi, M., Figurnov, M., Cowie, A., Hobbs, N., Kohli, P., Kleywegt, G., Birney, E., Hassabis, D., and Velankar, S. (2022) AlphaFold Protein Structure Database: massively expanding the structural coverage of protein-sequence space with high-accuracy models. Nucleic Acids Res. 50, D439–D444

60. Graf, L., Dick, A., Sendker, F., Barth, E., Marz, M., Daumke, O., and Kochs, G. (2018) Effects of allelic variations in the human myxovirus resistance protein A on its antiviral activity. J Biol Chem. 293, 3056–3072

61. Chen, Y., Graf, L., Chen, T., Liao, Q., Bai, T., Petric, P. P., Zhu, W., Yang, L., Dong, J., Lu, J., Chen, Y., Shen, J., Haller, O., Staeheli, P., Kochs, G., Wang, D., Schwemmle, M., and Shu, Y. (2021) Rare variant MX1 alleles increase human susceptibility to zoonotic H7N9 influenza virus. Science. 373, 918–922

