## Supplementary Figure 1 for "The N-terminal domain of MX1 proteins is essential for their antiviral activity against different families of RNA viruses"

| Entry | Organism | Entry | Organism |
|---|---|---|---|
| P20591 | Homo sapiens (Human) | S7NSW7 | Myotis brandtii (Brandt's bat) |
| K4JBQ0 | Gorilla gorilla (western gorilla) | A0A1N7THY9 | Pipistrellus pipistrellus (Common pipistrelle) |
| H2QL18 | Pan troglodytes (Chimpanzee) | A0A1N7THZ0 | Rousettus aegyptiacus (Egyptian rousette) |
| A1E2I4 | Macaca mulatta (Rhesus macaque) | A0A1N7THY7 | Hypsignathus monstrosus (Hammer-headed fruit bat) |
| Q5R5G3 | Pongo abelii (Sumatran orangutan) | A0A1N7THZ1 | Sturnira lilium (Lesser yellow-shouldered bat) |
| A0A6P8QPW8 | Geotrypetes seraphini (Gaboon caecilian) | B2X026 | Gallus gallus (Chicken) |
| U3EGN4 | Callithrix jacchus (White-tufted-ear marmoset) | A0A172PZ65 | Anser cygnoid (Swan goose) |
| Q9N0Y3 | Canis lupus familiaris (Dog) | A0A0Q3PUG8 | Amazona aestiva (Blue-fronted Amazon parrot) |
| M3W8X0 | Felis catus (Cat) | Q4ADG6 | Phoca vitulina (Harbor seal) |
| A0A667GNL8 | Lynx canadensis (Canada lynx) | Q4ADG7 | Otaria byronia (South American sea lion) |
| A0A6P6GZN4 | Puma concolor (Mountain lion) | A0A2Y9G772 | Neomonachus schauinslandi (Hawaiian monk seal) |
| A0A6J1XXY6 | Acinonyx jubatus (Cheetah) | A0A6J2C7I6 | Zalophus californianus (California sealion) |
| A0A6P4TS13 | Panthera pardus (Leopard) | A0A2U3WDI8 | Odobenus rosmarus divergens (Pacific walrus) |
| A0A3Q7SCG7 | Vulpes vulpes (Red fox) | A0A1U7SNC8 | Alligator sinensis (Chinese alligator) |
| I3MJX6 | Ictidomys tridecemlineatus (Thirteen-lined ground squirrel) | A0A7M4FFF4 | Crocodylus porosus (Saltwater crocodile) |
| G9KBY2 | Mustela putorius furo (European domestic ferret) | A0A340WTU5 | Lipotes vexillifer (Yangtze river dolphin) |
| A0A2Y9KR88 | Enhydra lutris kenyoni (sea otter) | A0A6J3RAX5 | Tursiops truncatus (Atlantic bottle-nosed dolphin) |
| P09922 | Mus musculus (Mouse) | A0A341CI24 | Neophocaena asiaeorientalis asiaeorientalis (Yangtze finless porpoise) |
| P18588 | Rattus norvegicus (Rat) | A0A2Y9LTL6 | Delphinapterus leucas (Beluga whale) |
| A0A6P6D2J7 | Pteropus vampyrus (Large flying fox) | A0A383ZCI9 | Balaenoptera acutorostrata scammoni (North Pacific minke whale) |
| A0A6J0DKA3 | Peromyscus maniculatus bairdii (Prairie deer mouse) | V9KIB1 | Callorhinchus milii (Ghost shark) |
| G5C5C9 | Heterocephalus glaber (Naked mole rat) | A0A674K7E1 | Terrapene carolina triunguis (Three-toed box turtle) |
| A0A3Q0CWV5 | Mesocricetus auratus (Golden hamster) | A0A4D9DW37 | Platysternon megacephalum (big-headed turtle) |
| A0A6I9K5X3 | Chrysochloris asiatica (Cape golden mole) | H2DQ54 | Xenopus laevis (African clawed frog) |
| P79135 | Bos taurus (Bovine) | A0A0F6MUS9 | Anguilla japonica (Japanese eel) |
| A0A6P3H2K4 | Bison bison bison | Q98990 | Salmo salar (Atlantic salmon) |
| L8IPR7 | Bos mutus (wild yak) | A0A674EGU5 | Salmo trutta (Brown trout) |
| P27594 | Sus scrofa (Pig) | Q91192 | Oncorhynchus mykiss (Rainbow trout) |
| A0A1U7U9K3 | Carlito syrichta (Philippine tarsier) | A0A286K200 | Mylopharyngodon piceus (Black carp) |
| P33237 | Ovis aries (Sheep) | A0A498MHV8 | Labeo rohita (Indian major carp) |
| A0A7D4WPX4 | Capra hircus (Goat) | Q7T2P0 | Ictalurus punctatus (Channel catfish) |
| Q28379 | Equus caballus (Horse) | A0A556TK66 | Bagarius yarrelli (Goonch) |
| A0A6I9I6Y4 | Vicugna pacos (Alpaca) | A0A3G6IID3 | Squaliobarbus curriculus (barbel chub) |
| A0A6P5BYP3 | Bos indicus (Zebu) | Q8JH68 | Danio rerio (Zebrafish) (Brachydanio rerio) |
| S9XPZ9 | Camelus ferus (Wild bactrian camel) | A0A672MEY2 | Sinocyclocheilus grahami (Dianchi golden-line fish) |
| A0A384DMX3 | Ursus maritimus (Polar bear) | Q7T2M3 | Carassius auratus (Goldfish) |
| A0A3Q7W4X3 | Ursus arctos horribilis (Grizzly bear) | A0A673LWX8 | Sinocyclocheilus rhinocerous |
| A0A7E6D288 | Phyllostomus discolor (pale spear-nosed bat) | A0A8B9KF29 | Astyanax mexicanus (Blind cave fish) |

CLUSTAL O(1.2.4) multiple sequence alignment

```
SP|P20591|MX1_HUMAN                 ------------------------------------------------------------
TR|K4JBQ0|K4JBQ0_9PRIM              ------------------------------------------------------------
TR|H2QL18|H2QL18_PANTR              ------------------------------------------------------------
SP|A1E2I4|MX1_MACMU                 ------------------------------------------------------------
SP|Q5R5G3|MX1_PONAB                 ------------------------------------------------------------
TR|A0A6P8QPW8|A0A6P8QPW8_GEOSA      ------------------------------------------------------------
TR|U3EGN4|U3EGN4_CALJA              ------------------------------------------------------------
SP|Q9N0Y3|MX1_CANLF                 ------------------------------------------------------------
TR|M3W8X0|M3W8X0_FELCA              ------------------------------------------------------------
TR|A0A667GNL8|A0A667GNL8_LYNCA      ------------------------------------------------------------
TR|A0A6P6GZN4|A0A6P6GZN4_PUMCO      ------------------------------------------------------------
TR|A0A6J1XXY6|A0A6J1XXY6_ACIJB      ------------------------------------------------------------
TR|A0A6P4TS13|A0A6P4TS13_PANPR      ---------------------------------------M-----------TYQKQP--- 7
TR|A0A3Q7SCG7|A0A3Q7SCG7_VULVU      ---------------------------------------M-----------AASRGP--- 7
TR|I3MJX6|I3MJX6_ICTTR              ------------------------------------------------------------
TR|G9KBY2|G9KBY2_MUSPF              ------------------------------------------------------------
TR|A0A2Y9KR88|A0A2Y9KR88_ENHLU      ------------------------------------------------------------
SP|P09922|MX1B_MOUSE                ------------------------------------------------------------
SP|P18588|MX1_RAT                   ------------------------------------------------------------
TR|A0A6P6D2J7|A0A6P6D2J7_PTEVA      ------------------------------------------------------------
TR|A0A6J0DKA3|A0A6J0DKA3_PERMB      ------------------------------------------------------------
TR|G5C5C9|G5C5C9_HETGA              ------------------------------------------------------------
TR|A0A3Q0CWV5|A0A3Q0CWV5_MESAU      ---------------------------------------M-----------SNTKTP--- 7
TR|A0A6I9K5X3|A0A6I9K5X3_CHRAS      ------------------------------------------------------------
SP|P79135|MX1_BOVIN                 ------------------------------------------------------------
TR|A0A6P3H2K4|A0A6P3H2K4_BISBI      ------------------------------------------------------------
TR|L8IPR7|L8IPR7_9CETA              ------------------------------------------------------------
SP|P27594|MX1_PIG                   ------------------------------------------------------------
TR|A0A1U7U9K3|A0A1U7U9K3_CARSF      ------------------------------------------------------------
SP|P33237|MX1_SHEEP                 ------------------------------------------------------------
TR|A0A7D4WPX4|A0A7D4WPX4_CAPHI      ------------------------------------------------------------
SP|Q28379|MX1_HORSE                 ------------------------------------------------------------
TR|A0A6I9I6Y4|A0A6I9I6Y4_VICPA      ------------------------------------------------------------
TR|A0A6P5BYP3|A0A6P5BYP3_BOSIN      ------------------------------------------------------------
TR|S9XPZ9|S9XPZ9_CAMFR              ------------------------------------------------------------
TR|A0A384DMX3|A0A384DMX3_URSMA      ------------------------------------------------------------
TR|A0A3Q7W4X3|A0A3Q7W4X3_URSAR      ------------------------------------------------------------
TR|A0A7E6D288|A0A7E6D288_9CHIR      ------------------------------------------------------------
TR|S7NSW7|S7NSW7_MYOBR              ------------------------------------------------------------
TR|A0A1N7THY9|A0A1N7THY9_PIPPI      ------------------------------------------------------------
TR|A0A1N7THZ0|A0A1N7THZ0_ROUAE      ------------------------------------------------------------
TR|A0A1N7THY7|A0A1N7THY7_HYPMN      ------------------------------------------------------------
TR|A0A1N7THZ1|A0A1N7THZ1_STULI      ------------------------------------------------------------
TR|B2X026|B2X026_CHICK              ------------------------------------------------------------
TR|A0A172PZ65|A0A172PZ65_ANSCY      ------------------------------------------------------------
TR|A0A0Q3PUG8|A0A0Q3PUG8_AMAAE      ------------------------------------------------------------
SP|Q4ADG6|MX1_PHOVI                 ------------------------------------------------------------
SP|Q4ADG7|MX1_OTABY                 ------------------------------------------------------------
TR|A0A2Y9G772|A0A2Y9G772_NEOSC      ------------------------------------------------------------
TR|A0A6J2C7I6|A0A6J2C7I6_ZALCA      ------------------------------------------------------------
TR|A0A2U3WDI8|A0A2U3WDI8_ODORO      ------------------------------------------------------------
TR|A0A1U7SNC8|A0A1U7SNC8_ALLSI      ------------------------------------------------------------
TR|A0A7M4FFF4|A0A7M4FFF4_CROPO      ------------------------------------------------------------
TR|A0A340WTU5|A0A340WTU5_LIPVE      ------------------------------------------------------------
TR|A0A6J3RAX5|A0A6J3RAX5_TURTR      ------------------------------------------------------------
TR|A0A341CI24|A0A341CI24_NEOAA      -----MRR-----------------SALRGGPLTQGLESC-----------PGSAFPVAC 27
TR|A0A2Y9LTL6|A0A2Y9LTL6_DELLE      ------------------------------------------------------------
TR|A0A383ZCI9|A0A383ZCI9_BALAS      ------------------------------------------------------------
TR|V9KIB1|V9KIB1_CALMI              ------------------------------------------------------------
TR|A0A674K7E1|A0A674K7E1_TERCA      ------------------------------------------------------------
TR|A0A4D9DW37|A0A4D9DW37_9SAUR      MYKPSLRRMPAPSGPGAYLMAKESQVFLGGTPVIQSNAAFAASLSTQSQAVFAGSFPTQS 60
TR|H2DQ54|H2DQ54_XENLA              ------------------------------------------------------------
TR|A0A0F6MUS9|A0A0F6MUS9_ANGJA      ------------------------------------------------------------
TR|Q98990|Q98990_SALSA              ------------------------------------------------------------
TR|A0A674EGU5|A0A674EGU5_SALTR      ------------------------------------------------------------
SP|Q91192|MX1_ONCMY                 ------------------------------------------------------------
TR|A0A286K200|A0A286K200_MYLPI      ------------------------------------------------------------
TR|A0A498MHV8|A0A498MHV8_LABRO      ------------------------------------------------------------
SP|Q7T2P0|MX1_ICTPU                 ------------------------------------------------------------
TR|A0A556TK66|A0A556TK66_BAGYA      ------------------------------------------------------------
TR|A0A3G6IID3|A0A3G6IID3_9TELE      ------------------------------------------------------------
SP|Q8JH68|MXA_DANRE                 ------------------------------------------------------------
TR|A0A672MEY2|A0A672MEY2_SINGR      ------------------------------------------------------------
TR|Q7T2M3|Q7T2M3_CARAU              ------------------------------------------------------------
TR|A0A673LWX8|A0A673LWX8_9TELE      ------------------------------------------------------------
TR|A0A8B9KF29|A0A8B9KF29_ASTMX      ------------------------------------------------------------


SP|P20591|MX1_HUMAN                 ------------------------------------------------------------
TR|K4JBQ0|K4JBQ0_9PRIM              ------------------------------------------------------------
TR|H2QL18|H2QL18_PANTR              ------------------------------------------------------------
SP|A1E2I4|MX1_MACMU                 ------------------------------------------------------------
SP|Q5R5G3|MX1_PONAB                 ------------------------------------------------------------
TR|A0A6P8QPW8|A0A6P8QPW8_GEOSA      ------------------------------------------------------------
TR|U3EGN4|U3EGN4_CALJA              ------------------------------------------------------------
SP|Q9N0Y3|MX1_CANLF                 ------------------------------------------------------------
```

```
TR|M3W8X0|M3W8X0_FELCA           ------------------------------------------------------------
TR|A0A667GNL8|A0A667GNL8_LYNCA ------------------------------------------------------------
TR|A0A6P6GZN4|A0A6P6GZN4_PUMCO ------------------------------------------------------------
TR|A0A6J1XXY6|A0A6J1XXY6_ACIJB ------------------------------------------------------------
TR|A0A6P4TS13|A0A6P4TS13_PANPR ------PPPPPT------------SRTFLLEEQTWAS----------------------- 26
TR|A0A3Q7SCG7|A0A3Q7SCG7_VULVU ------G-----------------CR---SGGCRWETYYC-------------------- 21
TR|I3MJX6|I3MJX6_ICTTR           ------------------------------------------------------------
TR|G9KBY2|G9KBY2_MUSPF           ------------------------------------------------------------
TR|A0A2Y9KR88|A0A2Y9KR88_ENHLU ------------------------------------------------------------
SP|P09922|MX1B_MOUSE             ------------------------------------------------------------
SP|P18588|MX1_RAT                ------------------------------------------------------------
TR|A0A6P6D2J7|A0A6P6D2J7_PTEVA ------------------------------------------------------------
TR|A0A6J0DKA3|A0A6J0DKA3_PERMB ------MTPPAL------------CR-PCQGPSQWQCFETIL-----NQTLYF----HHK 32
TR|G5C5C9|G5C5C9_HETGA           ------------------------------------------------------------
TR|A0A3Q0CWV5|A0A3Q0CWV5_MESAU ------V--------------------KNTRMSSW----K-------------------- 17
TR|A0A6I9K5X3|A0A6I9K5X3_CHRAS ------------------------------------------------------------
SP|P79135|MX1_BOVIN              ------------------------------------------------------------
TR|A0A6P3H2K4|A0A6P3H2K4_BISBI ------------------------------------------------------------
TR|L8IPR7|L8IPR7_9CETA           ------------------------------------------------------------
SP|P27594|MX1_PIG                ------------------------------------------------------------
TR|A0A1U7U9K3|A0A1U7U9K3_CARSF ------------------------------------------------------------
SP|P33237|MX1_SHEEP              ------------------------------------------------------------
TR|A0A7D4WPX4|A0A7D4WPX4_CAPHI ------------------------------------------------------------
SP|Q28379|MX1_HORSE              ------------------------------------------------------------
TR|A0A6I9I6Y4|A0A6I9I6Y4_VICPA ------------------------------------------------------------
TR|A0A6P5BYP3|A0A6P5BYP3_BOSIN ------------------------------------------------------------
TR|S9XPZ9|S9XPZ9_CAMFR           ------------------------------------------------------------
TR|A0A384DMX3|A0A384DMX3_URSMA ------------------------------------------------------------
TR|A0A3Q7W4X3|A0A3Q7W4X3_URSAR ------------------------------------------------------------
TR|A0A7E6D288|A0A7E6D288_9CHIR ------------------------------------------------------------
TR|S7NSW7|S7NSW7_MYOBR           --------------------------MLNPGASPW---WS-------------------- 11
TR|A0A1N7THY9|A0A1N7THY9_PIPPI ------------------------------------------------------------
TR|A0A1N7THZ0|A0A1N7THZ0_ROUAE ------------------------------------------------------------
TR|A0A1N7THY7|A0A1N7THY7_HYPMN ------------------------------------------------------------
TR|A0A1N7THZ1|A0A1N7THZ1_STULI ------------------------------------------------------------
TR|B2X026|B2X026_CHICK           ---------------------------------------------MNNPRSNFSSAFGCP 15
TR|A0A172PZ65|A0A172PZ65_ANSCY ---------------------------------------------MYHRSPKFKAVAKRH 15
TR|A0A0Q3PUG8|A0A0Q3PUG8_AMAAE ------------------------------------------------------------
SP|Q4ADG6|MX1_PHOVI              ------------------------------------------------------------
SP|Q4ADG7|MX1_OTABY              ------------------------------------------------------------
TR|A0A2Y9G772|A0A2Y9G772_NEOSC ------------------------------------------------------------
TR|A0A6J2C7I6|A0A6J2C7I6_ZALCA ------------------------------------------------------------
TR|A0A2U3WDI8|A0A2U3WDI8_ODORO ------------------------------------------------------------
TR|A0A1U7SNC8|A0A1U7SNC8_ALLSI --------------------------------MEAQSTTGLFGIS--PSKITFPKVFQTP 26
TR|A0A7M4FFF4|A0A7M4FFF4_CROPO ------------------------------------------------------------
TR|A0A340WTU5|A0A340WTU5_LIPVE ------------------------------------------------------------
TR|A0A6J3RAX5|A0A6J3RAX5_TURTR ------------------------------------------------------------
TR|A0A341CI24|A0A341CI24_NEOAA AAGCDRVEPAHL------------VFRMEAGLGSWELERREI-----PSAVPA----SKL 66
TR|A0A2Y9LTL6|A0A2Y9LTL6_DELLE ------------------------------------------------------------
TR|A0A383ZCI9|A0A383ZCI9_BALAS ------------------------------------------------------------
TR|V9KIB1|V9KIB1_CALMI           ------------------------------------------------------------
TR|A0A674K7E1|A0A674K7E1_TERCA ------------------------------------------------------------
TR|A0A4D9DW37|A0A4D9DW37_9SAUR EENFAVPLPPESEDNFVTSPTPAQTNTVFTGLHSTQSNAVFTGSHPVQSKGAFPGYFPAQ 120
TR|H2DQ54|H2DQ54_XENLA           ------------------------------------------------------------
TR|A0A0F6MUS9|A0A0F6MUS9_ANGJA ------------------------------------------------------------
TR|Q98990|Q98990_SALSA           ------------------------------------------------------------
TR|A0A674EGU5|A0A674EGU5_SALTR ------------------------------------------------------------
SP|Q91192|MX1_ONCMY              ------------------------------------------------------------
TR|A0A286K200|A0A286K200_MYLPI ------------------------------------------------------------
TR|A0A498MHV8|A0A498MHV8_LABRO ------------------------------------------------------------
SP|Q7T2P0|MX1_ICTPU              ------------------------------------------------------------
TR|A0A556TK66|A0A556TK66_BAGYA ------------------------------------------------------------
TR|A0A3G6IID3|A0A3G6IID3_9TELE ------------------------------------------------------------
SP|Q8JH68|MXA_DANRE              ------------------------------------------------------------
TR|A0A672MEY2|A0A672MEY2_SINGR ------------------------------------------------------------
TR|Q7T2M3|Q7T2M3_CARAU           ------------------------------------------------------------
TR|A0A673LWX8|A0A673LWX8_9TELE ------------------------------------------------------------
TR|A0A8B9KF29|A0A8B9KF29_ASTMX ------------------------------------------------------------


SP|P20591|MX1_HUMAN              ------------MVVS----EVDIAKADPAAAS----HPLLLNGDAT------------- 27
TR|K4JBQ0|K4JBQ0_9PRIM           ------------MVVS----EVDIAKADPAAAS----HPLLLNGDAN------------- 27
TR|H2QL18|H2QL18_PANTR           ------------MVVS----EVDIAKADPAAAS----HPLLLNGDAN------------- 27
SP|A1E2I4|MX1_MACMU              ------------MVLS----EVDIVKADPAAAS----QPLLLNGDAD------------- 27
SP|Q5R5G3|MX1_PONAB              ------------MVLS----EVDIAKADPAAAS----HPVLLNGDAN------------- 27
TR|A0A6P8QPW8|A0A6P8QPW8_GEOSA ------------------------------------------------------------
TR|U3EGN4|U3EGN4_CALJA           ------------MVLS----EVGVGKADPAVES----HHQLLNGDAN------------- 27
SP|Q9N0Y3|MX1_CANLF              ------------MVNS--QGKI--TDSNPVP------NHVLLNGL--------------- 23
TR|M3W8X0|M3W8X0_FELCA           -----------------------MVNS-KGKIT----DSDP---VSDH-----L------ 18
TR|A0A667GNL8|A0A667GNL8_LYNCA -----------------------MVNS-KGKIT----DSDP---VSDH-----L------ 18
TR|A0A6P6GZN4|A0A6P6GZN4_PUMCO -----------------------MVNS-KGKIT----DSDP---VSDH-----L------ 18
TR|A0A6J1XXY6|A0A6J1XXY6_ACIJB -----------------------MVNS-KGKIT----DSDP---VSNH-----L------ 18
TR|A0A6P4TS13|A0A6P4TS13_PANPR -------------------SGIAAIWN-PGRAS----HGKTIGGVGEELRVSIQ------ 56
TR|A0A3Q7SCG7|A0A3Q7SCG7_VULVU ---LFLRGGP--PRRV--LGGIEEEWYRPLP------ASPLLRGRFGF------------ 56
TR|I3MJX6|I3MJX6_ICTTR           ------------MSPP--PSEVN-CANPAAVES----NHLFPNGDAS------------- 28
TR|G9KBY2|G9KBY2_MUSPF           ------------MVNS--TGKITDSD--P---------GLILNGLAD------------- 22
TR|A0A2Y9KR88|A0A2Y9KR88_ENHLU ------------MVNS--TEKITDSD--P---------GLILNGLAD------------- 22
SP|P09922|MX1B_MOUSE             ------------------------------------------------------------
```

```
SP|P18588|MX1_RAT              -------------------------MKERTSAC----RHGTPQKH--------------- 16
TR|A0A6P6D2J7|A0A6P6D2J7_PTEVA ------MSLPT-VKRS--PLKGKVKNHNSASAS----SPRVPDA---------------- 31
TR|A0A6J0DKA3|A0A6J0DKA3_PERMB ------STWKN-MRM---SSSWKVNKKELTSAS----SHLNPPEHPD------------- 65
TR|G5C5C9|G5C5C9_HETGA         ------------MSHS----APNAIKRCVAATS----KHLSLNGDAN------------- 27
TR|A0A3Q0CWV5|A0A3Q0CWV5_MESAU ---VN--------------------QKELTAAA----YPPKPHK---------------- 34
TR|A0A6I9K5X3|A0A6I9K5X3_CHRAS -------------MTD--DKERTSTSNSAATPN----NHSLLNGDVD------------- 28
SP|P79135|MX1_BOVIN            ------------MVHS----DLGIEELDSP-ES-------SLNGSED------------- 23
TR|A0A6P3H2K4|A0A6P3H2K4_BISBI ------------MVHS----DLGIKELDSP-ES-------SLNGSED------------- 23
TR|L8IPR7|L8IPR7_9CETA         --------------------MLGVMGGRRRRGR-------RRMRCLD------------- 20
SP|P27594|MX1_PIG              ------------MVYS----SCESKEPDSVSAS----NHLLLNGNDE------------- 27
TR|A0A1U7U9K3|A0A1U7U9K3_CARSF ------------------------------------------------------------
SP|P33237|MX1_SHEEP            ------------MVLS----DLDIKEPDSP-ES-------GLNGSDD------------- 23
TR|A0A7D4WPX4|A0A7D4WPX4_CAPHI ------------MVLS----DLDIKEPDSP-ES-------NLNGSDD------------- 23
SP|Q28379|MX1_HORSE            ------------MVHS----EAKMTRPDSASAS----KQQLLNGNAD------------- 27
TR|A0A6I9I6Y4|A0A6I9I6Y4_VICPA ------------MVYS----DSENKEFDSASAS----NHVSLNGNDD------------- 27
TR|A0A6P5BYP3|A0A6P5BYP3_BOSIN --------------------MLGVMGGRRRRGR-------RRMRCLD------------- 20
TR|S9XPZ9|S9XPZ9_CAMFR         --------------MG----EQ------RGLLL----QHILLTAHPETVSLF-------- 24
TR|A0A384DMX3|A0A384DMX3_URSMA ------------MVKS--KGKIT--DSDPGS------NQLLLNGLAD------------- 25
TR|A0A3Q7W4X3|A0A3Q7W4X3_URSAR ------------MVKS--KGKIT--DSDPGS------NQLLLNGLAD------------- 25
TR|A0A7E6D288|A0A7E6D288_9CHIR ------------MGH----SGLQTKNPSSASGS----NHELSNENAN------------- 27
TR|S7NSW7|S7NSW7_MYOBR         ---VSSHRGN--AAQS--EARLTTGECSSGGGS----L-SRFHGNAKK------------ 47
TR|A0A1N7THY9|A0A1N7THY9_PIPPI ---------------------METATPDSASAS----SQPLPNGDAV------------- 22
TR|A0A1N7THZ0|A0A1N7THZ0_ROUAE ------MGVPQ----------VKVESHGSASAP----IPQLPNA---------------- 24
TR|A0A1N7THY7|A0A1N7THY7_HYPMN ------MGVPQ----------VKVQGHSSDLAS----APQLPNA---------------- 24
TR|A0A1N7THZ1|A0A1N7THZ1_STULI ------------MVR----SGLQTRNPNSS-TS----NREQSNENAD------------- 26
TR|B2X026|B2X026_CHICK         --------IQIPKQNSNVPPSLPVPVGVFG--------VPLPLGCSNQMAF--CAPELTD 57
TR|A0A172PZ65|A0A172PZ65_ANSCY --------SPILMKDGFPSLSLPVCVDASG--------VPLPPDLDDQIDF--PVTEQTT 57
TR|A0A0Q3PUG8|A0A0Q3PUG8_AMAAE ------------MKGVFASHSSQPSFKAFA--------VPLPPELEDEDM---MMDNVME 37
SP|Q4ADG6|MX1_PHOVI            ------------MVNS--KGEIT--DSDPGS------NHLLLNGLPD------------- 25
SP|Q4ADG7|MX1_OTABY            ------------MVNS--KGKIT--DSDPGS------SHLLLNGLAD------------- 25
TR|A0A2Y9G772|A0A2Y9G772_NEOSC ------------MVNS--KGKIT--DSDPGS------NHLLLNGLPD------------- 25
TR|A0A6J2C7I6|A0A6J2C7I6_ZALCA ------------MVNS--KGKIT--DSDPGS------SHLLLNGLAD------------- 25
TR|A0A2U3WDI8|A0A2U3WDI8_ODORO ------------MFNS--KGKIT--DSDPGS------SHLLLNGLAD------------- 25
TR|A0A1U7SNC8|A0A1U7SNC8_ALLSI SNMGFAAQSPSQLQAA---PTLPLSRSPIPVVSP-LSTTTPPPERNKSIQQGLATEHHIE 82
TR|A0A7M4FFF4|A0A7M4FFF4_CROPO ------------------------------------------------------------
TR|A0A340WTU5|A0A340WTU5_LIPVE ------------MVHS----DLKVKEPDSPSAS----SHLLLNGNDD------------- 27
TR|A0A6J3RAX5|A0A6J3RAX5_TURTR ------------MVHS----DLKIKEPDSPSAS----SHLLLNGNDD------------- 27
TR|A0A341CI24|A0A341CI24_NEOAA PAILSADTGKK-VVHS----DLKIKEPDSPSTS----SHLLLNGNDD------------- 104
TR|A0A2Y9LTL6|A0A2Y9LTL6_DELLE ------------MVHS----DLKIKEPDSPSAS----SHLLLNGNDD------------- 27
TR|A0A383ZCI9|A0A383ZCI9_BALAS ------------MVHS----DLEIKESDSPPAS----SHLLLNGNDD------------- 27
TR|V9KIB1|V9KIB1_CALMI         ------------------------------------------------------------
TR|A0A674K7E1|A0A674K7E1_TERCA ----------RDMRRT---SDVRVTVSHSSTLLP-SLRIPPPPP---------------- 30
TR|A0A4D9DW37|A0A4D9DW37_9SAUR TKAVFPGSLPTKSKEAL--TRLQPTESKGATAVPHPCMQSAPPEWNEMIQREIAAAQWAE 178
TR|H2DQ54|H2DQ54_XENLA         ------------------------------------------------------------
TR|A0A0F6MUS9|A0A0F6MUS9_ANGJA ------------------------------------------------------------
TR|Q98990|Q98990_SALSA         ------------------------------------------------------------
TR|A0A674EGU5|A0A674EGU5_SALTR ------------------------------------------------------------
SP|Q91192|MX1_ONCMY            ------------------------------------------------------------
TR|A0A286K200|A0A286K200_MYLPI ------------------------------------------------------------
TR|A0A498MHV8|A0A498MHV8_LABRO ------------------------------------------------------------
SP|Q7T2P0|MX1_ICTPU            ------------------------------------------------------------
TR|A0A556TK66|A0A556TK66_BAGYA ------------------------------------------------------------
TR|A0A3G6IID3|A0A3G6IID3_9TELE ------------------------------------------------------------
SP|Q8JH68|MXA_DANRE            ------------------------------------------------------------
TR|A0A672MEY2|A0A672MEY2_SINGR ------------------------------------------------------------
TR|Q7T2M3|Q7T2M3_CARAU         ------------------------------------------------------------
TR|A0A673LWX8|A0A673LWX8_9TELE ------------------------------------------------------------
TR|A0A8B9KF29|A0A8B9KF29_ASTMX ------------------------------------------------------------


SP|P20591|MX1_HUMAN            -----------------VAQKNPGSVAENNLCSQYEEKVRPCIDLIDSLRALGVEQDLAL 70
TR|K4JBQ0|K4JBQ0_9PRIM         -----------------VAQKNPGLVAENNLCSQYEEKVRPCIDLIDSLRALGVEQDLAL 70
TR|H2QL18|H2QL18_PANTR         -----------------VAQKNPGSVAENNLCSQYEEKVRPCIDLIDSLRALGVEQDLAL 70
SP|A1E2I4|MX1_MACMU            -----------------VAQKSPGSVAENNLCSQYEEKVRPCIDLIDSLRALGVEQDLAL 70
SP|Q5R5G3|MX1_PONAB            -----------------VAQKNLGSVAENNLCSQYEEKVRPCIDLIDSLRALGVEQDLAL 70
TR|A0A6P8QPW8|A0A6P8QPW8_GEOSA --------------------------MENYLNNAYEQKIRPCIDLIDGLRSLGIQKELAL 34
TR|U3EGN4|U3EGN4_CALJA         -----------------VAKKNPASVAENNLYSQYEEKVRPCIDLIDSLRALGVEQDLAL 70
SP|Q9N0Y3|MX1_CANLF            -----------------TDKAEKNQGIGNSLCSQYEEKVRPCIDLIDSLRALGVEQDLAL 66
TR|M3W8X0|M3W8X0_FELCA         ----------LLNGLADKAEKNQETGLKKNLCSQYEEKVRPCIDLIDSLRALGVEQDLAL 68
TR|A0A667GNL8|A0A667GNL8_LYNCA ----------LLNGLADKTEKNQEMGLKKNLCSQYEEKVRPCIDLIDSLRALGVEQDLAL 68
TR|A0A6P6GZN4|A0A6P6GZN4_PUMCO ----------LLNGLADKAEKNQEMGLKKSLCSQYEEKVRPCIDLIDSLRALGVEQDLAL 68
TR|A0A6J1XXY6|A0A6J1XXY6_ACIJB ----------LLNGLADKAEKNQEMGLEKSLCSQYEEKVRPCIDLIDSLRALGVEQDLAL 68
TR|A0A6P4TS13|A0A6P4TS13_PANPR ----------QVNVASSREKEGVLEGLEKSLCSQYEEKVRPCIDLIDSLRALGVEQDLAL 106
TR|A0A3Q7SCG7|A0A3Q7SCG7_VULVU -----------VTF-PTVGERKPGERTGNSLCSQYEEKVRPCIDLIDSLRALGVEQDLAL 104
TR|I3MJX6|I3MJX6_ICTTR         -----------------TAERNEALGSEFNLCRQYEEKVRPCIDLIDSLRALGVERDLAL 71
TR|G9KBY2|G9KBY2_MUSPF         -----------------KAEKNPDMEPENSLCNHYEEKVRPCIDLIDSLRALGVEQDLAL 65
TR|A0A2Y9KR88|A0A2Y9KR88_ENHLU -----------------KAEKNPDMEPKNSLCNHYEEKVRPCIDLIDSLRALGVEQDLAL 65
SP|P09922|MX1B_MOUSE           ------------------------MDSVNNLCRHYEEKVRPCIDLIDTLRALGVEQDLAL 36
SP|P18588|MX1_RAT              ---------------PDTSEESQAMESVDNLCSQYEEKVRPCIDLIDSLRALGVEQDLAL 61
TR|A0A6P6D2J7|A0A6P6D2J7_PTEVA ----------------KVSEENLKKDPESSLCSYYEEKIRPCIDLIDSLRALGVEQDLAL 75
TR|A0A6J0DKA3|A0A6J0DKA3_PERMB -----------------ASKDSQAMESLNNLCRQYEEKVRPCIDLIDSLQALGVEQDLAL 108
TR|G5C5C9|G5C5C9_HETGA         -----------------KGEQAQPMVPENNLCRQYKEKVQPCIDLIDSLRALGVEQDLAL 70
TR|A0A3Q0CWV5|A0A3Q0CWV5_MESAU -----------------HPEDSQSMVSVNNLCRQYEEKVRPCIDLIDSLRALGVEQDLAL 77
TR|A0A6I9K5X3|A0A6I9K5X3_CHRAS -----------------MSKSSPWLVAENTLYSQYEEKVRPCIDLIDSLRALGVEQDLAL 71
SP|P79135|MX1_BOVIN            -----------------M-------ESKSNLYSQYEEKVRPCIDLIDSLRSLGVEQDLAL 59
TR|A0A6P3H2K4|A0A6P3H2K4_BISBI -----------------MVRE-HETESKSNLYSQYEEKVRPCIDLIDSLRSLGVEQDLAL 65
TR|L8IPR7|L8IPR7_9CETA         -----------------GITD-SMDEPKSNLYSQYEEKVRPCIDLIDSLRSLGVEQDLAL 62
SP|P27594|MX1_PIG              -----------------LVEKSHKTGPENNLYSQYEEKVRPCIDLIDSLRALGVEQDLAL 70
```

```
TR|A0A1U7U9K3|A0A1U7U9K3_CARSF -------------------------EPENNLCNQYEEKVRPCIDLIDSLRALGVEQDLAL 35
SP|P33237|MX1_SHEEP            -----------------MVRE-HETESKGNLYSQYEEKVRPCIDLIDSLRALGVEQDLAL 65
TR|A0A7D4WPX4|A0A7D4WPX4_CAPHI -----------------MVRE-HEMESKGNLYSQYEEKVRPCIDLIDSLRALGVEQDLAL 65
SP|Q28379|MX1_HORSE            -----------------IQETNQKRSIEKNLCSQYEEKVRPCIDLIDSLRALGVEQDLAL 70
TR|A0A6I9I6Y4|A0A6I9I6Y4_VICPA -----------------MVEKNQETGPENNLYSQYEEKIRPCIDLIDSLRALGVEQDLAL 70
TR|A0A6P5BYP3|A0A6P5BYP3_BOSIN -----------------GITD-SMDESKSNLYSQYEEKVRPCIDLIDSLRSLGVEQDLAL 62
TR|S9XPZ9|S9XPZ9_CAMFR         --------------FVTRKYLRPRRGPENNLYSQYEEKIRPCIDLVDSLRALGVEQDLAL 70
TR|A0A384DMX3|A0A384DMX3_URSMA -----------------EEEKNQDTEPKLSLCSQYEEKVRPCIDLIDSLRALGVEQDLAL 68
TR|A0A3Q7W4X3|A0A3Q7W4X3_URSAR -----------------EEEKNQDTEPKLSLCSQYEEKVRPCIDLIDSLRALGVEQDLAL 68
TR|A0A7E6D288|A0A7E6D288_9CHIR -----------------EPKENWETKLENRLCRQYEEKVRPCIDLIDSLRALGVEQDLAL 70
TR|S7NSW7|S7NSW7_MYOBR         -----------------QQVECSEQTLENSLCSQYEEKVRPCIDLIDSLRALGVEQDLAL 90
TR|A0A1N7THY9|A0A1N7THY9_PIPPI -----------------VPEVNMETMLETSLCSQYEEKVRPCIDLIDSLRALGVEQDLAL 65
TR|A0A1N7THZ0|A0A1N7THZ0_ROUAE ----------------EVPEKNLDKGDEINLCSYYEEKVRPCIDLIDSLRALGVEQDLAL 68
TR|A0A1N7THY7|A0A1N7THY7_HYPMN ----------------EVPEKNLKKGEETDLCSHYEEKVRPCIDLVDSLRALGVEQDLAL 68
TR|A0A1N7THZ1|A0A1N7THZ1_STULI -----------------EPKKNRD-TLEHHLCSQYEEKVRPCIDLVDSLRALGVEQDLAL 68
TR|B2X026|B2X026_CHICK         RKPEHEQ---KVSKRLNDREEDKDEAAACSLDNQYDRKIRPCIDLVDSLRKLDIGNDLML 114
TR|A0A172PZ65|A0A172PZ65_ANSCY RESQQEQ---KVSMKLHEGEQDM-QAAEHTLYNQYEEKIRPCIDLIDSLRALGIEKDLSL 113
TR|A0A0Q3PUG8|A0A0Q3PUG8_AMAAE RKPNYDQ---KEXXKQYYGK--EDKAAEHTLYNQYEKKIRPCINLXDSLRALGIEKDMAL 92
SP|Q4ADG6|MX1_PHOVI            -----------------KAGKNQDTEPENSLCSQYEEKVRPCIDLIDSLRALGVEQDLAL 68
SP|Q4ADG7|MX1_OTABY            -----------------KAGKNQDTEPENSLCSQYEEKVRPCIDLIDSLRALGVEQDLAL 68
TR|A0A2Y9G772|A0A2Y9G772_NEOSC -----------------KAGKNQDTEPENSLCSQYEEKVRPCIDLIDSLRALGVEQDLAL 68
TR|A0A6J2C7I6|A0A6J2C7I6_ZALCA -----------------KAGKNQDTEPENSLCSQYEEKVRPCIDLIDSLRALGVEQDLAL 68
TR|A0A2U3WDI8|A0A2U3WDI8_ODORO -----------------KAGKNQDTEPENSLCSQYEEKVRPCIDLIDSLRALGVEQDLAL 68
TR|A0A1U7SNC8|A0A1U7SNC8_ALLSI SEKNRHQREKTQQGEIYASEEKNKQKEENTLYNQYEMRIRPCIDLVDSLRALGIEKDLAL 142
TR|A0A7M4FFF4|A0A7M4FFF4_CROPO ----------MQQGEIYPSEEMITQKAENTLYNQYEMRIRPCIDFIDALRALGIEKNLAL 50
TR|A0A340WTU5|A0A340WTU5_LIPVE -----------------LVGKNEETGSENSLCNQYEEKVRPCIDLIDSLRXLGVEQGPAP 70
TR|A0A6J3RAX5|A0A6J3RAX5_TURTR -----------------LVEKNEETGSENNLYNQYEEKVRPCIDLIDSLQALGVEQGLAL 70
TR|A0A341CI24|A0A341CI24_NEOAA -----------------LVGKNEETGSENNLYNQYEEKVRPCIDLIDSLQALGVEQGLAL 147
TR|A0A2Y9LTL6|A0A2Y9LTL6_DELLE -----------------LVGKNEETGSENNLYNQYEEKVRPCSDLIDSLQALGVEQGLAL 70
TR|A0A383ZCI9|A0A383ZCI9_BALAS -----------------LVGK-KETGSESNLYSQYEEKVRPCIDLIDSLRALGVEQDLAL 69
TR|V9KIB1|V9KIB1_CALMI         --------------------------MDVALYSHYEEEVRPCIDLIDHLRSLGLEKDLSL 34
TR|A0A674K7E1|A0A674K7E1_TERCA --------------GVQTYTIFFYQKAENTLCSQYEEKVRPCIDLIDSLRALGVEKDLAL 76
TR|A0A4D9DW37|A0A4D9DW37_9SAUR -------------SEKYQQEKMKQQNTEHTLYNQYEEKIRPCIDLIDSLRALGVEKDLAL 225
TR|H2DQ54|H2DQ54_XENLA         -------------------------MENILSSQYEEKIRPCIDLIDSLRALGVDKDLGL 34
TR|A0A0F6MUS9|A0A0F6MUS9_ANGJA -----------MAETSLGMPEATEAKLSSTLNQHYEEKVRPCIDLIDSLRSLGVEKDLAL 49
TR|Q98990|Q98990_SALSA         --------------------------MNNTLNQHYEEKVRPCIDLIDSLRSLGVEKDLAL 34
TR|A0A674EGU5|A0A674EGU5_SALTR --------------------------MNNTLNQHYEEKVRPCIDLIDSLRSLGVEKDLAL 34
SP|Q91192|MX1_ONCMY            --------------------------MNNTLNQHYEEKVRPCIDLIDSLRSLGVEKDLAL 34
TR|A0A286K200|A0A286K200_MYLPI -----------------------MEKMSYTFSQHYEEKIRPCIDTIDNLRSLGVEKDLAL 37
TR|A0A498MHV8|A0A498MHV8_LABRO -----------------------MEKMSYTFSQQYEEKIRPCIDTIDSLRSLGVEKDLAL 37
SP|Q7T2P0|MX1_ICTPU            --------------------------MSASLSEQYEEKVRPCIDLIDSLRALGVEKDLAL 34
TR|A0A556TK66|A0A556TK66_BAGYA --------------------------MSASLSEQYEEKVRPCIDLIDSLRALGVEKDLAL 34
TR|A0A3G6IID3|A0A3G6IID3_9TELE -----------------------MEKMSYTFSQHYEEKIRPCIDTIDNLRSLGVEKDLAL 37
SP|Q8JH68|MXA_DANRE            -----------------------MEKLSYTFSQQYEEKIRPCIDTIDNLRSLGVEKDLAL 37
TR|A0A672MEY2|A0A672MEY2_SINGR -----------------------KIKLTYTFHNHFIESVRPLIELIDSLKLIGLDEDIGL 37
TR|Q7T2M3|Q7T2M3_CARAU         -----------------------MEKMSYTFSQHYEEKIRPCIDTIDNLRSLGVEKDLAL 37
TR|A0A673LWX8|A0A673LWX8_9TELE -----------------------MEKMSYTFSQQYEEKIRPCIDTIDNLRSLGVEKDLAL 37
TR|A0A8B9KF29|A0A8B9KF29_ASTMX --------------------------MGSTLNEQYEEKVRPCIDLIDSLRSQGVEKDLAL 34
                                                          :    :    ::*  :   * *:   .: .

SP|P20591|MX1_HUMAN            PAIAVIGDQSSGKSSVLEALSGVALPRGSGIVTRCPLVLKLKKLVNE-DKWRGKVSYQDY 129
TR|K4JBQ0|K4JBQ0_9PRIM         PAIAVIGDQSSGKSSVLEALSGVALPRGSGIVTRCPLVLKLKKLVNE-DKWRGKVSYQDY 129
TR|H2QL18|H2QL18_PANTR         PAIAVIGDQSSGKSSVLEALSGVALPRGSGIVTRCPLVLKLKKLVNE-DKWRGKVSYQDY 129
SP|A1E2I4|MX1_MACMU            PAIAVIGDQSSGKSSVLEALSGVALPRGSGIVTRCPLVLKLKKLVNE-DEWRGKVSYQDY 129
SP|Q5R5G3|MX1_PONAB            PAIAVIGDQSSGKSSVLEALSGVALPRGSGIVTRCPLVLKLKKLVNE-DKWRGKVSYQDY 129
TR|A0A6P8QPW8|A0A6P8QPW8_GEOSA PAITVIGDQGSGKSSVLEALSGVALPRGTGIVTRCPLELKLKKAKNG-QKWSGKISYEDV 93
TR|U3EGN4|U3EGN4_CALJA         PAIAVIGDQSSGKSSVLEALSGVALPRGSGIVTRCPLVLKLKKLVNE-EEWKGKVSYQDF 129
SP|Q9N0Y3|MX1_CANLF            PAIAVIGDQSSGKSSVLEALSGVALPRGSGIVTRCPLVLKLKKLINE-DEWRGKVSYQDT 125
TR|M3W8X0|M3W8X0_FELCA         PAIAVIGDQSSGKSSVLEALSGVALPRGSGIVTRCPLVLKLKKLMNE-DEWRGKVSYQDF 127
TR|A0A667GNL8|A0A667GNL8_LYNCA PAIAVIGDQSSGKSSVLEALSGVALPRGSGIVTRCPLVLKLKKLMNE-DEWRGKVSYQDF 127
TR|A0A6P6GZN4|A0A6P6GZN4_PUMCO PAIAVIGDQSSGKSSVLEALSGVALPRGSGIVTRCPLVLKLKKLMNE-DEWRGKVSYQDF 127
TR|A0A6J1XXY6|A0A6J1XXY6_ACIJB PAIAVIGDQSSGKSSVLEALSGVALPRGSGIVTRCPLVLKLKKLMNE-DEWRGKVSYQDF 127
TR|A0A6P4TS13|A0A6P4TS13_PANPR PAIAVIGDQSSGKSSVLEALSGVALPRGSGIVTRCPLVLKLKKLMNE-DEWRGKVSYQDF 165
TR|A0A3Q7SCG7|A0A3Q7SCG7_VULVU PAIAVIGDQSSGKSSVLEALSGVALPRGSGIVTRCPLVLKLKKLINE-DEWRGKVSYQDT 163
TR|I3MJX6|I3MJX6_ICTTR         PAIAVIGDQSSGKSSVLEALSGVALPRGSGIVTRCPLVLRMKKLGQE-DEWRGKVSYLDM 130
TR|G9KBY2|G9KBY2_MUSPF         PAIAVIGDQSSGKSSVLEALSGVALPRGSGIVTRCPLVLKLKKVTNQ-DEWRGKVSYQDF 124
TR|A0A2Y9KR88|A0A2Y9KR88_ENHLU PAIAVIGDQSSGKSSVLEALSGVALPRGSGIVTRCPLVLKLKKVTNQ-DEWRGKVSYQDF 124
SP|P09922|MX1B_MOUSE           PAIAVIGDQSSGKSSVLEALSGVALPRGSGIVTRCPLVLKLRKLKEG-EEWRGKVSYDDI 95
SP|P18588|MX1_RAT              PAIAVIGDQSSGKSSVLEALSGVALPRGSGIVTRCPLVLKLKQLKQG-EKWSGKVIYKDT 120
TR|A0A6P6D2J7|A0A6P6D2J7_PTEVA PAIAVIGDQSSGKSSVLEALSGVALPRGSGIVTRCPLVLKLRKLRDG-DEWKGKVSYQDQ 134
TR|A0A6J0DKA3|A0A6J0DKA3_PERMB PAMAVIGDQSSGKSSVLEALSGVALPRA-------------------------------- 136
TR|G5C5C9|G5C5C9_HETGA         PAIAVIGDQSSGKSSVLEALSGVALPRGSGIVTRCPLVLKLKKLDQE-EGWRGKLSYLQV 129
TR|A0A3Q0CWV5|A0A3Q0CWV5_MESAU PAIACI----------------------LGIVTRCPLVLKLKQLGQG-EEWRGKVTYKDT 114
TR|A0A6I9K5X3|A0A6I9K5X3_CHRAS PAIAVIGDQSSGKSSVLEALSGVALPRGSGIVTRCPLVLKLKKLLHG-DEWKGKVSYKDL 130
SP|P79135|MX1_BOVIN            PAIAVIGDQSSGKSSVLEALSGVALPRGSGIVTRCPLVLRLKKLGN-EDEWKGKVSFLDK 118
TR|A0A6P3H2K4|A0A6P3H2K4_BISBI PAIAVIGDQSSGKSSVLEALSGVALPRGSGIVTRCPLVLRLKKLGN-EDEWKGKVSFLDK 124
TR|L8IPR7|L8IPR7_9CETA         PAIAVIGDQSSGKSSVLEALSGVALPRGSGIVTRCPLVLRLKKLGN-EDEWKGKVSFLDK 121
SP|P27594|MX1_PIG              PAIAVIGDQSSGKSSVLEALSGVALPRGSGIVTRCPLVLKLKKLVNEEDEWKGKVSYRDS 130
TR|A0A1U7U9K3|A0A1U7U9K3_CARSF PAIAVIGDQSSGKSSVLEALSGVALPRGSGIVTRCPLVLKLRKLRHG-EEWKGKVSYQHI 94
SP|P33237|MX1_SHEEP            PAIAVIGDQSSGKSSVLEALSGVALPRGSGIVTRCPLVLRLKKLEK-EGEWKGKVSFLDR 124
TR|A0A7D4WPX4|A0A7D4WPX4_CAPHI PAIAVIGDQSSGKSSVLEALSGVALPRGSGIVTRCPLVLRLKKLEK-EGEWKGKVSFLDR 124
SP|Q28379|MX1_HORSE            PAIAVIGDQSSGKSSVLEALSGVALPRGSGIVTRCPLVLKLKRLVKE-DEWKGKVSYRDI 129
TR|A0A6I9I6Y4|A0A6I9I6Y4_VICPA PAIAVIGDQSSGKSSVLEALSGVALPRGSGIVTRCPLVLKLKKLVN-EDGWRGKVSFRDY 129
TR|A0A6P5BYP3|A0A6P5BYP3_BOSIN PAIAVIGDQSSGKSSVLEALSGVALPRGSGIVTRCPLVLRLKKLGN-EDEWKGKVSFLDK 121
TR|S9XPZ9|S9XPZ9_CAMFR         PAIAVIGDQSSGKSSVLEALSGVALPRGSGIVTRCPLVLKLKKLVN-EDGWRGKVSFRDY 129
TR|A0A384DMX3|A0A384DMX3_URSMA PAIAVIGDQSSGKSSVLEALSGVALPRGNGIVTRCPLVLKLKKLLNE-DEWRGKVSYQDF 127
TR|A0A3Q7W4X3|A0A3Q7W4X3_URSAR PAIAVIGDQSSGKSSVLEALSGVALPRGNGIVTRCPLVLKLKKLLNE-DEWRGKVSYQDF 127
TR|A0A7E6D288|A0A7E6D288_9CHIR PAIAVIGDQSSGKSSVLEALSGVALPRGSGIVTRCPLVLKLKKLKSQ-EKWTGKISYKDI 129
```

```
TR|S7NSW7|S7NSW7_MYOBR         PAIAVIGDQSSGKSSVLEALSGVSLPRGSGIVTRCPLVLKLRKLRHD-DEWKGKVTYRDL 149
TR|A0A1N7THY9|A0A1N7THY9_PIPPI PAIAVIGDQSSGKSSVLEALSGVALPRGSGIVTRCPLVLKLRKLRHDDDEWKGKVTYRDM 125
TR|A0A1N7THZ0|A0A1N7THZ0_ROUAE PAIAVIGDQSSGKSSVLEALSGVALPRGSGIVTRCPLVLKLRKVTDG-EGWKGKVSYQDV 127
TR|A0A1N7THY7|A0A1N7THY7_HYPMN PAIAVIGDQSSGKSSVLEALSGVALPRGSGIVTRCPLVLKLRQARDG-GAWKGKVSYQDE 127
TR|A0A1N7THZ1|A0A1N7THZ1_STULI PAIAVIGDQSSGKSSVLEALSGVALPRGSGIVTRCPLVLKLKNLKDE-EKWTGKISYKDI 127
TR|B2X026|B2X026_CHICK         PAIAVIGDRNSGKSSVLEALSGVALPRDKGVITRCPLELKLKKMTAP-QEWKGVIYYRNT 173
TR|A0A172PZ65|A0A172PZ65_ANSCY PAIAVIGDQSSGKSSILEALSGVSLPRGNGIVTRCPLELKLKKIPTS-QQWKGKISYRNI 172
TR|A0A0Q3PUG8|A0A0Q3PUG8_AMAAE PAIAVIGDQSSGKSSVLEALSGIALPRGNGIVTRCPLELKLKRTPAT-QEWKGKIHYRNF 151
SP|Q4ADG6|MX1_PHOVI            PAIAVIGDQSSGKSSVLEALSGVALPRGSGIVTRCPLVLKLKKLLNE-DEWRGKVSYQDF 127
SP|Q4ADG7|MX1_OTABY            PAIAVIGDQSSGKSSVLEALSGVALPRGSGIVTRCPLVLKLKKLLNK-DEWRGKVSYQDF 127
TR|A0A2Y9G772|A0A2Y9G772_NEOSC PAIAVIGDQSSGKSSVLEALSGVALPRGSGIVTRCPLVLKLKKLLNE-DEWRGKVSYQDF 127
TR|A0A6J2C7I6|A0A6J2C7I6_ZALCA PAIAVIGDQSSGKSSVLEALSGVALPRGSGIVTRCPLVLKLKKLLNK-DEWRGKVSYQDF 127
TR|A0A2U3WDI8|A0A2U3WDI8_ODORO PAIAVIGDQSSGKSSVLEALSGVALPRGSGIVTRCPLVLKLKKLLNE-DEWRGKVSYQDF 127
TR|A0A1U7SNC8|A0A1U7SNC8_ALLSI PAIAVIGDQSSGKSSVLEALSGVALPRGSGIVTRCPLELKLRKTP-E-QVWKGKISYRGM 200
TR|A0A7M4FFF4|A0A7M4FFF4_CROPO PAIAVIGDQSSGKSSVLEALSGVALPRGSGIVTRCPLELKLRKAP-E-QVWKGSIKYQGM 108
TR|A0A340WTU5|A0A340WTU5_LIPVE PAIAVTGDXSSGKSSVLEALSGVALPRGSGIVTRCPLVLKLKKLVN-EDEWKGKVSFRDK 129
TR|A0A6J3RAX5|A0A6J3RAX5_TURTR PTIAVTGDQSSGKSSVLEALSGFALPRGSGIVTRCPLVLKLKTLVN-EDEWKGKVSVRDK 129
TR|A0A341CI24|A0A341CI24_NEOAA PAIAVTGDQSSGKSSVLEALSGFALPRGSGIVTRCPLVLKLKTRVN-EDERKGKVSFRDK 206
TR|A0A2Y9LTL6|A0A2Y9LTL6_DELLE PAIAVTGDQSSGKSSVLEALSGFALPRGSGIVTRCPLVLKLKTCVN-EDEWRGKVSFRDK 129
TR|A0A383ZCI9|A0A383ZCI9_BALAS PAIAVIGDQSSGKSSVLEALSGVALPRGSGIVTRCPLVLKLKKLVN-EDEWKGKVSFRDK 128
TR|V9KIB1|V9KIB1_CALMI         PAIAVIGDQSSGKSSVLEALSGVSLPRGTGIVTRCPLELKLKNVKEK-DVWTGFITYNNH 93
TR|A0A674K7E1|A0A674K7E1_TERCA PAIAVIGDQSSGKSSVLEALSGVALPRGSGIVTRCPLELKLKKVHYT-QEWKGKITYLAI 135
TR|A0A4D9DW37|A0A4D9DW37_9SAUR PAIAVIGDQSSGKSSVLEALSGVALPRGSGITTRCPLVLRLKKLAPQ-QKWKGKISYRDI 284
TR|H2DQ54|H2DQ54_XENLA         PSIAVIGDQSSGKSSVLEALSGVTLPRGSGIVTRCPLELKLKKATKS-TTWSGKISYRDH 93
TR|A0A0F6MUS9|A0A0F6MUS9_ANGJA PAIAVIGDQSSGKSSVLEALSGVALPRGSGIVTRCPLELKMKRLKED-DAWHGKISYN-- 106
TR|Q98990|Q98990_SALSA         PAIAVIGDQSSGKSSVLEALSGVALPRGSGIVTRCPLELKMKRKKEG-EEWHGKISYQ-- 91
TR|A0A674EGU5|A0A674EGU5_SALTR PAIAVIGDQSSGKSSVLEALSGVALPRGSGIVTRCPLELKMKRKKEG-EEWHGKISYQ-- 91
SP|Q91192|MX1_ONCMY            PAIAVIGDQSSGKSSVLEALSGVALPRGSGIVTRCPLELKMKRKKEG-EEWHGKISYQ-- 91
TR|A0A286K200|A0A286K200_MYLPI PAIAVIGDRSSGKSSVLEALSGVALPRGSGIVTRCPLELKMIRTKGE-EKWHAKISYQNH 96
TR|A0A498MHV8|A0A498MHV8_LABRO PAIAVIGDQSSGKSSVLEALSGVALPRGSGIVTRCPLELKMIRTKDE-EKWHGKISYMGK 96
SP|Q7T2P0|MX1_ICTPU            PAIAVIGDQSSGKSSVLEALSGVALPRGSGIVTRCPLELKMKRSREE-DFWHGKIKYKKD 93
TR|A0A556TK66|A0A556TK66_BAGYA PAIAVIGDQSSGKSSVLEALSGVALPRGSGIVTRCPLELKMKRSQEK-EIWHGKIKFKKG 93
TR|A0A3G6IID3|A0A3G6IID3_9TELE PAIAVIGDQSSGKSSVLEALSGVALPRGSGIVTRCPLELKMIRTKGE-EKWHAKISYQNH 96
SP|Q8JH68|MXA_DANRE            PAIAVIGDQSSGKSSVLEALSGVPLPRGSGIVTRCPLELKMIRTKDQ-DRWHGRISYKTC 96
TR|A0A672MEY2|A0A672MEY2_SINGR PSIAVVGDQSSGKSSVLEALSGVALPRGSGIVTRCPLELKLQKLKN--GPWSGIISYSGH 95
TR|Q7T2M3|Q7T2M3_CARAU         PAIAVIGDQSSGKSSVLEALSGVALPRGSGIVTRCPLELKMIRTKGE-GKWHAKISYQDY 96
TR|A0A673LWX8|A0A673LWX8_9TELE PAIAVIGDQSSGKSSVLEALSGVALPRGSGIVTRCPLELKMIRTKDE-DKWHGRISYNKK 96
TR|A0A8B9KF29|A0A8B9KF29_ASTMX PAIAVIGDQSSGKSSVLEALSGVALPRGSGIVTRCPLELKMKRCREE-DSWHGKISYD-- 91
                               *:::

SP|P20591|MX1_HUMAN            -----EIEISDASEVEKEI-NKAQNAIAGEGMGISHELITLEISSRDVPDLTLIDLPGIT 183
TR|K4JBQ0|K4JBQ0_9PRIM         -----EIEISDASEVEKEI-NKAQNTIAGEGMGISHELITLEVSSRDVPDLTLIDLPGIT 183
TR|H2QL18|H2QL18_PANTR         -----ENEISDASEVEKEI-NKAQNAIAGEGMGISHELITLEISSRDVPDLTLIDLPGIT 183
SP|A1E2I4|MX1_MACMU            -----EIEILDASEVEKEI-NKAQNTIAGEGMGISHELITLEISSRDVPDLTLIDLPGIT 183
SP|Q5R5G3|MX1_PONAB            -----EIEISDASEVEKEI-NKAQNTIAGEGMGISHELITLEISSRDVPDLTLIDLPGIT 183
TR|A0A6P8QPW8|A0A6P8QPW8_GEOSA -----KRTLRSSSEVEYEI-IKAQNAIAGEEGRICENVISLEIISPDVPDLTLIDLPGIV 147
TR|U3EGN4|U3EGN4_CALJA         -----EIEISDASEVEKEV-NKAQNTIAGEGMGISHELITLEISSRDVPDLTLIDLPGIT 183
SP|Q9N0Y3|MX1_CANLF            -----EMEISDPSEVEVEI-NKAQDAIAGEGQGISHELISLEVSSPHVPDLTLIDLPGIT 179
TR|M3W8X0|M3W8X0_FELCA         -----ETEISDPSEVEEAI-NTAQNAIAGEGLGISHELINLEVSSPHVPDLTLIDLPGIT 181
TR|A0A667GNL8|A0A667GNL8_LYNCA -----ETEISDPSEVEEAI-NTAQNAIAGEGLGISHELINLEISSPHVPDLTLIDLPGIT 181
TR|A0A6P6GZN4|A0A6P6GZN4_PUMCO -----ETEISDPSEVEEAI-NTAQNAIAGEGLGISHELINLEVSSPHVPDLTLIDLPGIT 181
TR|A0A6J1XXY6|A0A6J1XXY6_ACIJB -----ETEISDPSEVEEAI-NTAQNAIAGEGLGISHELINLEVSSPHVPDLTLIDLPGIT 181
TR|A0A6P4TS13|A0A6P4TS13_PANPR -----ETEISDPSEVEEAI-NTAQNAIAGEGLGISHELINLEVSSPHVPDLTLIDLPGIT 219
TR|A0A3Q7SCG7|A0A3Q7SCG7_VULVU -----EMEISDPSEVEVEI-NKAQDAIAGEGQGISHELISLEVSSPHVPDLTLIDLPGIT 217
TR|I3MJX6|I3MJX6_ICTTR         -----ELELSEASQVEEEI-NKAQNLIAGTGLGISDELISLEVSSPSVPDLTLIDLPGIT 184
TR|G9KBY2|G9KBY2_MUSPF         -----EKEISDPSEVEAEI-NKAQNAIAGEGQGISHELINLEVSSSHVPDLTLIDLPGIT 178
TR|A0A2Y9KR88|A0A2Y9KR88_ENHLU -----EKEISDPSEVEAEI-NKAQNAIAGEGQGISHELINLEVSSSHVPDLTLIDLPGIT 178
SP|P09922|MX1B_MOUSE           -----EVELSDPSEVEEAI-NKGQNFIAGVGLGISDKLISLDVSSPNVPDLTLIDLPGIT 149
SP|P18588|MX1_RAT              -----EIEISHPSLVEREI-NKAQNLIAGEGLKISSDLISLEVSSPHVPDLTLIDLPGIT 174
TR|A0A6P6D2J7|A0A6P6D2J7_PTEVA -----EIEMSDASEVEEEI-RKAQDVIAGVGMDISHELINLEISSPHVPDLTLIDLPGIT 188
TR|A0A6J0DKA3|A0A6J0DKA3_PERMB -----------------------QNFIAGNGLMISSKLISLEVSSPNVPDLTLIDLPGIT 173
TR|G5C5C9|G5C5C9_HETGA         -----QEDIADTSKVEGAV-IKAQNFITGTRLGIKDELISLEISSPSVPDLTLIDLPGIT 183
TR|A0A3Q0CWV5|A0A3Q0CWV5_MESAU -----EVEISDPTQVEGEI-NKAQNFIAGKGLGISDELISLDVSSPSVPDLTLIDLPGIT 168
TR|A0A6I9K5X3|A0A6I9K5X3_CHRAS -----EIDIANPLEVEKEV-NKAQNAIAGDGMGISRELINLEIISPNVPDLTLIDLPGIT 184
SP|P79135|MX1_BOVIN            -----EIEIPDASQVEKEI-SEAQIAIAGEGTGISHELISLEVSSPHVPDLTLIDLPGIT 172
TR|A0A6P3H2K4|A0A6P3H2K4_BISBI -----EIEIPDASQVEKEI-SEAQIAIAGEGTGISHELISLEVSSPHVPDLTLIDLPGIT 178
TR|L8IPR7|L8IPR7_9CETA         -----EIEIPDASQVEKEI-SEAQIAIAGEGTGISHELISLEVSSPHVPDLTLIDLPGIT 175
SP|P27594|MX1_PIG              -----EIELSDASQVEKEV-SAAQIAIAGEGVGISHELISLEVSSPHVPDLTLIDLPGIT 184
TR|A0A1U7U9K3|A0A1U7U9K3_CARSF -----ETEISDALEVEKEI-TKAQNVIAGEGMGISHELISLEVSSPHVPDLTLIDLPGIT 148
SP|P33237|MX1_SHEEP            -----EIEISDASQVEKEI-SEAQIAIAGEGMGISHELISLEVSSPHVPDLTLIDLPGIT 178
TR|A0A7D4WPX4|A0A7D4WPX4_CAPHI -----EIEISDASQVEKEI-SEAQIAIAGEGMGISHELISLEVSSPHVPDLTLIDLPGIT 178
SP|Q28379|MX1_HORSE            -----EVEISNALDVEEQV-RKAQNVLAGEGVGISQELVTLEVSSPHVPDLTLIDLPGIT 183
TR|A0A6I9I6Y4|A0A6I9I6Y4_VICPA -----EAEISDPLQVEKEV-NQAQIAIAGEGVGISHELISLEVSSPNVPDLTLIDLPGIT 183
TR|A0A6P5BYP3|A0A6P5BYP3_BOSIN -----EIEIPDASQVEKEI-SEAQIAIAGEGTGISHELISLEVSSPHVPDLTLIDLPGIT 175
TR|S9XPZ9|S9XPZ9_CAMFR         -----EAEISDPLQVEKEV-NQAQIAIAGEGVGISHELITLEVSSPNVPDLTLIDLPGIT 183
TR|A0A384DMX3|A0A384DMX3_URSMA -----EMEISDPSEVEVEI-NKAQDAIAGEGQGISHELITLEVSSPHVPDLTLIDLPGIT 181
TR|A0A3Q7W4X3|A0A3Q7W4X3_URSAR -----EMEISDPSEVEVEI-NKAQDAIAGEGQGISHELITLEVSSPHVPDLTLIDLPGIT 181
TR|A0A7E6D288|A0A7E6D288_9CHIR -----EINLSSPSEVEKEI-SKSQNLMAGEGVGISHELISLEVSSPHVPDLTLIDLPGIT 183
TR|S7NSW7|S7NSW7_MYOBR         -----EIDLSAASEVEQEI-RKAQNVIAGEGVGISQELINLEVSSPHVPDLTLIDLPGIT 203
TR|A0A1N7THY9|A0A1N7THY9_PIPPI -----EIDLTAASEVEQEI-RKAQNVIAGEGVGISQELINLEVSSPHVPDLTLIDLPGIT 179
TR|A0A1N7THZ0|A0A1N7THZ0_ROUAE -----EIEISDPLAVEKEI-SKAQDVIAGVGVDISHELINLEVSSPHAPDLTLIDLPGIT 181
TR|A0A1N7THY7|A0A1N7THY7_HYPMN -----EVEIPDPSAVEREI-RKAQDAMAGVGVGISHELITLEVSSPHAPDLTLIDLPGIT 181
TR|A0A1N7THZ1|A0A1N7THZ1_STULI -----EINLSDPSEVEKEI-NKSQNVIAGEGVGISHELISLEVRSPHVPDLTLIDLPGIT 181
TR|B2X026|B2X026_CHICK         -----EIQLQNASEVKKAI-RKAQDIVAGTNGSITGELISLEIWSPDVPDLTLIDLPGIA 227
TR|A0A172PZ65|A0A172PZ65_ANSCY -----SIELQHASEVESAI-SDAQDVVAGTKRNISGELISLEIYSPDVPDLTLIDLPGIA 226
TR|A0A0Q3PUG8|A0A0Q3PUG8_AMAAE -----NIELHNASEVDNAI-R--------------------------------------- 166
SP|Q4ADG6|MX1_PHOVI            -----EMEISDPSEVEVEI-SKAQNVIAGEGQGISHELISLEVSSPHVPDLTLIDLPGIT 181
SP|Q4ADG7|MX1_OTABY            -----EMEISDPSEVEVEI-NKAQNAIAGEGQGISHELISLEVSSPHVPDLTLIDLPGIT 181
```

```
TR|A0A2Y9G772|A0A2Y9G772_NEOSC -----EMEISDPSEVEVEI-SKAQNVIAGEGQGISHELISLEVSSPHVPDLTLIDLPGIT 181
TR|A0A6J2C7I6|A0A6J2C7I6_ZALCA -----EMEISDPSEVEVEI-NKAQNAIAGEGQGISHELISLEVSSPHVPDLTLIDLPGIT 181
TR|A0A2U3WDI8|A0A2U3WDI8_ODORO -----EMEISDPSEVEVEI-SKAQNAIAGEGQGISHELISLEVSSPHVPDLTLIDLPGIT 181
TR|A0A1U7SNC8|A0A1U7SNC8_ALLSI -----EEKLENPMQVEKAI-RKAQNIIAGEGVGISQELISLEISSPDVPDLTLIDLPGIV 254
TR|A0A7M4FFF4|A0A7M4FFF4_CROPO -----EEKLENPMQVEKAI-RKAQNTIAGEGVGISQELISLEISSPTVPDLTLIDLPGIV 162
TR|A0A340WTU5|A0A340WTU5_LIPVE -----ETEISDASQVEKESVKVTQVAIAGGGTGISHELISLEVTSPHVPDLTLIDLPGIT 184
TR|A0A6J3RAX5|A0A6J3RAX5_TURTR -----ETEISDASQVEKEI-SEAQVAIAGEGTGISHELISLEVTSPHVPDLTLIDLPGIT 183
TR|A0A341CI24|A0A341CI24_NEOAA -----ETEISNASQVEKEI-SEAQVAIAGKGMGISHELINLEVTSPHVPDLTLIGLPGIT 260
TR|A0A2Y9LTL6|A0A2Y9LTL6_DELLE -----ETEISNASQVEKEI-SDAQVAIAGEGMGISHELINLEVTSPHVPDLTLIDLPGIT 183
TR|A0A383ZCI9|A0A383ZCI9_BALAS -----ETEISDASQVEKEI-SEAQVAIAGEGTGISHELISLEVSSPHVPDLTLIDLPGIT 182
TR|V9KIB1|V9KIB1_CALMI         -----RQELTDPSEVEQEI-RIAQDIIAGKGVGISHELISLQIEASNVPDLTLIDLPGIA 147
TR|A0A674K7E1|A0A674K7E1_TERCA -----KQELNGPSEVEKEI-RKAQDAMAGEGVGISQELISLEISSPNVPDLTLIDLPGIA 189
TR|A0A4D9DW37|A0A4D9DW37_9SAUR -----DEELHHPLVVDKEI-RKAQNAIAGEGVGISQELISLEIRSPNVPDLTLIDLPGIA 338
TR|H2DQ54|H2DQ54_XENLA         -----ELTIASAAEVENQV-KQAQNLMAGNGKGISDELISLEVISPDVPDLTLIDLPGIT 147
TR|A0A0F6MUS9|A0A0F6MUS9_ANGJA ---DREVEIYDPDDVERLI-REAQDEMAGVGVGISDDLISLEISSPGVPDLTLIDLPGIA 162
TR|Q98990|Q98990_SALSA         ---DHEEEIEDPSDVEKKI-REAQDEMAGVGVGISDDLISLEIGSPDVPDLTLIDLPGIA 147
TR|A0A674EGU5|A0A674EGU5_SALTR ---DHEEEIEDPSDVEKKI-REAQDDMAGVGVGISDDLISLEIGSPDVPDLTLIDLPGIA 147
SP|Q91192|MX1_ONCMY            ---DHEEEIEDPSDVEKKI-REAQDEMAGVGVGISDDLISLEIGSPDVPDLTLIDLPGIA 147
TR|A0A286K200|A0A286K200_MYLPI -----EEDIDDPAEVEKKI-REAQDEMAGAGVGISDELISLQITSANVPDLTLIDLPGIA 150
TR|A0A498MHV8|A0A498MHV8_LABRO ----EAEDIHDPAEVEKKI-REAQDEMAGVGVGISDELISLQITSANVPDLTLIDLPGIA 151
SP|Q7T2P0|MX1_ICTPU            HDEDYEEEIQNPADVEKKI-REAQDHMAGVGVGISDELISLEVTSADVPDLTLIDLPGIA 152
TR|A0A556TK66|A0A556TK66_BAGYA -DQDYEEEIEDPAEVESLI-REAQDQMAGIGVGISDDLISLEVTSSDVPDLTLIDLPGIT 151
TR|A0A3G6IID3|A0A3G6IID3_9TELE -----EEDIDDPAEVEKKI-REAQDEMAGAGVGISDELISLQITSANVPDLTLIDLPGIA 150
SP|Q8JH68|MXA_DANRE            -----EEDFDDPAEVEKKI-RQAQDEMAGAGVGISEELISLQITSADVPDLTLIDLPGIA 150
TR|A0A672MEY2|A0A672MEY2_SINGR -----RETFNDPLKVDTLV-REAQNKLAGKAVGICDELITLEISSPDVCDLTLIDLPGIT 149
TR|Q7T2M3|Q7T2M3_CARAU         -----EEDIDDPAEVEKKI-REAQDEMAGAGVGISDELISLQITSANVPDLTLIDLPGIA 150
TR|A0A673LWX8|A0A673LWX8_9TELE -----EKDIHDPAEVDKMI-CEAQDEIAGKGVGISDELISLQITSANVPDLTLIDLSGIA 150
TR|A0A8B9KF29|A0A8B9KF29_ASTMX ---GYEEELDDPAEVESKI-REAQDKIAGVGVGISDDLISLEITSSNVPDLTLIDLPGIA 147


SP|P20591|MX1_HUMAN            RVAVGNQPADIGYKIKTLIKKYIQRQETISLVVVPSNVDIATTEALSMAQEVDPEGDR-- 241
TR|K4JBQ0|K4JBQ0_9PRIM         RVAVGNQPADIGYKIKTLIKKYIQRQETISLVVVPSNVDIATTEALSMAQEVDPEGDR-- 241
TR|H2QL18|H2QL18_PANTR         RVAVGNQPADIGYKIKTLIKKYIQRQETISLVVVPSNVDIATTEALSMAQEVDPEGDR-- 241
SP|A1E2I4|MX1_MACMU            RVAVGNQPPDIGYKIKTLIRKYIQRQETINLVVVPSNVDIATTEALSMAQEVDPEGDR-- 241
SP|Q5R5G3|MX1_PONAB            RVAVGNQPADIGYKIKTLIKKYIQRQETISLVVVPSNVDIATTEALSMAQEVDPEGDR-- 241
TR|A0A6P8QPW8|A0A6P8QPW8_GEOSA RVAVGNQPENIGTQIKKLIQNYIEKQETIILVVAPCNVDIATIEALAIAHEVDPNGKR-- 205
TR|U3EGN4|U3EGN4_CALJA         RVAVGNQPADIGRKIKALIRKYIQRQETISLVVVPSNVDIATTEALSMAQEVDPEGDR-- 241
SP|Q9N0Y3|MX1_CANLF            RVAVGNQPADIGRQTKQLIRKYILKQETINLVVVPCNVDIATTEALSMAQEVDPDGDR-- 237
TR|M3W8X0|M3W8X0_FELCA         RVAVGNQPADIGRQTKQLIRKYIVKQETINLVVVPCNVDIATTEALSMAQEVDPDGDR-- 239
TR|A0A667GNL8|A0A667GNL8_LYNCA RVAVGNQPADIGRQTKQLIRKYIVKQETINLVVVPCNVDIATTEALSMAQEVDPDGDR-- 239
TR|A0A6P6GZN4|A0A6P6GZN4_PUMCO RVAVGNQPADIGRQTKQLIRKYIVKQETINLVVVPCNVDIATTEALSMAQEVDPDGDR-- 239
TR|A0A6J1XXY6|A0A6J1XXY6_ACIJB RVAVGNQPADIGRQTKQLIRKYIVKQETINLVVVPCNVDIATTEALSMAQEVDPDGDR-- 239
TR|A0A6P4TS13|A0A6P4TS13_PANPR RVAVGNQPADIGRQTKQLIRKYIVKQETINLVVVPCNVDIATTEALSMAQEVDPDGDR-- 277
TR|A0A3Q7SCG7|A0A3Q7SCG7_VULVU RVAVGNQPADIGRQTKQLIRKYILKQETINLVVVPCNVDIATTEALSMAQEVDPDGDR-- 275
TR|I3MJX6|I3MJX6_ICTTR         RVAVGNQPTDIGRQIKRLIRKYIQKQETINLVVVPSNVDIATTEALSMAQEVDPDGDR-- 242
TR|G9KBY2|G9KBY2_MUSPF         RVAVGNQPADIGRQTKQLIRKYILRQETINLVVVPCNVDIATTEALSMAQEVDPDGDR-- 236
TR|A0A2Y9KR88|A0A2Y9KR88_ENHLU RVAVGNQPADIGRQTKQLIRKYILRQETINLVVVPCNVDIATTEALSMAQEVDPDGDR-- 236
SP|P09922|MX1B_MOUSE           RVAVGNQPADIGRQIKRLIKTYIQKQETINLVVVPSNVDIATTEALSMAQEVDPEGDR-- 207
SP|P18588|MX1_RAT              RVAVGDQPADIEHKIKRLITEYIQKQETINLVVVPSNVDIATTEALKMAQEVDPQGDR-- 232
TR|A0A6P6D2J7|A0A6P6D2J7_PTEVA RVALGNQPADIGFQIKRLIKKYILKQQTINLVVVPSNVDIATTEALSMAQEVDPDGDR-- 246
TR|A0A6J0DKA3|A0A6J0DKA3_PERMB RVAVGNQPADIGC------------QETINLVVVPSNVDIATTEALSMAQEVDPEGDR-- 219
TR|G5C5C9|G5C5C9_HETGA         RVAVGNQPEDIGCQIKRLIMKYIEKAETINLVVVPSNVDIATTEALSMAKQVDPEGDR-- 241
TR|A0A3Q0CWV5|A0A3Q0CWV5_MESAU RVAVGNQPADIGHQIKRLIKKYIQRQETINLVVVPSNVDIATTEALSMAKEVDPEGDR-- 226
TR|A0A6I9K5X3|A0A6I9K5X3_CHRAS RVAVGDQPIDIGRQIKTLIKSYIQKQETINLVVVPCNVDIATTEALSMAQEVDPDGDR-- 242
SP|P79135|MX1_BOVIN            RVAVGNQPPDIEYQIKSLIRKYILRQETINLVVVPANVDIATTEALRMAQEVDPQGDR-- 230
TR|A0A6P3H2K4|A0A6P3H2K4_BISBI RVAVGNQPPDIEYQIKSLIRKYILRQETINLVVVPANVDIATTEALRMAQEVDPQGDR-- 236
TR|L8IPR7|L8IPR7_9CETA         RVAVGNQPPDIEYQIKSLIRKYILRQETINLVVVPANVDIATTEALHMAQEVDPQGDR-- 233
SP|P27594|MX1_PIG              RVAVGNQPYDIEYQIKSLIKKYICKQETINLVVVPCNVDIATTEALRMAQEVDPEGDR-- 242
TR|A0A1U7U9K3|A0A1U7U9K3_CARSF RVAVGNQPADIGRQIKALIRKYIYRQQTINLVVVPCNVDIATTEALSMAQEVDPEGDR-- 206
SP|P33237|MX1_SHEEP            RVAVGNQPHDIEYQIKSLIRKYILRQETINLVVVPANVDIATTEALRMAQDVDPQGDR-- 236
TR|A0A7D4WPX4|A0A7D4WPX4_CAPHI RVAVGNQPPDIEYQIKSLIRKYILRQETINLVVVPANVDIATTEALRMAQDVDPQGDR-- 236
SP|Q28379|MX1_HORSE            RVAVGNQPADIGRQIKTLIRKYIQRQETINLVVVPSNVDIATTEALSMAQEVDPEGDR-- 241
TR|A0A6I9I6Y4|A0A6I9I6Y4_VICPA RVAVGNQPHDIEYQIKSLIRKYIQRQETINLVVVPCNVDIATTEALSMAQEVDPDGDR-- 241
TR|A0A6P5BYP3|A0A6P5BYP3_BOSIN RVAVGNQPPDIEYQIKSLIRKYILRQETINLVVVPANVDIATTEALRMAQEVDPQGDR-- 233
TR|S9XPZ9|S9XPZ9_CAMFR         RVAVGNQPHDIEYQIKSLIRKYIQRQETINLVVVPCNVDIATTEALRMAQEVDPDGDR-- 241
TR|A0A384DMX3|A0A384DMX3_URSMA RVAVGNQPADIGRQSKQLIRKYIFKQETINLVVVPCNVDIATTEALSMAQEVDPDGDR-- 239
TR|A0A3Q7W4X3|A0A3Q7W4X3_URSAR RVAVGNQPADIGRQSKQLIRKYIFKQETINLVVVPCNVDIATTEALSMAQEVDPDGDR-- 239
TR|A0A7E6D288|A0A7E6D288_9CHIR RVPVGNQPPDIEYQIKTLIRKYIQRQETINLVVVPCNVDIATTEALRMAQEVDPEGDR-- 241
TR|S7NSW7|S7NSW7_MYOBR         RVAVGNQPADIGRQITALIKKYILRQQTIMLVVVPSNVDIATTEALSMAHEVDPDGDR-- 261
TR|A0A1N7THY9|A0A1N7THY9_PIPPI RVAVGNQPADIGRQITTLIKKYILRQQTIMLVVVPSNVDIATTEALSMAQEVDPEGDR-- 237
TR|A0A1N7THZ0|A0A1N7THZ0_ROUAE RVAVGNQPADIGRQIKALIRKYIRKQETINLVVVPSNVDIATTEALSMAQEVDPDGDR-- 239
TR|A0A1N7THY7|A0A1N7THY7_HYPMN RVAVGNQPADIGWQIKALIRKYIQKQETINLVVVPSNVDIATTEALSMAQEVDPDGDR-- 239
TR|A0A1N7THZ1|A0A1N7THZ1_STULI RVAMGNQPADIGHQIKRLIRKYIQRQETINLVVVPCNVDIVTTEALSMAQEVDPEGDR-- 239
TR|B2X026|B2X026_CHICK         REAVGNQPQDNGQQIKTLLKKYIGCKETIIVVVVPCNVDIATTEALKMAQEVDPTGER-- 285
TR|A0A172PZ65|A0A172PZ65_ANSCY RVAVRNQPQDIGEQIKMLIKKIISSKETINLVVVPCNVDIATTEALKMAQEVDPKGER-- 284
TR|A0A0Q3PUG8|A0A0Q3PUG8_AMAAE ------------------------------------------------------------
SP|Q4ADG6|MX1_PHOVI            RVAVGNQPADIGRQTKQLIRKYILKQETINLVVVPCNVDIATTEALSMAQEVDPSGDR-- 239
SP|Q4ADG7|MX1_OTABY            RVAVGNQPADIGHQTKKLIKKYILKQETINLVVVPCNVDIATTEALSMAQEVDPDGDR-- 239
TR|A0A2Y9G772|A0A2Y9G772_NEOSC RVAVGNQPADIGRQTKQLIKKYILKQETINLVVVPCNVDIATTEALSMAQEVDPDGDR-- 239
TR|A0A6J2C7I6|A0A6J2C7I6_ZALCA RVAVGNQPADIGHQTKKLIKKYILKQETINLVVVPCNVDIATTEALSMAEEVDPDGDR-- 239
TR|A0A2U3WDI8|A0A2U3WDI8_ODORO RVAVGNQPADIGCQTKKLIKKYILKQETINLVVVPCNVDIATTEALSMAQEVDPDGDR-- 239
TR|A0A1U7SNC8|A0A1U7SNC8_ALLSI RVAVGNQPTDIGEQIKNLIKKFIVKRETINLVVISSNVDLATTEALKMAQEVDPEGER-- 312
TR|A0A7M4FFF4|A0A7M4FFF4_CROPO RVAVGNQPTDIGEQIKTLIKTHIVKQETINLVVISSNVDLATTEALKMAQEVDPKGER-- 220
TR|A0A340WTU5|A0A340WTU5_LIPVE RIAVGNQPADIEYQIKSLIRKYILKQETINLVVVPANVDIATMEALCMAQEVDLQGLDHS 244
TR|A0A6J3RAX5|A0A6J3RAX5_TURTR RIAVGNQPANIEYQIKSLIRNYILKQETINLVVVPANVDIATVEALRMAQEVEXSKD--- 240
TR|A0A341CI24|A0A341CI24_NEOAA RIAVGNQPANTEYQIKSLIRKYILKQETINLVVVPANVDITTMEALCMAQEVDLQGQDHS 320
TR|A0A2Y9LTL6|A0A2Y9LTL6_DELLE RIAVGNQPANTEYQIKSLIRKYILKQETINLVVVPANVDITTMEALCMAQEVDLQGQDHS 243
TR|A0A383ZCI9|A0A383ZCI9_BALAS RIAVGNQPADIEYQIKSLIRKYILKQETINLVVVPANVDIATTEALRMAQEVDPQGDR-- 240
```

```
TR|V9KIB1|V9KIB1_CALMI         RVAVGKQPLNIGDQIKQLIKSFIEKQETINLVVVPCNVDIATTEALKMAQEVDPSGER-- 205
TR|A0A674K7E1|A0A674K7E1_TERCA RVAVGNQPKDIGNQIINLIKKYICKQQTINLVVVPSNVDIATTEALKMAKEVDPKGER-- 247
TR|A0A4D9DW37|A0A4D9DW37_9SAUR RVAVGNQPQDIGEQIKKLIKKFIAKQETINLVVVPSNVDIATTEALKMAQEVDPDGER-- 396
TR|H2DQ54|H2DQ54_XENLA         RVALPDQPKDIGQQIKKMIRKYIQRQETINLVVVPSNVDIATTEALEMAREVDPNGER-- 205
TR|A0A0F6MUS9|A0A0F6MUS9_ANGJA RVAVKGQPENIGDQIKRLITKFVRKQETINLVVVPCNVDIATTEALKMAQEEDPEGER-- 220
TR|Q98990|Q98990_SALSA         RVAVKGQPENIGEQIKRLIRKFITKQETINLVVVPCNVDIATTEALKMAQEVDPEGER-- 205
TR|A0A674EGU5|A0A674EGU5_SALTR RVAVKGQPENIGEQIKRLIRKFIMKQETINLVVVPCNVDIATTEALKMAQEVDPEGER-- 205
SP|Q91192|MX1_ONCMY           RVAVKGQPENIGEQIKRLIRKFIMKQETISLVVVPCNVDIATTEALKMAQEVDPEGER-- 205
TR|A0A286K200|A0A286K200_MYLPI RVAVKGQPENIGDQIKRLIRKFVTKQETINLVVVPCNVDIATTEALQMAQEEDPEGER-- 208
TR|A0A498MHV8|A0A498MHV8_LABRO RVAVKGQPENIGDQIKRLIRKFVTRQETINLVVVPCNVDIATTEALKMAQEEDPEGER-- 209
SP|Q7T2P0|MX1_ICTPU           RVAVKGQPENIGEQIKRLIKKFITKQETINLVVVPSNVDIATTEALKMAQEVDPNGER-- 210
TR|A0A556TK66|A0A556TK66_BAGYA RVAVKGQPENIGEQINRLIRKFITKQETINLVVVPSNVDIATTEALKMAQEVDPDGER-- 209
TR|A0A3G6IID3|A0A3G6IID3_9TELE RVAVKGQPENIGEQIKRLIRKFVTKQETINLVVVPCNVDIATTEALQMAQEEDPEGER-- 208
SP|Q8JH68|MXA_DANRE           RVAVKGQPENIGDQIKRLIRKFITKQETINLVVVPCNVDIATTEALQMAQAEDPDGER-- 208
TR|A0A672MEY2|A0A672MEY2_SINGR RVPVHDQPEDIGDQIKTLILKYIAKSETIILVVVPCNVDIATTEALRMAQKVDPEGLR-- 207
TR|Q7T2M3|Q7T2M3_CARAU         RVAVKGQPENIGDQIKRLIRKFVTKQETINLVVVPCNVDIATTEALQMAQEEDPEGER-- 208
TR|A0A673LWX8|A0A673LWX8_9TELE RVAVKGQPENIGDQIKRLIRMFVSKQETINLVVVPCNVDIATTEALHMAQKDDPEGER-- 208
TR|A0A8B9KF29|A0A8B9KF29_ASTMX RVAVKGQPEDIGDQIKRLIRKFITKQETINLVVVPCNVDIATTEALKMAQVEDPNGER-- 205


SP|P20591|MX1_HUMAN           --TIGILTKPDLVDKGTEDKVVDVVRNLVFHLKKGYMIVKCRGQQEIQDQLSLSEALQRE 299
TR|K4JBQ0|K4JBQ0_9PRIM         --TIGILTKPDLVDRGTEDKVVDVVRNLVFHLKKGYMIVKCRGQQEIQDQLSLSEALQRE 299
TR|H2QL18|H2QL18_PANTR         --TIGILTKPDLVDKGTEDKVVDVVRNLVFHLKKGYMIVKCRGQQEIQDQLSLSEALQRE 299
SP|A1E2I4|MX1_MACMU           --TIGILTKPDLVDKGTEDKVVDVVRNLVFHLKKGYMIVKCRGQQEIQDQLSLSEALQRE 299
SP|Q5R5G3|MX1_PONAB           --TIGILTKPDLVDKGTEDKVVDVVRNLVFHLKKGYMIVKCRGQQEIQDQLSLSEALQRE 299
TR|A0A6P8QPW8|A0A6P8QPW8_GEOSA --TLGILTKPDLVDGGSENLILETMRNSKISLNKGYMIVKCRGQQAIHDNLSLKDAVQEE 263
TR|U3EGN4|U3EGN4_CALJA         --TIGILTKPDLVDKGTEDKVVDVVRNLVFHLKKGYMIVKCRGQQEIQDQLSLSEALERE 299
SP|Q9N0Y3|MX1_CANLF           --TIGILTKPDLVDRGTEGKVVDVAQNLVCHLKKGYMIVKCRGQQDIQDQVSLAEALQKE 295
TR|M3W8X0|M3W8X0_FELCA         --TIGILTKPDLVDRGTEDKVVDVAQNLVCHLKKGYMVVKCRGQQDIQDQLSLAEALEKE 297
TR|A0A667GNL8|A0A667GNL8_LYNCA --TIGILTKPDLVDRGTEDKVVDVAQNLVCHLKKGYMVVKCRGQQDIQDQLSLAEALEKE 297
TR|A0A6P6GZN4|A0A6P6GZN4_PUMCO --TIGILTKPDLVDRGTEDKVVDVAQNLVCHLKKGYMVVKCRGQQDIQDQLSLAEALEKE 297
TR|A0A6J1XXY6|A0A6J1XXY6_ACIJB --TIGILTKPDLVDRGTEDKVVDVAQNLVCHLKKGYMVVKCRGQQDIQDQLSLAEALEKE 297
TR|A0A6P4TS13|A0A6P4TS13_PANPR --TIGILTKPDLVDRGTEDKVVDVAQNLVCHLKKGYMVVKCRGQQDIQDQLSLAEALEKE 335
TR|A0A3Q7SCG7|A0A3Q7SCG7_VULVU --TIGILTKPDLVDRGTEGKVVDVAQNLVCHLKKGYMIVKCRGQQDIQDQVSLAEALQKE 333
TR|I3MJX6|I3MJX6_ICTTR         --TIGILTKPDLVDKGTEDTVVDVVRNLVCHLKKGYMIVKCRGQQDIQSRLSLAEALQKE 300
TR|G9KBY2|G9KBY2_MUSPF         --TIGILTKPDLVDRGTESKVVDVAQNLVCHLKKGYMIVKCRGQQDIQDQVTLAEALQKE 294
TR|A0A2Y9KR88|A0A2Y9KR88_ENHLU --TIGILTKPDLVDRGTEGKVVDVAQNLVCHLKKGYMIVKCRGQQDIQDQVTLAEALQKE 294
SP|P09922|MX1B_MOUSE          --TIGVLTKPDLVDRGAEGKVLDVMRNLVYPLKKGYMIVKCRGQQDIQEQLSLTEAFQKE 265
SP|P18588|MX1_RAT             --TIGILTKPDLVDRGTEDKVVDVVRNLVCHLKKGYMIVKCRGQQDIQEQLSLAEALQKE 290
TR|A0A6P6D2J7|A0A6P6D2J7_PTEVA --TIGILTKPDLVDKGTENKVVDVVRNLVCHLKKGYMIVKCRGQQDIQDQLSLAEALQKE 304
TR|A0A6J0DKA3|A0A6J0DKA3_PERMB --TIGRRVGG-------NSDVGLLVRTQA-LDPDGPILVHTQGPTEEG-----------R 258
TR|G5C5C9|G5C5C9_HETGA         --TIGILTKPDLVDKGTEDKVVDVVHNLLCPLKKGYMVVRCRSQQDIQARLSPTEALQRE 299
TR|A0A3Q0CWV5|A0A3Q0CWV5_MESAU --TIGILTKPDLVDRGTEDKVVDVVRNLVYHLKKGYMVVKCRGQQDVQKQLNLAEALKME 284
TR|A0A6I9K5X3|A0A6I9K5X3_CHRAS --TIGILTKPDLVDRGTEGKVVDVVRNLVYHLKKGYMIVKCRGQQDIQDKQSLAEALQRE 300
SP|P79135|MX1_BOVIN           --TIGILTKPDLVDKGTEDKVVDVVRNLVFHLKKGYMIVKCRGQQDIKHRMSLDKALQRE 288
TR|A0A6P3H2K4|A0A6P3H2K4_BISBI --TIGILTKPDLVDKGTEDKVVDVVRNLVFHLKKGYMIVKCRGQQDIKHQMSLDKALQRE 294
TR|L8IPR7|L8IPR7_9CETA         --TIGILTKPDLVDKGTEDKVVDVVRNLVFHLKKGYMIVKCRGQQDIKHRMSLDKALQRE 291
SP|P27594|MX1_PIG             --TIGILTKPDLVDKGTEDKIVDVARNLVFHLKKGYMIVKCRGQQDIQEQLSLAKALQKE 300
TR|A0A1U7U9K3|A0A1U7U9K3_CARSF --TIGILTKPDLVDKGTEDQIVDVARNLVCHLKKGYMIVKCRGQQEIQDQLSLSEALQRE 264
SP|P33237|MX1_SHEEP           --TIGILTKPDLVDKGTEDKVVDVVRNLVFHLKKGYMIVKCRGQQEIQHRLSLDKALQRE 294
TR|A0A7D4WPX4|A0A7D4WPX4_CAPHI --TIGILTKPDLVDKGTEDKVVDVVRNLVFHLKKGYMIVKCRGQQEIQHRLSLDKALQRE 294
SP|Q28379|MX1_HORSE           --TIGILTKPDLVDKGTEEQVVDVVRNLICHLKKGYMIVKCRGQQDIQDRLSLAEALQRE 299
TR|A0A6I9I6Y4|A0A6I9I6Y4_VICPA --TIGILTKPDLVDKGTEDKVVDVVRNLVFHLKKGYMVVKCRGQQDIQARLSLAKALQRE 299
TR|A0A6P5BYP3|A0A6P5BYP3_BOSIN --TIGILTKPDLVDKGTEDKVVDVVRNLVFHLKKGYMIVKCRGQQDIKHRMSLDKALQRE 291
TR|S9XPZ9|S9XPZ9_CAMFR         --TIGILTKPDLVDKGTEDKVVDVVRNLIFHLKKGYMIVKCRGQQDIQARLSLAKALQRE 299
TR|A0A384DMX3|A0A384DMX3_URSMA --TIGILTKPDLVDRGTESKVVDVAQNLVCHLKKGYMIVKCRGQQDIQNQVTLAEALQKE 297
TR|A0A3Q7W4X3|A0A3Q7W4X3_URSAR --TIGILTKPDLVDRGTESKVVDVAQNLVCHLKKGYMIVKCRGQQDIQDQVTLAEALQKE 297
TR|A0A7E6D288|A0A7E6D288_9CHIR --TIGILTKPDLVDRGTEDKVVDVVRNLVFHLKKGYMIVKCRGQQDIRGKMSLASALCKE 299
TR|S7NSW7|S7NSW7_MYOBR         --TIGILTKPDLVDRGTEDKVVDVVRNLVYHLKKGYMIVKCRGQQDIQYQMSLSKALQKE 319
TR|A0A1N7THY9|A0A1N7THY9_PIPPI --TIGILTKPDLVDRGTEDRIVDVVRNLVYHLKKGYMIVKCRGQQDIQYQISLAKALQRE 295
TR|A0A1N7THZ0|A0A1N7THZ0_ROUAE --TIGILTKPDLVDKGTEHRVVDVVRNLVCQLKKGYMIVKCRGQQDIQDQLSLAEALRKE 297
TR|A0A1N7THY7|A0A1N7THY7_HYPMN --TIGILTKPDLVDKGTEHRVVDVVRNLVCPLKKGYMIVKCRGQQDVQGQLSLAEALRKE 297
TR|A0A1N7THZ1|A0A1N7THZ1_STULI --TIGILTKPDLVDKGAEDKIVDVARNLVCHLKKGYMIVKCRGQQDIQDRMSLAQALCNE 297
TR|B2X026|B2X026_CHICK         --TLGVLTKPDLVNEGTEETVLKIIQNEVIPLRKGYMIVKCYGQMDFCNELSFTSAIQQE 343
TR|A0A172PZ65|A0A172PZ65_ANSCY --TLGILTKPDLVDIGTEESIISIVQNEVIPLRKGYMIVKCRGQRDIHNKLTLASAIQQE 342
TR|A0A0Q3PUG8|A0A0Q3PUG8_AMAAE ---QGILTKPDLVDRGTEESVIKILRNKVIPLKKGYMIVKCRGQEDIHNKLSLAAAIQQE 223
SP|Q4ADG6|MX1_PHOVI           --TIGILTKPDLVDRGTESKVVDVAQNLVCHLKKGYMIVKCRGQQDIQDQVTLTEALQKE 297
SP|Q4ADG7|MX1_OTABY           --TIGILTKPDLVDRGTESKVVDVAQNLVCHLKKGYMIVKCRGQQDIQDQVTLTEALQKE 297
TR|A0A2Y9G772|A0A2Y9G772_NEOSC --TIGILTKPDLVDRGTESKVVDVAQNLVCHLKKGYMIVKCRGQQDIQNQVTLTEALQKE 297
TR|A0A6J2C7I6|A0A6J2C7I6_ZALCA --TIGILTKPDLVDRGTESKVVDVAQNLVCHLKKGYMIVKCRGQQDIQDQVTLTEALQKE 297
TR|A0A2U3WDI8|A0A2U3WDI8_ODORO --TIGILTKPDLVDRGTESKVVDVAQNLVCHLKKGYMIVKCRGQQDIQDQVTLTEALQKE 297
TR|A0A1U7SNC8|A0A1U7SNC8_ALLSI --TLGILTKPDLLDKGTEHAVVDIVQNRVILLRKGYMMVKCRGQQDIQNNITLASAIQEE 370
TR|A0A7M4FFF4|A0A7M4FFF4_CROPO --TLGILTKPDLVDKGTEEAIVDIVRNLVIHLKKGYMVVKCRGQQDIQNNLSLDCAIQKE 278
TR|A0A340WTU5|A0A340WTU5_LIPVE LWVSGILTKPDLVDKGPXDKVVDMVRNLVFHLKKGYMVVKCRGQQDIQHQLSLTKALQEE 304
TR|A0A6J3RAX5|A0A6J3RAX5_TURTR -RTIGILTKPDLVDKSTEDKVVDMVRNLVFHLKKGYMIVKCRGQQDIQHQLSLTKALQKE 299
TR|A0A341CI24|A0A341CI24_NEOAA LWVAGILTKPDLVDKSTEDKVVDVVRNLVFHLKKGYMIIKCWGQQDIQHQLSLTKALQKE 380
TR|A0A2Y9LTL6|A0A2Y9LTL6_DELLE LWVAGILTKPDLVDKSTEDKVVDVVRNLVFHLKKGYMIIKCRGQQDIQHQLSLTKALQKE 303
TR|A0A383ZCI9|A0A383ZCI9_BALAS --TIGILTKPDLVDKGTEDKVVDVVRNLVFHLKKGYMIVKCRGQQDIQHQLSLTKALQKE 298
TR|V9KIB1|V9KIB1_CALMI         --TVGILTKPDLVDKGTEKTVVEIIQNQVVELKKGYIVVKCRGQQDINEKLTLRRALEKE 263
TR|A0A674K7E1|A0A674K7E1_TERCA --TLGILTKPDLVDRGTESNVVDIVRNLSIPLRKGYLIVKCRGQQDIHDNLTLASAIQKE 305
TR|A0A4D9DW37|A0A4D9DW37_9SAUR --TLELGK----------------------------TLLSCLA-------------NVS 412
TR|H2DQ54|H2DQ54_XENLA         --TLGILTKPDLVDRGAESDVVSVVRNLVYSLQKGYMIVKCRGQLEIQENLSLKEALDNE 263
TR|A0A0F6MUS9|A0A0F6MUS9_ANGJA --TLGILTKPDLVDKGTEETVVDIVHNEVIHLTKGYMIVKCRGQKEIKDKVSLSEAAYKE 278
TR|Q98990|Q98990_SALSA         --TLGILTKPDLVDKGTEETVVDIVHNEVIHLTKGYMIVKCRGQKEIMERVSLSEATERE 263
TR|A0A674EGU5|A0A674EGU5_SALTR --TLGILTKPDLVDKGTEETVVDIVHNEVIHLTKGYMIVKCRGQKEIMERVSLSEATERE 263
SP|Q91192|MX1_ONCMY           --TLGILTKPDLVDKGTEETVVDIVHNEVIHLTKGYMIVKCRGQKEIMERVSLTEATERE 263
TR|A0A286K200|A0A286K200_MYLPI --TLGILTKPDLVDKGTEGTVVDIVHNEVIHLTKGYMIVRCRGQKEIIDQVTLNEATATE 266
TR|A0A498MHV8|A0A498MHV8_LABRO --TLGILTKPDLVDKGTEETVVDIVHNEVIHLSKGYMIVRCRGQKEIIDQVTLNEATETE 267
```

```
SP|Q7T2P0|MX1_ICTPU            --TLGILTKPDLVDKGTEETVVSIIHNEIIYLTKGYMIVRCRGQKEIMDRVSLHEATEKE 268
TR|A0A556TK66|A0A556TK66_BAGYA --TLGILTKPDLVDKGTEDTVVSIIRNEIIYLKKGYMIVN-------------------- 247
TR|A0A3G6IID3|A0A3G6IID3_9TELE --TLGILTKPDLVDKGTEGTVVDIVHNEVIHLTKGYMIVRCRGQKEIIDQVTLNEATATE 266
SP|Q8JH68|MXA_DANRE            --TLGILTKPDLVDKGTEGTVVDIVHNEVIHLTKGYMIVRCRGQKEIMDQVTLNEATETE 266
TR|A0A672MEY2|A0A672MEY2_SINGR --TLAILTKPDLIDRGAEIDVLNIVQGKVIPLSKGYIIVRCRGQSDINDRIPFDKAMEAE 265
TR|Q7T2M3|Q7T2M3_CARAU         --TLGILTKPDLVDKGTEGTVVDIVHNEVIHLTKGYMIVRCRGQKEIIDHVTLNEATATE 266
TR|A0A673LWX8|A0A673LWX8_9TELE --TLGILTKPDLADKGTEETVVDIVHNEVIHLTKGYMIVRCRGQKEIIDQVTLHEATAAE 266
TR|A0A8B9KF29|A0A8B9KF29_ASTMX --TLGILTKPDLVDKGTEETVVEIVNNEIINLTKGYMIVRCRGQKEIIDRVSLTEATEKE 263
                                                                     ::

SP|P20591|MX1_HUMAN            KIFFENHPYFRDLLEE-----GKA----------------TVPCLAEKLTSE-------- 330
TR|K4JBQ0|K4JBQ0_9PRIM         KIFFQDHPYFRDLLEE-----GKA----------------TVPCLAEKLTSE-------- 330
TR|H2QL18|H2QL18_PANTR         KIFFQDHPYFRDLLEE-----GKA----------------TVPCLAEKLTSE-------- 330
SP|A1E2I4|MX1_MACMU            KIFFEDHPHFRDLLEE-----GKA----------------TIPCLAEKLTSE-------- 330
SP|Q5R5G3|MX1_PONAB            KIFFEDHPYFRDLLEE-----GKA----------------TVPCLAEKLTSE-------- 330
TR|A0A6P8QPW8|A0A6P8QPW8_GEOSA EEFFKKHKHFSPLLDD-----GKA----------------TIPILAEKLTTE-------- 294
TR|U3EGN4|U3EGN4_CALJA         KLFFEDHPYFRDLLEE-----GKA----------------TVPCLAEKLTSE-------- 330
SP|Q9N0Y3|MX1_CANLF            KDFFEDHPHFRVLLEE-----GRA----------------TVPNLAEKLTSE-------- 326
TR|M3W8X0|M3W8X0_FELCA         RVFFEDHPHFRVLLEE-----GRA----------------TVPCLADRLTTE-------- 328
TR|A0A667GNL8|A0A667GNL8_LYNCA RVFFEDHPHFRVLLEE-----GRA----------------TVPCLADRLTTE-------- 328
TR|A0A6P6GZN4|A0A6P6GZN4_PUMCO RVFFEDHPHFRVLLEE-----GRA----------------TVPCLADRLTTE-------- 328
TR|A0A6J1XXY6|A0A6J1XXY6_ACIJB RVFFEDHPHFRVLLEE-----GRA----------------TVPCLADRLTTE-------- 328
TR|A0A6P4TS13|A0A6P4TS13_PANPR RVFFEDHPHFRVLLEE-----GRA----------------TVPCLADRLTTE-------- 366
TR|A0A3Q7SCG7|A0A3Q7SCG7_VULVU KDFFEDHPHFRVLLEE-----GRA----------------TVPNLAEKLTSE-------- 364
TR|I3MJX6|I3MJX6_ICTTR         KAFFEDHPQFRALLDE-----GKA----------------TIPCLAERLTTE-------- 331
TR|G9KBY2|G9KBY2_MUSPF         RDFFEDHPHFRVLLEE-----GRA----------------TVPCLADKLTSE-------- 325
TR|A0A2Y9KR88|A0A2Y9KR88_ENHLU KDFFKDHPHFRVLLEE-----GRA----------------TVPCLADKLTSE-------- 325
SP|P09922|MX1B_MOUSE           QVFFKDHSYFSILLED-----GKA----------------TVPCLAERLTEE-------- 296
SP|P18588|MX1_RAT              QVFFKEHPQFRVLLED-----GKA----------------TVPCLAKRLTME-------- 321
TR|A0A6P6D2J7|A0A6P6D2J7_PTEVA KAFFEDHPHFRYAQLRFSKVKGKLRTSSLPGEDCCRYWGTRVMCLLPDGATEFAWKNKRG 364
TR|A0A6J0DKA3|A0A6J0DKA3_PERMB SDPFISSP--HILLED-----GKA----------------TVPLLAERLTTE-------- 287
TR|G5C5C9|G5C5C9_HETGA         QAFFEDHPLFRVLLEE-----GKA----------------TVPRLAERLSSE-------- 330
TR|A0A3Q0CWV5|A0A3Q0CWV5_MESAU KVFFKEHPHFRVLLED-----RKA----------------TVPHLAERLTAE-------- 315
TR|A0A6I9K5X3|A0A6I9K5X3_CHRAS QDFFEDHDYFRILLEE-----GKA----------------TIKNLAEKLSIE-------- 331
SP|P79135|MX1_BOVIN            RIFFEDHAHFRDLLEE-----GKA----------------TIPCLAERLTSE-------- 319
TR|A0A6P3H2K4|A0A6P3H2K4_BISBI RIFFEDHAHFRDLLEE-----GKA----------------TIPCLAERLTSE-------- 325
TR|L8IPR7|L8IPR7_9CETA         RIFFEDHAHFRDLLEE-----GKA----------------TIPCLAERLTSE-------- 322
SP|P27594|MX1_PIG              QAFFENHAHFRDLLEE-----GRA----------------TIPCLAERLTSE-------- 331
TR|A0A1U7U9K3|A0A1U7U9K3_CARSF KAFFEDHPHFRELLEE-----GRA----------------TVPYLADRLTTE-------- 295
SP|P33237|MX1_SHEEP            RIFFEDHTHFRDLLEE-----GRA----------------TIPCLAERLTNE-------- 325
TR|A0A7D4WPX4|A0A7D4WPX4_CAPHI RIFFEDHAHFRDLLEE-----GRA----------------TIPCLAERLTNE-------- 325
SP|Q28379|MX1_HORSE            KAFFEENPYFRGLLEE-----GRA----------------SVPCLAERLTTE-------- 330
TR|A0A6I9I6Y4|A0A6I9I6Y4_VICPA QAFFEDHTHFRDLLEE-----GRA----------------TVPRLAEKLTAE-------- 330
TR|A0A6P5BYP3|A0A6P5BYP3_BOSIN RIFFEDHAHFRDLLEE-----GKA----------------TIPCLAERLTSE-------- 322
TR|S9XPZ9|S9XPZ9_CAMFR         QAFFEDHTHFRDLLEE-----GRA----------------TVPRLAEKLTAE-------- 330
TR|A0A384DMX3|A0A384DMX3_URSMA RVFFEDHPHFRVLLEE-----GRA----------------TVPCLADKLTSE-------- 328
TR|A0A3Q7W4X3|A0A3Q7W4X3_URSAR RVFFEDHPHFRVLLEE-----GRA----------------TVPCLADKLTSE-------- 328
TR|A0A7E6D288|A0A7E6D288_9CHIR KEFFEAHPQYRELLKE-----GRA----------------TVPCLAERLTTE-------- 330
TR|S7NSW7|S7NSW7_MYOBR         RAFFEDHPYFRDLLEE-----GKA----------------TIPCLAERLTNE-------- 350
TR|A0A1N7THY9|A0A1N7THY9_PIPPI RAFFEDHPYFRDLLEE-----GKA----------------TIPCLAERLTNE-------- 326
TR|A0A1N7THZ0|A0A1N7THZ0_ROUAE KAFFEDNPYYRELLEE-----GKA----------------TIPCLAERLTAE-------- 328
TR|A0A1N7THY7|A0A1N7THY7_HYPMN KAFFADHPSYRDLLEE-----GMA----------------TIPCLAERLTAE-------- 328
TR|A0A1N7THZ1|A0A1N7THZ1_STULI KAFFQEHPQYRDLLTE-----GRA----------------TVPCLAERLTME-------- 328
TR|B2X026|B2X026_CHICK         REFFETHKHFSTLLDE-----NKA----------------TIPHLANKLTDE-------- 374
TR|A0A172PZ65|A0A172PZ65_ANSCY RQFFKTHKYFSILLDQ-----NKA----------------TIPHLAMKLTNE-------- 373
TR|A0A0Q3PUG8|A0A0Q3PUG8_AMAAE RKFFQTHKHFSILLEE-----KRA----------------TIPYLAEKLTNE-------- 254
SP|Q4ADG6|MX1_PHOVI            RDFFEDHPHFRVLLEE-----GRA----------------TVPCLADKLTSE-------- 328
SP|Q4ADG7|MX1_OTABY            RDFFEDHPHFRVLLEE-----GRA----------------TVPCLADRLTSE-------- 328
TR|A0A2Y9G772|A0A2Y9G772_NEOSC RDFFEDHPHFRVLLEE-----GRA----------------TVPCLADRLTSE-------- 328
TR|A0A6J2C7I6|A0A6J2C7I6_ZALCA RDFFEDHPHFRVLLEE-----GRA----------------TVPCLADRLTSE-------- 328
TR|A0A2U3WDI8|A0A2U3WDI8_ODORO RDFFEDHPHFRVLLEE-----GRA----------------TVPCLADKLTSE-------- 328
TR|A0A1U7SNC8|A0A1U7SNC8_ALLSI RKFFENHKYFGDLLKE-----KKA----------------TIPLLAEKLTNE-------- 401
TR|A0A7M4FFF4|A0A7M4FFF4_CROPO RKFFQDHEYFSVLLDE-----RKA----------------TIPLLAEKLTAE-------- 309
TR|A0A340WTU5|A0A340WTU5_LIPVE QDFSGNHAHFRDLLEE-----GRA----------------TVPCLAERLTAE-------- 335
TR|A0A6J3RAX5|A0A6J3RAX5_TURTR QDFFGNHAHFRDLLEE-----GRA----------------TVPCLAERLTTE-------- 330
TR|A0A341CI24|A0A341CI24_NEOAA QDFFGNHAHFRDLLEE-----GRA----------------TAPCLAERLT-X-------- 410
TR|A0A2Y9LTL6|A0A2Y9LTL6_DELLE QDFFGNHAHFRDLLEE-----GRA----------------TVPCLAERLTTE-------- 334
TR|A0A383ZCI9|A0A383ZCI9_BALAS QDFFENHAHFRDLLEE-----GRA----------------TVPCLAERLTSE-------- 329
TR|V9KIB1|V9KIB1_CALMI         KVFFEEHQHFRFLIDE-----GKA----------------SIPCLAKRLTSE-------- 294
TR|A0A674K7E1|A0A674K7E1_TERCA REFFEEHEYFSILLEE-----RKA----------------TIPLLAEKLTQE-------- 336
TR|A0A4D9DW37|A0A4D9DW37_9SAUR RLYSYSLFCFRVLLDE-----NKA----------------TIPILAEKLTTE-------- 443
TR|H2DQ54|H2DQ54_XENLA         QNFFKEHEHFSVLLEE-----GNA----------------TIACLAEKLTNE-------- 294
TR|A0A0F6MUS9|A0A0F6MUS9_ANGJA KAFFEDHAHFSTLLDE-----GFA----------------TIPKLAEKLTLE-------- 309
TR|Q98990|Q98990_SALSA         KAFFKEHAHLSTLYDE-----GHA----------------TIPKLAEKLTLE-------- 294
TR|A0A674EGU5|A0A674EGU5_SALTR KAFFKEHAHLSTLYDE-----GHA----------------TIPKLAEKLTLE-------- 294
SP|Q91192|MX1_ONCMY            KAFFKEHAHLSTLYDE-----GHA----------------TIPKLAEKLTLE-------- 294
TR|A0A286K200|A0A286K200_MYLPI NAFFQDHPQFSKLYEE-----GFA----------------TIPKLAEKLTIE-------- 297
TR|A0A498MHV8|A0A498MHV8_LABRO NAFFKDHPHFSKLYEE-----GFA----------------TIPKLAEKLTIE-------- 298
SP|Q7T2P0|MX1_ICTPU            KDFFKDHPHFSTLYEE-----GMA----------------TIPNLAEKLTLE-------- 299
TR|A0A556TK66|A0A556TK66_BAGYA -----------ALYDN-----GLA----------------TIPKLAEKLTLE-------- 267
TR|A0A3G6IID3|A0A3G6IID3_9TELE SAFFQDHPQFSKLYEE-----GFA----------------TIPKLAEKLTIE-------- 297
SP|Q8JH68|MXA_DANRE            SAFFKDHPHFSKLYEE-----GFA----------------TIPKLAEKLTIE-------- 297
TR|A0A672MEY2|A0A672MEY2_SINGR LKFFRNHHYFSSLLDD-----GKA----------------STQCLAAKLTKE-------- 296
TR|Q7T2M3|Q7T2M3_CARAU         NAFFQDHPQFSKLYEE-----GFA----------------TIPKLAEKLTIE-------- 297
TR|A0A673LWX8|A0A673LWX8_9TELE DAFFKDHPHFSKLYDE-----GFA----------------TIPKLAEKLTIE-------- 297
TR|A0A8B9KF29|A0A8B9KF29_ASTMX KAFFRDHPHFSALYDD-----GKA----------------TVPKLAEKLTLE-------- 294
                                            *    :
```

```
SP|P20591|MX1_HUMAN            -LITHICKSLP-------LLENQIKETHQRITEELQKYGVDIPEDENEKMFFL------- 375
TR|K4JBQ0|K4JBQ0_9PRIM         -LITHICKSLP-------LLENQIKESHQRITEELQKYGVDIPEDENEKMFFL------- 375
TR|H2QL18|H2QL18_PANTR         -LITHICKSLP-------LLENQIKESHQRITEELQKYGVDIPEDENEKMFFL------- 375
SP|A1E2I4|MX1_MACMU            -LIAHICKSLP-------LLENQIKESHQGITEELQKYGVDIPEDENEKMFFL------- 375
SP|Q5R5G3|MX1_PONAB            -LITHICKSLP-------LLENQIRESHQRITEELQKYGVDVPEDENEKMFFL------- 375
TR|A0A6P8QPW8|A0A6P8QPW8_GEOSA -LVEHIHKSLP-------DLVQQINSKLKESTEELEHFGTGIPESNEGMYNFL------- 339
TR|U3EGN4|U3EGN4_CALJA         -LITHICKSLP-------LLENQIKESHQKITEELQKYGVDIPEDENEKMFFL------- 375
SP|Q9N0Y3|MX1_CANLF            -LITHICKTLP-------LLENQIKENHEKITEELQKYGSDVPEDEHEKMFFL------- 371
TR|M3W8X0|M3W8X0_FELCA         -LIMHICKSLP-------LLENQIKENHEKITEELQKYGSDVPEDEHEKMFFL------- 373
TR|A0A667GNL8|A0A667GNL8_LYNCA -LIMHICKSLP-------LLENQIKENHEKITEELQKYGSDVPEDEHEKMFFL------- 373
TR|A0A6P6GZN4|A0A6P6GZN4_PUMCO -LIMHICKSLP-------LLENQIKENHEKITEELQKYGSDVPEDEHEKMFFL------- 373
TR|A0A6J1XXY6|A0A6J1XXY6_ACIJB -LIMHICKSLP-------LLENQIKENHEKITEELQKYGSDVPEDEHEKMFFL------- 373
TR|A0A6P4TS13|A0A6P4TS13_PANPR -LIMHICKSLP-------LLENQIKENHEKITEELQKYGSDVPEDEHEKMFFL------- 411
TR|A0A3Q7SCG7|A0A3Q7SCG7_VULVU -LITHICKTLP-------LLENQIKENHEKITEELQKYGSDVPEDEHEKMFFL------- 409
TR|I3MJX6|I3MJX6_ICTTR         -LIAHICKSLP-------LLENQIKENIQKVTEELKNFGVDVPEEESLKTFFL------- 376
TR|G9KBY2|G9KBY2_MUSPF         -LIMHICKTLP-------LLENQIKENHEKITEELQKYGSDVPEDEHEKMFFL------- 370
TR|A0A2Y9KR88|A0A2Y9KR88_ENHLU -LIMHICKTLP-------LLENQIKENHEKITEELQKYGSDVPEDEHEKMFFL------- 370
SP|P09922|MX1B_MOUSE           -LTSHICKSLP-------LLEDQINSSHQSASEELQKYGADIPEDDRTRMSFL------- 341
SP|P18588|MX1_RAT              -LTSHICKSLP-------ILENQINVNHQIASEELQKYGADIPEDDSKRLSFL------- 366
TR|A0A6P6D2J7|A0A6P6D2J7_PTEVA VHDYGEKKSLP-------QLENEIKEKQQSITEELQMYGMETPEDENEKMFFL------- 410
TR|A0A6J0DKA3|A0A6J0DKA3_PERMB -LISHICKSLP-------LLENQINKSHQSTIEELQKYGIDIPEDSNKRMFFL------- 332
TR|G5C5C9|G5C5C9_HETGA         -LITHICKSLP-------LLENTIKERHQEITEELQKYGMDIPEDGNEKTIFL------- 375
TR|A0A3Q0CWV5|A0A3Q0CWV5_MESAU -LISHICKSLP-------LLENQIKESHQRASEELQKYGTDVPEDSNERTFFL------- 360
TR|A0A6I9K5X3|A0A6I9K5X3_CHRAS -LIIHISKSLP-------LLEKQIKESHQHITEQLQNYGTDIPQDESEKMFFL------- 376
SP|P79135|MX1_BOVIN            -LIMHICKTLP-------LLENQIKETHQRITEELQKYGKDIPEEESEKMFCL------- 364
TR|A0A6P3H2K4|A0A6P3H2K4_BISBI -LIMHICKTLP-------LLENQIKETHQRITEELQKYGKDIPEEESEKMFCL------- 370
TR|L8IPR7|L8IPR7_9CETA         -LIMHICKTLP-------LLENQIKETHQRITEELQKYGKDIPEEESEKMFCL------- 367
SP|P27594|MX1_PIG              -LIMHICKTLP-------LLENQIKESHQKITEELQKYGSDIPEDESGKMFFL------- 376
TR|A0A1U7U9K3|A0A1U7U9K3_CARSF -LITHISKSMP-------LLEEQIKVNHQKITEDLQKYGVDIPEDETEKMFFL------- 340
SP|P33237|MX1_SHEEP            -LIMHICKTLP-------LLENQIKETHQRITEELQKYGKDIPEEESEKMFSL------- 370
TR|A0A7D4WPX4|A0A7D4WPX4_CAPHI -LIMHICKTLP-------LLENQIKETHQRITEELQKYGKDIPEEESEKMFSL------- 370
SP|Q28379|MX1_HORSE            -LITHISKSLP-------LLENQIKESYQNLSDELQKYGTDIPEDETEKTFFL------- 375
TR|A0A6I9I6Y4|A0A6I9I6Y4_VICPA -LITHICKSLP-------LLENQIKESHQKISEELQKCGSDIPEDESGKMFLL------- 375
TR|A0A6P5BYP3|A0A6P5BYP3_BOSIN -LIMHICKTLP-------LLENQIKETHQRITEELQKYGKDIPEEESEKMFCL------- 367
TR|S9XPZ9|S9XPZ9_CAMFR         -LITHICKSLP-------LLENQIKESHQKISEELQKCGSDIPEDKSGKMFLL------- 375
TR|A0A384DMX3|A0A384DMX3_URSMA -LISHICKTLP-------LLENQIKENHEKITEELQKYGSDVPEDEHEKMFFL------- 373
TR|A0A3Q7W4X3|A0A3Q7W4X3_URSAR -LISHICKTLP-------LLENQIKENHEKITEELQKYGSDVPEDEHEKMFFL------- 373
TR|A0A7E6D288|A0A7E6D288_9CHIR -LIEHICKSLP-------KLENQIKEKKEKVTEDLQKCGTDIPEEEGEKMIFL------- 375
TR|S7NSW7|S7NSW7_MYOBR         -LIAHISKSLP-------LLENQIKENQQSITEQLQKYGMDIPEEETEKMFFL------- 395
TR|A0A1N7THY9|A0A1N7THY9_PIPPI -LIAHISKSLP-------LLESQIKENQQTITEQLQKYGLDIPEDETEKMFFL------- 371
TR|A0A1N7THZ0|A0A1N7THZ0_ROUAE -LIAHICKSLP-------RLENEIKEKQQTITEELQKYGTEIPEDESEKMFFL------- 373
TR|A0A1N7THY7|A0A1N7THY7_HYPMN -LIAHICKSLP-------WLENEIKEKQQNITEELQKYGTEIPEDESEKMFFL------- 373
TR|A0A1N7THZ1|A0A1N7THZ1_STULI -LTEHISKSLP-------QLEKQIKEKQQKVTEDLQKCGMDIPEEESEKVIFL------- 373
TR|B2X026|B2X026_CHICK         -LVGRIIKTLP-------AIEKQVHDALQQAKKELQKYTQSTHPTVSDKTIFL------- 419
TR|A0A172PZ65|A0A172PZ65_ANSCY -LVAHIIKTLP-------TIENQIREVLQKSVQELQKYRRGTPTIETEKLAFL------- 418
TR|A0A0Q3PUG8|A0A0Q3PUG8_AMAAE -LVGHIIDL--------------------------EKYRKVXPITESEKLTVLTDVSINI 287
SP|Q4ADG6|MX1_PHOVI            -LITHICKTLP-------LLENQIKENHEKITEELKKYGSDVPEEEHEKMFFL------- 373
SP|Q4ADG7|MX1_OTABY            -LITHICKTLP-------LLEKQIKENYEKITEELQKYGSDVPEEEHEKMFFL------- 373
TR|A0A2Y9G772|A0A2Y9G772_NEOSC -LITHICKTLP-------LLENQIKENHEKITQELQKYGSDVPEEEHEKMFFL------- 373
TR|A0A6J2C7I6|A0A6J2C7I6_ZALCA -LITHICKTLP-------LLENQIKENYEKITEELQKYGSDVPEEEHEKMFFL------- 373
TR|A0A2U3WDI8|A0A2U3WDI8_ODORO -LITHICKTLP-------LLENQIKENHEKITEELQKYGSDVPEEDHEKMFFL------- 373
TR|A0A1U7SNC8|A0A1U7SNC8_ALLSI -LVIHISRSLP-------ALEERVSEELRKAIEKLQKNCGSVPEEESEKMMLL------- 446
TR|A0A7M4FFF4|A0A7M4FFF4_CROPO -LVFHIYKNIP-------RLEDQISEELQKSSQELRKYGKGTPRTDGEKLFFL------- 354
TR|A0A340WTU5|A0A340WTU5_LIPVE -LIMHFCKSLP-------LLESQIKGSHQRITEELQKYDPDTPEDESGKMFSL------- 380
TR|A0A6J3RAX5|A0A6J3RAX5_TURTR -LITHICKSLP-------LLENQIKESHQRITEELQKYGSDIPEDASGKMFSL------- 375
TR|A0A341CI24|A0A341CI24_NEOAA -LIMHTCKSLP-------LLENQIKESHQRITEELQXYGSDIPEDASGKMFSL------- 455
TR|A0A2Y9LTL6|A0A2Y9LTL6_DELLE -LIVHICKSLP-------LLENQIKESHQRITEELQXYGSDIPEDASGKMFSL------- 379
TR|A0A383ZCI9|A0A383ZCI9_BALAS -LITHICKSLP-------LLENQIKESHQRITEELQKYGVDIPEDESGKMFFL------- 374
TR|V9KIB1|V9KIB1_CALMI         -LVDHIKKWLP-------KFEKELSDRIYAITQELKYYGEGVPDTENEKITFL------- 339
TR|A0A674K7E1|A0A674K7E1_TERCA -LIEHINKSLP-------TLENQIKDKLRVANNKLQKYGKGVPESVEEQMLFL------- 381
TR|A0A4D9DW37|A0A4D9DW37_9SAUR -LVEHINKSLP-------ALEEQINIQLQKATEELLKYGKGTPKAEDEKLFFL------- 488
TR|H2DQ54|H2DQ54_XENLA         -LVDHIVKTLP-------ALDTQIRKKLNEAEEKLRNLGTSVPDSETEKLRFL------- 339
TR|A0A0F6MUS9|A0A0F6MUS9_ANGJA -LVHHIEKSLP-------RLEQQIETKLDDTRAELDKYGKGPPEDPAERMCFL------- 354
TR|Q98990|Q98990_SALSA         -LVHHIEKSLP-------RLEEQIEAKLAETHAELERYGTGPPEDSAERMYFL------- 339
TR|A0A674EGU5|A0A674EGU5_SALTR -LVHHIEVNSSSITFHSMTLRCQIEAKLAETHAELERYGTGPPEDSAERMYFL------- 346
SP|Q91192|MX1_ONCMY            -LVHHIEKSLP-------RLEEQIEAKLSETHAELERYGTGPPEDSAERLYFL------- 339
TR|A0A286K200|A0A286K200_MYLPI -LVHHIQKSLP-------RLEEQIEAKLAETQKELEAYGNGPPSDPAERLSFF------- 342
TR|A0A498MHV8|A0A498MHV8_LABRO -LVHHIQKSLP-------RLEEQIETKLAETMKELEAYGNGPPSDPAERLSFL------- 343
SP|Q7T2P0|MX1_ICTPU            -LVHHIELSLP-------RLEEQIDIKLADSQAELDRYGSGPPTEPAERICFL------- 344
TR|A0A556TK66|A0A556TK66_BAGYA -LVHHIELSLP-------RLEEQIEVKLAETQCELDRYGSGPPTEPGDRIVFL------- 312
TR|A0A3G6IID3|A0A3G6IID3_9TELE -LVHHIQKSLP-------RLEEQIEAKLAETQKELEAYGNGPPSDPAERLSFF------- 342
SP|Q8JH68|MXA_DANRE            -LVHHIQKSLP-------RLEEQIETKLAETQKELEAYGNGPPSEPAARLSFF------- 342
TR|A0A672MEY2|A0A672MEY2_SINGR -LADHIKKSLP-------SLTEQIQTRLLNVQRDLKNYEQGPPLEEEMMGPFLSEVR--- 345
TR|Q7T2M3|Q7T2M3_CARAU         -LVHHIQKSLP-------RLEEQIETKLAETQKELEAYGNGPPSDPVVRLSFL------- 342
TR|A0A673LWX8|A0A673LWX8_9TELE -LVHHIQKSLP-------RLEEQIETKLAETQKELEAYGNGPPSDPAERLSFL------- 342
TR|A0A8B9KF29|A0A8B9KF29_ASTMX -LVHHIEKSLP-------RLEEQIEAKLAETQDELEKYGNGPPMEPSERIYYL------- 339
                                                                                   :

SP|P20591|MX1_HUMAN            -------IDKVNAFNQDITALMQGEETVG-EEDIRLFTRLRHEFHKWSTIIENNFQEG-- 425
TR|K4JBQ0|K4JBQ0_9PRIM         -------IDKINAFNQDITALMQGEETVG-EEDIRLFTRLRHEFHKWSTIIENNFQEG-- 425
TR|H2QL18|H2QL18_PANTR         -------IDKVNAFNQDISALMQGEETVG-EEDIRLFTRLRHEFHKWSTIIENNFQEG-- 425
SP|A1E2I4|MX1_MACMU            -------IDKINAFNQDITALIQGEETVG-EDDSRLFTRLRREFHKWGIIIENNLQEG-- 425
SP|Q5R5G3|MX1_PONAB            -------IDKINAFNQDITALIQGEETVG-EEDIRLFTRLRHEFHKWSIIIENNFQEG-- 425
TR|A0A6P8QPW8|A0A6P8QPW8_GEOSA -------LDKIRPFNKEILSLIKGEENLSIPNTSRLFKIVRQNFELWQKLLDDNSILM-- 390
TR|U3EGN4|U3EGN4_CALJA         -------IDKINAFNQDIAAFLHGEETVG-EEDVRLFTKLRNEFRKWSVVIENNFQES-- 425
SP|Q9N0Y3|MX1_CANLF            -------IDKLNAFNQDISSLIQGEESVG-EDESRLFTKIRNEFHKWSAVIEKKFQRG-- 421
TR|M3W8X0|M3W8X0_FELCA         -------IDKINAFNHDITSLIQGEESVG-EGESRLFTKIRHEFHKWSTVIEKKFQKG-- 423
TR|A0A667GNL8|A0A667GNL8_LYNCA -------IDKINAFNHDITSLIQGEESVG-EGESRLFTKIRHEFHKWSTVIEKKFQKG-- 423
```

```
TR|A0A6P6GZN4|A0A6P6GZN4_PUMCO -------IDKINAFNHDITSLIQGEESVG-EGESRLFTKIRHEFHKWSTVIEKKFQKG-- 423
TR|A0A6J1XXY6|A0A6J1XXY6_ACIJB -------IDKINAFNHDITSLIQGEESVG-EGESRLFTKIRHEFHKWSTVIEKKFQKG-- 423
TR|A0A6P4TS13|A0A6P4TS13_PANPR -------IDKINAFNHDITSLIQGEESVG-EGESRLFTKIRHEFHKWSIVIEKKFQKG-- 461
TR|A0A3Q7SCG7|A0A3Q7SCG7_VULVU -------IDKLNAFNQDINSLIQGEESVG-EDESRLFTKIRNEFHKWSAVIEKKFQRG-- 459
TR|I3MJX6|I3MJX6_ICTTR         -------IDKINQFNQDINAIVQAEEVVG-PEEMRLFTRMRKEFHKWSNEIEENFQKG-- 426
TR|G9KBY2|G9KBY2_MUSPF         -------IDKINAFSHDINSLIEGEESVG-ENDSRLFTKIRNEFHKWNTVIEKAFQKG-- 420
TR|A0A2Y9KR88|A0A2Y9KR88_ENHLU -------VDKINAFSQDIKSLIEGEESVG-ENESRLFTKIRNEFHKWSTVIEKTFQKG-- 420
SP|P09922|MX1B_MOUSE           -------VNKISAFNRNIMNLIQAQETVS-EGDSRLFTKLRNEFLAWDDHIEEYFKKD-- 391
SP|P18588|MX1_RAT              -------MNKINVFNKDILSLVQAQENIS-WEESRLFTKLRNEFLAWNDYIEEHFKKTLG 418
TR|A0A6P6D2J7|A0A6P6D2J7_PTEVA -------IDKINAFNQGILTLIQGEEIVD-KSNTRLFTKIRKEFNNWSKVIENNFQDG-- 460
TR|A0A6J0DKA3|A0A6J0DKA3_PERMB -------IDKINAFNKDITALVKAQETVS-GGGSWLFTNLRKEFLNWDNYIEEHFKKS-- 382
TR|G5C5C9|G5C5C9_HETGA         -------IQKISAFNKDIRALVEGEESVG-KNGTRLFARLRQKFQGWNSMIEEHFEKG-- 425
TR|A0A3Q0CWV5|A0A3Q0CWV5_MESAU -------IDKINAFNKDITALVQAQETVP-AGGSRLFTNMRNKFLDWNNYIDEHLKKS-- 410
TR|A0A6I9K5X3|A0A6I9K5X3_CHRAS -------IDKISAFNQDILALVQGEELVK-KEESRLFTKIRHQFSEWSDKIEKNFQEG-- 426
SP|P79135|MX1_BOVIN            -------IEKIDTFNKEIISTIEGEEFVE-QYDSRLFTKVRAEFSKWSAVVEKNFEKG-- 414
TR|A0A6P3H2K4|A0A6P3H2K4_BISBI -------IEKIDTFNKEIISTIEGEEFVE-QYDSRLFTKVRAEFSKWSAVVEKNFEKG-- 420
TR|L8IPR7|L8IPR7_9CETA         -------IEKIDTFNKEIISTIEGEEFVE-QYDSRLFTKVRAEFSKWSAVVEKNFEKG-- 417
SP|P27594|MX1_PIG              -------IDKIDAFNSDITALIQGEELVV-EYECRLFTKMRNEFCRWSAVVEKNFKNG-- 426
TR|A0A1U7U9K3|A0A1U7U9K3_CARSF -------IDKIKAFNQDIAALVQGEETVE-DEDTRLFTKLRNEFQKWSVIIEKNFKSS-- 390
SP|P33237|MX1_SHEEP            -------IEKIDTFNKEIISTIEGEEHVG-QYDSRLFTKVRAEFCKWSAVVEKNFEKG-- 420
TR|A0A7D4WPX4|A0A7D4WPX4_CAPHI -------IEKIDTFNKEIISTIEGEELVG-QYDSRLFTKVRAEFCKWSAVVEKNFEKG-- 420
SP|Q28379|MX1_HORSE            -------IVKITTFNQNITSFVQGEELVG-PNDTRLFNKIRQEFQKWSGVIENNFRKG-- 425
TR|A0A6I9I6Y4|A0A6I9I6Y4_VICPA -------IEKIDAFNKEITALIQGEECVG-EHKSRLFTRIRSEFSKWSTVIEKHFQKG-- 425
TR|A0A6P5BYP3|A0A6P5BYP3_BOSIN -------IEKIDTFNKEIISTIEGEEFVE-QYDSRLFTKVRAEFSKWSAVVEKNFEKG-- 417
TR|S9XPZ9|S9XPZ9_CAMFR         -------IEKIDAFNKEITALIQGEECVG-EHKSRLFTRIRSYDDICK------------ 415
TR|A0A384DMX3|A0A384DMX3_URSMA -------IDKINAFNHDINSLIEGEESVG-EDESRLFTKIRTEFHKWSAVIEKSFQKG-- 423
TR|A0A3Q7W4X3|A0A3Q7W4X3_URSAR -------IDKINAFNHDINSLIEGEESVG-EDESRLFTKIRTEFHKWSAVIEKRFQKG-- 423
TR|A0A7E6D288|A0A7E6D288_9CHIR -------IEKIKAFNEDIRTLIDGEEFVE-EGDTRMFNKIRNEFCRWSQKVHASFQKS-- 425
TR|S7NSW7|S7NSW7_MYOBR         -------IDKINTFNQDIQALVEGEESVC-GDNSRLFTRIRMEFGKWSIEIEKSFQRG-- 445
TR|A0A1N7THY9|A0A1N7THY9_PIPPI -------IDRINTFNQDIEALVEGEESVG-GDDSRLFTRIRLEFCKWGLEIENNFQRG-- 421
TR|A0A1N7THZ0|A0A1N7THZ0_ROUAE -------IEKINAFNQGIRALTQAEELVG-KNDTRLFTKIRNEFNNWSKLIEKSFQMG-- 423
TR|A0A1N7THY7|A0A1N7THY7_HYPMN -------IDKINAFNQGIALTQAEELVG-KNDTRLFTKIRNEFNSWSKMIDRSFQAG-- 423
TR|A0A1N7THZ1|A0A1N7THZ1_STULI -------IEKIQAFNEDIRTLIDGEEFVE-EDDTRMFNKVRSEFYNWSLIVHSSFKKG-- 423
TR|B2X026|B2X026_CHICK         -------VGLIKAFNEDISQTMHGKESWF-GNEIRLFPKIRREFRTWGVKLLESSAKV-- 469
TR|A0A172PZ65|A0A172PZ65_ANSCY -------TDLIKLFNQDVSRVIRGEEHLL-GNENRLFAKLRKEFQTWGLILMENAAKV-- 468
TR|A0A0Q3PUG8|A0A0Q3PUG8_AMAAE DLVTEQNEELIKLFNQDVSRTICGEEQLV-GNEIRLFTXIRKEFQTWEAVLLECAV---- 342
SP|Q4ADG6|MX1_PHOVI            -------IEKINAFNHDINSLIEGEEFVG-EDESRLFTKIRNEFHKWSCVIEKKFQQG-- 423
SP|Q4ADG7|MX1_OTABY            -------IEKINAFNHDITSLTEGEEFVG-EDECRLFTKIRNEFHKWSLVIEKRFQQG-- 423
TR|A0A2Y9G772|A0A2Y9G772_NEOSC -------IEKINAFNHDINSLIEGEEFVG-EDESRLFTKIRNEFHKWSSVIEKKFQQG-- 423
TR|A0A6J2C7I6|A0A6J2C7I6_ZALCA -------IEKINAFNHDITSLTEGEEFVG-EDECRLFTKIRNEFHKWSLVIEKRFQRG-- 423
TR|A0A2U3WDI8|A0A2U3WDI8_ODORO -------IEKINAFNHDITSLIKGEEFVG-EDECRLFTKIRNEFHKWSFVIEKKFQQG-- 423
TR|A0A1U7SNC8|A0A1U7SNC8_ALLSI -------IKKIKCFNTDITNLIRGEEDLYE-TEIRMFSKIRKEFQEWGRILDVRSDKV-- 496
TR|A0A7M4FFF4|A0A7M4FFF4_CROPO -------IDKIKLFNQDIVNSAHGEEKVSG-NESRLFTKVRKEFQKWGKALDESSVKV-- 404
TR|A0A340WTU5|A0A340WTU5_LIPVE -------IDKIDAFNKEITDVIEGEEFVG-EYDSRLFNKMRTEFYGWSVVVENGFQKG-- 430
TR|A0A6J3RAX5|A0A6J3RAX5_TURTR -------IDKIDAFNKEITNIIEGEEFVG-ERDSRLFNKMRTEFYRWSIVVANSFRKG-- 425
TR|A0A341CI24|A0A341CI24_NEOAA -------TDKIDAFNKEITDIIEGEEFVG-EYDSRLFNKMRTEFYRWSIVVANSFRKG-- 505
TR|A0A2Y9LTL6|A0A2Y9LTL6_DELLE -------TDKIDAFNKDITNIIEGEEFVG-EYDSRLFNKMRTEFYRWSIVVANNFWKG-- 429
TR|A0A383ZCI9|A0A383ZCI9_BALAS -------IDKIDAFNKEITNKIEGEEFVG-EYDSRLFNKMRTEFYKWSIVVEKNFQKG-- 424
TR|V9KIB1|V9KIB1_CALMI         -------IQKINTFGQHVVSLALGEEPETFIQKERCYSKVRKEFQQWKVYLNHALHQF-- 390
TR|A0A674K7E1|A0A674K7E1_TERCA -------VDKLKLFNQDIMNSVQGEEILTDKTKQRLFTKVRKYFNEWEAHINNSTLKV-- 432
TR|A0A4D9DW37|A0A4D9DW37_9SAUR -------IDKIKLFNQDIVSSVVGEEKLFE-NEIRLFTKIRTEFQKWGKILDDSALTD-- 538
TR|H2DQ54|H2DQ54_XENLA         -------IDKITRFAGAIINATQGDEEMTN-GSLKLFTVIRKHFYFWESFIQTQSKSF-- 389
TR|A0A0F6MUS9|A0A0F6MUS9_ANGJA -------IDKVTAFTQDITNLTLGEELRN-GYKMNLFSQLRSEFGKWKACLDRSGDKF-- 404
TR|Q98990|Q98990_SALSA         -------IDKVTAFTHDAINLSTGEELKS-GVRLNVFSTLRKEFGKWKLHLDHSGENF-- 389
TR|A0A674EGU5|A0A674EGU5_SALTR -------IDKVTAFTQDAINLSTGEELKS-GVRLNVFSTLRQEFGKWKLHLDHSGENF-- 396
SP|Q91192|MX1_ONCMY            -------IDKVTAFTQDAINLSTGEEMKS-GVRLNVFSTLRKEFGKWKLHLERSGEIF-- 389
TR|A0A286K200|A0A286K200_MYLPI -------IDKVTSFIQDTLNLTTGEEIKC-GSDLLIFPELRNEFAKWNSFLDHSGYLF-- 392
TR|A0A498MHV8|A0A498MHV8_LABRO -------IDKVTAFTHDMFNLTTGEEVKC-ASDLLIFPELRREFAKWNSLLDRSGDSF-- 393
SP|Q7T2P0|MX1_ICTPU            -------IDKVTAFTQDTINLTTGEELKN-IQHLNIFSSLRRQFALWKMHLDDSGETF-- 394
TR|A0A556TK66|A0A556TK66_BAGYA -------IDKVTAFTQDAINLTNGEELKN-IQHLNIFSSLRRQFVYWKTHLDKSGKTF-- 362
TR|A0A3G6IID3|A0A3G6IID3_9TELE -------IDKVTSFIQDTLNLTIGEEIKC-GSDLLIFPELRSEFAKWNGFLDHSGYLF-- 392
SP|Q8JH68|MXA_DANRE            -------IDKVTAFNQDMLNLTTGEDVKC-TTDLLLFPELRQEFAKWSHILDRSGDSF-- 392
TR|A0A672MEY2|A0A672MEY2_SINGR ----------ALRFSDQINELSR----TGHSKDKNIYISLRPVFKKWDIYLRSTEVSF-- 389
TR|Q7T2M3|Q7T2M3_CARAU         -------IDKVTAFTHDMFNLTTGEEVKC-SSDLLVFPELREEFLKWNGFLERSGHSF-- 392
TR|A0A673LWX8|A0A673LWX8_9TELE -------IDKVTAFTHDMFNLTTGEEVKC-ASDLLVFPELRDEFAEWNDVLYRSGVSF-- 392
TR|A0A8B9KF29|A0A8B9KF29_ASTMX -------IDKVTAFTQDALNLTMGEELRT-APHLNIFSNLRIKFESWKQHLDQSGETF-- 389
                                            *                         :  :*

SP|P20591|MX1_HUMAN            HKILSR---KIQKFENQYRGRELPGFVNYRTFETIVKQQIKALEEPAVDMLHTVTDMVRL 482
TR|K4JBQ0|K4JBQ0_9PRIM         HKILSR---KIQKFENQYRGRELPGFVNYRTFETIVKQQIKALEEPAVDMLHTVADMVRL 482
TR|H2QL18|H2QL18_PANTR         HKILSR---KIQKFENQYRGRELPGFVNYRTFETIVKQQIKALEEPAVDMLHTVTDMVRL 482
SP|A1E2I4|MX1_MACMU            HKITSR---KMQKFENQYRGRELPGFVNYRTFETIVKQQIKALEEPAVNMLHTVTDMVRL 482
SP|Q5R5G3|MX1_PONAB            HKILSR---KIQKFENQYRGRGLPGFVNYRTFETIVKQQIKALEEPAVDMLHTVTDMVRL 482
TR|A0A6P8QPW8|A0A6P8QPW8_GEOSA DKTVKD---KLLEFDGKYRGRELPGFINYQTFEMIVNDYIQKLRMSALENVKSLIHIIED 447
TR|U3EGN4|U3EGN4_CALJA         HEIMSR---KIQKFENQYRGRELPGFVNYRTFETIVKQQIKALEEPAVDMLHTVTDMVRL 482
SP|Q9N0Y3|MX1_CANLF            YKAIYK---QMEKFENRYRGRELPGFVNYKTFEIIIKQQIKELEEPAVDMLHTITDMVQV 478
TR|M3W8X0|M3W8X0_FELCA         YKAIYK---EIEKFENRYRGRELPGFVNYKTFEIIIKQQIRELEEPAVDMLRTVTDMVQL 480
TR|A0A667GNL8|A0A667GNL8_LYNCA YKAIYK---EIEKFENRYRGRELPGFVNYKTFEIIIKQQIRELEEPAVDMLRTVTDMVQL 480
TR|A0A6P6GZN4|A0A6P6GZN4_PUMCO YKAIYK---EIEKFENRYRGRELPGFVNYKTFEIIIKQQIRELEEPAVDMLRTVTDMVQL 480
TR|A0A6J1XXY6|A0A6J1XXY6_ACIJB YKAIYK---EIEKFENRYRGRELPGFVNYKTFEIIIKQQIRELEEPAVDMLRTVTDMVQL 480
TR|A0A6P4TS13|A0A6P4TS13_PANPR YKAIYK---EIEKFENRYRGRELPGFVNYKTFEIIIKQQIRELEEPAVDMLRTVTDMVQL 518
TR|A0A3Q7SCG7|A0A3Q7SCG7_VULVU YKAIYK---QMEKFENRYRGRELPGFVNYKTFEIIIKQQIKELEEPAVDMLHTITDMVQV 516
TR|I3MJX6|I3MJX6_ICTTR         HGVLRK---EIRRFENQYRGRELPGFVNYRTFETIIKEQLRSLEEPAVDMLHTVTDMVRV 483
TR|G9KBY2|G9KBY2_MUSPF         YKAIYK---QIEKFENRYRGRELPGFVNYKTFEIIIKQQIKELEEPAVDMLHTVTDMVQA 477
TR|A0A2Y9KR88|A0A2Y9KR88_ENHLU YQAIYK---PMEKFENRYRGRELPGFVNYKTFEIIIKQQIKELEEPAVDMLHTVMDMVQA 477
SP|P09922|MX1B_MOUSE           SPEVQS---KMKEFENQYRGRELPGFVDYKAFESIIKKRVKALEESAVNMLRRVTKMVQT 448
SP|P18588|MX1_RAT              SSEKHS---QMEKFESHYRGRELPGFVDYKAFENIIKKEVKALEEPALNMLHRVTTMVKN 475
TR|A0A6P6D2J7|A0A6P6D2J7_PTEVA YDAVYN---EICKFENQYRGRELPGFVNYKTFENIIKQQIKTLEEPAVDMLHIVTQMVWL 517
```

```
TR|A0A6J0DKA3|A0A6J0DKA3_PERMB SAEIDR---QIQVFENQYHGRELPGFVNYKTFESIIKNPIKALEEXAVVLLHKVTDMVRS 439
TR|G5C5C9|G5C5C9_HETGA         YDGLKS---KVWQFESQHCGKELPGFVNYRTFETLMKSQLQALEEPAVELLHEVTVMVRD 482
TR|A0A3Q0CWV5|A0A3Q0CWV5_MESAU SVQLYS---QIQVFENQYRGRELPGFVNYKTFENIIRKQIKALEEPAVVMLHEVTNMVRV 467
TR|A0A6I9K5X3|A0A6I9K5X3_CHRAS YRSMCM---EIYKFEKQYRGRELPGFVNYRTFEVIMKEQIKKLEEPAVEMLHTVTDIVRL 483
SP|P79135|MX1_BOVIN            YEAIRK---EIKQFENRYRGRELPGFVNYKTFETIIKKQVRVLEEPAVDMLHTVTDIIRN 471
TR|A0A6P3H2K4|A0A6P3H2K4_BISBI YEAIRK---EIKQFENRYRGRELPGFVNYKTFETIIKKQVRVLEEPAVDMLHTVTDIIRN 477
TR|L8IPR7|L8IPR7_9CETA         YEAIRK---EIKQFENRYRGRELPGFVNYKTFETIIKKQVRVLEEPAVDMLHTVTDIIRN 474
SP|P27594|MX1_PIG              YDAICK---QIQLFENQYRGRELPGFVNYKTFETIIKKQVSVLEEPAVDMLHTVTDLVRL 483
TR|A0A1U7U9K3|A0A1U7U9K3_CARSF YDLMSN---KIHKFENQYRGRELPGFVNYRTFESIIKQQIKTLEEPAVDMLHRVTDIVRL 447
SP|P33237|MX1_SHEEP            HEAIRK---EIKQFENRYRGRELPGFVNYKTFEIIIKKQVIVLEEPAVDMLHTVTDIIRN 477
TR|A0A7D4WPX4|A0A7D4WPX4_CAPHI HEAIRK---EIKQFENRYRGRELPGFVNYKTFEIIIKKQVIVLEEPAVDMLHTVTDIIRN 477
SP|Q28379|MX1_HORSE            GEAIRR---QIWTFENQYRGRELPGFVNYRTFETIIKQQIQLLEEPAIDMLHRISDLVRD 482
TR|A0A6I9I6Y4|A0A6I9I6Y4_VICPA EPQATRETWQGARTQGGYRGRELPGFVNYKTFETIIKKQVEILEEPAVDMLHRVTDIVRL 485
TR|A0A6P5BYP3|A0A6P5BYP3_BOSIN YEAIRK---EIKQFENRYRGRELPGFVNYKTFETIIKKQVRVLEEPAVDMLHTVTDIIRN 474
TR|S9XPZ9|S9XPZ9_CAMFR         ---------QIQQFENQYRGRELPGFVNYKTFETIIKKQVEILEEPAVDMLHRVTGTSRK 466
TR|A0A384DMX3|A0A384DMX3_URSMA YKAIYK---QIEKFENRYRGRELPGFVNYKTFEIIIKQQIKELEEPAVDMLHTVTDMVQA 480
TR|A0A3Q7W4X3|A0A3Q7W4X3_URSAR YKAIYK---QIEKFENRYRGRELPGFVNYKTFEIIIKQQIKELEEPAVDMLHTVTDMVQA 480
TR|A0A7E6D288|A0A7E6D288_9CHIR YQKTVN---KIIEFENRHRGRELPGFVNYKTFENLVREQLKTLEEPALDMLHNVTKIVHV 482
TR|S7NSW7|S7NSW7_MYOBR         YDDIFR---QIRKFENQYRGRELPGFVNYKTFETIIKKQVKTLEEPAVEMLHKITDMVRL 502
TR|A0A1N7THY9|A0A1N7THY9_PIPPI YEDIVR---HIKKFENQYRGKELPGFVNYKTFEMIIKKQVKALEDPAVEMLHRVTDMVRL 478
TR|A0A1N7THZ0|A0A1N7THZ0_ROUAE ICAMQN---ELYKFENQYRGRELPGFVNYKTFESIIKQQIKTLEEPAVDMLHTVTEMIWQ 480
TR|A0A1N7THY7|A0A1N7THY7_HYPMN YSALES---EILKFENQYRGRELPGFVNYKTFESIVKQQMKTLEEPAVEMLHTVTEMVWR 480
TR|A0A1N7THZ1|A0A1N7THZ1_STULI YEKIVN---KIIEFENRHRGRELPGFVNYKTFENLVRQQLKTLEEPALDMLHTVTKIVHL 480
TR|B2X026|B2X026_CHICK         EEIVCS---KLPKYEDQYRGREFPDFISYWTFEDIIKEQITKLEEPAVAMLNKVIYMVEE 526
TR|A0A172PZ65|A0A172PZ65_ANSCY QKSIPS---KMWKYEDQYRGREFPGFINYRTFEDIIKEHILDLEEPAIVILNNVIRLVEE 525
TR|A0A0Q3PUG8|A0A0Q3PUG8_AMAAE ------------------------------------------------------KGLVEE 348
SP|Q4ADG6|MX1_PHOVI            YKAIYK---QIEKFENRYRGRELPGFVNYKTFEIIIKQQIKELEEPAVYMLHMVTDMVQA 480
SP|Q4ADG7|MX1_OTABY            YKAICK---QIERFENRYRGRELPGFVNYKTFEIIIKQQIKELEEPAVYMLHTITDMVQA 480
TR|A0A2Y9G772|A0A2Y9G772_NEOSC YRVIYK---QIEKFENRYRGRELPGFVNYKTFEIIIKQQLKELEEPAVYMLHMVTDMVQA 480
TR|A0A6J2C7I6|A0A6J2C7I6_ZALCA YKAICK---QIERFENQYRGRELPGFVNYKTFEIIIKQQIKELEEPAVYMLHTITDMVQA 480
TR|A0A2U3WDI8|A0A2U3WDI8_ODORO YKAIYK---QIEKFENRYRGRELPGFVNYKTFEIIIKQQIKELEEPAVYMLHTITDMVQA 480
TR|A0A1U7SNC8|A0A1U7SNC8_ALLSI KRSIQE---DVWRFDNQYRGRELPGFINYKSFEDIVKKYIRELEGPALEVLSKVIDIVGQ 553
TR|A0A7M4FFF4|A0A7M4FFF4_CROPO QKVIHH---EVWRFESQYRGRELPGFINYRTFEDIVKQQIKELEEPAIEILKTVVEIVRQ 461
TR|A0A340WTU5|A0A340WTU5_LIPVE RTLHQP------------------------------------------------------ 436
TR|A0A6J3RAX5|A0A6J3RAX5_TURTR GQHPHL---QPHPLDHPVLHAQ-----------------DVRPAAEGDAAAA--------- 457
TR|A0A341CI24|A0A341CI24_NEOAA ESKAPW---ETLRSAPAISSHT----------------PV---------ML--------- 528
TR|A0A2Y9LTL6|A0A2Y9LTL6_DELLE D-TRHL---QPHPLD-----AQ----------------DVRPAAEGH---A--------- 452
TR|A0A383ZCI9|A0A383ZCI9_BALAS YNAICK---EIQQCENRYRGRELPGFVNYKTFEAIIKKQVGVLEEPAVDMLHTVTDIIRA 481
TR|V9KIB1|V9KIB1_CALMI         QCDLKN---NVCAYERMYRGRELPGFTNYKTFEAIIKQQVCRLEQPAMLKLKNITEIVHE 447
TR|A0A674K7E1|A0A674K7E1_TERCA KEIMKE---EVWGFENHYRGRELPGFVNYKTFEAIVKQQIVLLKSPAIMILKTVTELVRE 489
TR|A0A4D9DW37|A0A4D9DW37_9SAUR RKTIQH---EVRRFEEQYRGRELPGFTSYKTFEAIVRQRIMALEKPAIEMLKKVIEIVRK 595
TR|H2DQ54|H2DQ54_XENLA         QDKSRD---DISEFENKYRGRELPGFISFKVFENIARGQIRCLEEPAMEKLKEVTEIVKA 446
TR|A0A0F6MUS9|A0A0F6MUS9_ANGJA NQRIED---EVQEYEVKYRGRELPGFINYKTFEVLVKDQLKMLEEPAVKKLKGVTDIVRK 461
TR|Q98990|Q98990_SALSA         NQRIEG---EVADYEKTYRGRELPGFINYKTFEVMVKDQIKQLEEPAVKKLKEISDAVRK 446
TR|A0A674EGU5|A0A674EGU5_SALTR NQRIEG---EVADYEMTYRGRELPGFINYKTFEVMVKDQIKQLEEPAVKKLKEISDAVRK 453
SP|Q91192|MX1_ONCMY            NQRIEG---EVDDYEKTYRGRELPGFINYKTFEVMVKDQIKQLEGPAVKKLKEISDAVRK 446
TR|A0A286K200|A0A286K200_MYLPI NKKIER---EVDNYEAKYRGRELPGFINYKTFEGLVRDQIKMLEEPALKTLKTIADVVRR 449
TR|A0A498MHV8|A0A498MHV8_LABRO KRKIEK---EIESYEAKYRGRELPGFINYKTFEGLVREQIKLLEDPALKTLKTISDMVRR 450
SP|Q7T2P0|MX1_ICTPU            KSRIEK---EVNEYEEKYRGRELPGFINYKTFEVIVKDQIKQLEEPAIRRLKEISDLIRK 451
TR|A0A556TK66|A0A556TK66_BAGYA NERIDK---EVSEYEEKYRGRELPGFINYKTFEVIVKDQIRQLEEPAIRRLKEISDLIRK 419
TR|A0A3G6IID3|A0A3G6IID3_9TELE NKKIEK---EVDNYEAKYRGRELPGFINYKTFEGLVREQIKMLEEPALKTLKTVSDVVRR 449
SP|Q8JH68|MXA_DANRE            NKKIEK---EVDNYEVKYRGRELPGFINYKTFEGLVRDQIKLLEEPALKTLKTVSDVVRK 449
TR|A0A672MEY2|A0A672MEY2_SINGR KARVTE---MIEKYNEIHRGRELLTFSEFCEYECVIQNHVAALQEPAMATLKDVRGTVSF 446
TR|Q7T2M3|Q7T2M3_CARAU         NKKIEK---EVDNYEAKYRGRELPGFSNYKTLEGLVREQIKLLEEPALKTLKTVSDVVRK 449
TR|A0A673LWX8|A0A673LWX8_9TELE NRRIEK---EVDNYETKYRGRELPGFINYKTFEGLVQEQIKLLEKPALKTLKTVSDVVRR 449
TR|A0A8B9KF29|A0A8B9KF29_ASTMX NRRIEK---EVQEYEVKYRGRELPGFINYKTFEVMVKDQIKKLEEPAVRKLKEISGR--- 443


SP|P20591|MX1_HUMAN            AFTDVSI--------KNFEEFFNLHRTAKSKIEDIRAEQER--EGEKLIRLHFQMEQIVY 532
TR|K4JBQ0|K4JBQ0_9PRIM         AFTDVSI--------KNFEEFFNLHRTAKSKIEDIRAEQER--EGEKLIRLHFQMEQIVY 532
TR|H2QL18|H2QL18_PANTR         AFTDVSI--------KNFEEFFNLHRTTKSKIEDIRAEQER--EGEKLIRLHFQMEQIVY 532
SP|A1E2I4|MX1_MACMU            AFTDVSM--------KNFEELFNLHRTAKSKIEDIRTEQER--EGEKLIRLHFQMEQIVY 532
SP|Q5R5G3|MX1_PONAB            AFTDVSI--------KNFEEFFNLHRTAKSKIEDIRAEQER--EGEKLIRLHFQMEQIVY 532
TR|A0A6P8QPW8|A0A6P8QPW8_GEOSA TFLKYSR--------KHFAHFHNLDKAAKVIIEDLRKIQEA--EAENMINNQFKMEQHIY 497
TR|U3EGN4|U3EGN4_CALJA         AFTDVSI--------KNFDEFFNLHRTAKSKIEDIRAEQER--EGEKQIRLQFQMEQIVY 532
SP|Q9N0Y3|MX1_CANLF            AFGDISK--------ANFDEFFNLYRTTKSKIEDIKFELEK--EAEKSIRLHFQMEQIVY 528
TR|M3W8X0|M3W8X0_FELCA         AFANISK--------GNFDEFFNLYRTTKSKIEDIKFELEN--EAEKSIRLHFQMEQIVY 530
TR|A0A667GNL8|A0A667GNL8_LYNCA AFANISK--------GNFDEFFNLYRTTKSKIEDIKFELEN--EAEKSIRLHFQMEQIVY 530
TR|A0A6P6GZN4|A0A6P6GZN4_PUMCO AFANISK--------GNFDEFFNLYRTTKSKIEDIKFELEN--EAEKSIRLHFQMEQIVY 530
TR|A0A6J1XXY6|A0A6J1XXY6_ACIJB AFANISK--------GNFDEFFNLYRTTKSKIEDIKFELEN--EAEKSIRLHFQMEQIVY 530
TR|A0A6P4TS13|A0A6P4TS13_PANPR AFANISK--------GNFDEFFNLYRTTKSKIEDIKFELEN--EAEKSIRLHFQMEQIVY 568
TR|A0A3Q7SCG7|A0A3Q7SCG7_VULVU AFGDISK--------ANFDEFFNLYRTTKSKIEDIKFELEK--EAEKSIRLHFQMEQIVY 566
TR|I3MJX6|I3MJX6_ICTTR         AFTNASE--------KNFAEFFNLHKTAKSKIEDIRLEQEQ--GAERCIRLHFQMEQIVY 533
TR|G9KBY2|G9KBY2_MUSPF         AFTGISK--------ANFDEFFNLYRTTKSKIEDIKFELEE--EAEKSIRLHFQMEQIVY 527
TR|A0A2Y9KR88|A0A2Y9KR88_ENHLU AFTGISK--------SNFDEFFNLYRTTKSKIEDIKFELEE--EAEKSIRLHFQMEQIVY 527
SP|P09922|MX1B_MOUSE           AFVKILS--------NDFGDFLNLCCTAKSKIKEIRLNQEK--EAENLIRLHFQMEQIVY 498
SP|P18588|MX1_RAT              AFTKVSS--------NNFGDFLNLHSTAKSKIEDIRFNQEK--EAEKLIRLHFQMEHIVY 525
TR|A0A6P6D2J7|A0A6P6D2J7_PTEVA AFTEVSQ--------KNYNEFFNLYKVSKSKIEDIKTEQER--EAEKSIRLHFQMEQIVY 567
TR|A0A6J0DKA3|A0A6J0DKA3_PERMB AFTKVSL--------NNFGEFFNLCSTVKSKIEDIRLNQEKKKKTEKLIQLHIQMEQIVY 491
TR|G5C5C9|G5C5C9_HETGA         TFTAVSE--------RNFSGFFNLHKTCKVSPGCAEKAAEIAREQASLL--PRAPLRCPL 532
TR|A0A3Q0CWV5|A0A3Q0CWV5_MESAU AFTTVSF--------NNFGEFFNLHSAAKSKMEDIRLSQEK--EAEKLIRLHFQMEQIVY 517
TR|A0A6I9K5X3|A0A6I9K5X3_CHRAS AFTDVSK--------KHFDEFFNLHRTTKTKIEDIKLEQEK--EAEKMIQLHFQMEQIVY 533
SP|P79135|MX1_BOVIN            TFTDVSG--------KHFNEFFNLHRTAKSKIEDIRLEQEN--EAEKSIRLHFQMEQLVY 521
TR|A0A6P3H2K4|A0A6P3H2K4_BISBI TFTDVSG--------KHFNEFFNLHRTAKSKIEDIRLEQEN--EAEKSIRLHFQMEQLVY 527
TR|L8IPR7|L8IPR7_9CETA         TFTDVSG--------KHFNEFFNLHRTAKSKIEDIRLEQEN--EAEKSIRLHFQMEQLVY 524
SP|P27594|MX1_PIG              AFTDVSE--------TNFNEFFNLHRTAKSKIEDIKLEQEK--EAETSIRLHFQMEQIVY 533
TR|A0A1U7U9K3|A0A1U7U9K3_CARSF AFTDVSV--------KHFEEFFNLHRTAKSKIEDIRAEQEK--EGDKLIRLHFQMEQIVY 497
SP|P33237|MX1_SHEEP            TFTEVSG--------KHFSEFFNLHRTAKSKIEDIRLEQEN--EAEKSIRLHFQMEQLVY 527
```

```
TR|A0A7D4WPX4|A0A7D4WPX4_CAPHI TFTEVSG--------KHFSEFFNLHRTAKSKIEDIRLEQEN--EAEKSIRLHFQMEQLVY 527
SP|Q28379|MX1_HORSE            TFTKVSE--------KNFSEFFNLHRTTKSKLEDIKLEQEN--EAEKSIRLHFQMEKIVY 532
TR|A0A6I9I6Y4|A0A6I9I6Y4_VICPA AFTNVSE--------THFNEFFNLHRTAKSKIEDIKLDLEN--EAEKVIRLHFQMEQIVY 535
TR|A0A6P5BYP3|A0A6P5BYP3_BOSIN TFTDVSG--------KHFNEFFNLHRTAKSKIEDIRLEQEN--EAEKSIRLHFQMEQLVY 524
TR|S9XPZ9|S9XPZ9_CAMFR         LCPPPPAETQQPTALTAPSPYVLCPLVSKSKVEDIKLDLEN--EAEKLIRLHFQMEQIVY 524
TR|A0A384DMX3|A0A384DMX3_URSMA AFTDISK--------ANFDEFFNLYRTTKSKIEDIKFELEK--EAEKSIRLHFQMEQIVY 530
TR|A0A3Q7W4X3|A0A3Q7W4X3_URSAR AFTDISK--------ANFDEFFNLYRTTKSKIEDIKFELEK--EAEKSIRLHFQMEQIVY 530
TR|A0A7E6D288|A0A7E6D288_9CHIR AFSDVTK--------KNYEEFFNFLKVCKANIEDIKVKQEE--EAEKCIRLQFQMEQIIY 532
TR|S7NSW7|S7NSW7_MYOBR         AFTDVSK--------KNYEEFFNLFRTCKSKIEDIKSEQEK--EAEKSIRLHFQMEQIVY 552
TR|A0A1N7THY9|A0A1N7THY9_PIPPI AFADACK--------KNYEEFFNLLRTCVSKIEDIKSEQEK--EAEKSIRLHFQMEQIVY 528
TR|A0A1N7THZ0|A0A1N7THZ0_ROUAE AFTDVSQ--------KNFSEFFNLYRVSKSKIEDIKTEQER--EAEKSIRLHFQMEQMVY 530
TR|A0A1N7THY7|A0A1N7THY7_HYPMN AFADVSQ--------KHFNEFFNLYRVSKSKIEDLKTEKER--EAEKFIRIHFEMEQMVY 530
TR|A0A1N7THZ1|A0A1N7THZ1_STULI AFSNVTK--------KNYEEFFNLLKVCKASVEDIKVKEEE--EAEKCIRLQFQMEQIVY 530
TR|B2X026|B2X026_CHICK         KFLQLAN--------KRFANFQNLNNAAQARIGCISDRQAT--TAKNCILTQFKMERIIY 576
TR|A0A172PZ65|A0A172PZ65_ANSCY KFLELTN--------KHFANFQNLNRAAQTRIDVIRDIQAM--NAENHIRNQFKMERIVY 575
TR|A0A0Q3PUG8|A0A0Q3PUG8_AMAAE KFTELTI--------RHFGNFHNLNRAVKTRIEDVREKQAE--EAQRHIRTQFKMERIVY 398
SP|Q4ADG6|MX1_PHOVI            AFTDISE--------ANFAEFFNLYRTTKSKIEDIKFELEK--EAEKSIRLHFQMEQIVY 530
SP|Q4ADG7|MX1_OTABY            AFTDISE--------ANFAEFFNLYRTTKSKIEDIKFELEK--EAEKSIRLHFQMEQIVY 530
TR|A0A2Y9G772|A0A2Y9G772_NEOSC AFTDISK--------ANFAEFFNLYRTTKSKIEDIKFELEE--EAEKSIRLHFQMEQIVY 530
TR|A0A6J2C7I6|A0A6J2C7I6_ZALCA AFTDISE--------ANFAEFFNLYRTTKSKIEDIKFELEK--EAEKSIRLHFQMEQIVY 530
TR|A0A2U3WDI8|A0A2U3WDI8_ODORO AFTDISE--------ANFAEFFNLYRTTKSKSEDIKFELEK--EAEKSIRLHFQMEQIVY 530
TR|A0A1U7SNC8|A0A1U7SNC8_ALLSI SFTEVAT--------ANFPEFFNLTRAAKGRIEDIKMQQRA--EAEKMIQTQFKIEQIVY 603
TR|A0A7M4FFF4|A0A7M4FFF4_CROPO HFTVLAK--------AHFDDFYNLNRAAKDRIEDIREKQAE--AAETMIRTQFKMEQMVY 511
TR|A0A340WTU5|A0A340WTU5_LIPVE -------------------------------------------------------CSLG- 440
TR|A0A6J3RAX5|A0A6J3RAX5_TURTR -----------------------------------------------------------A 458
TR|A0A341CI24|A0A341CI24_NEOAA -----------------------------------------------------------K 529
TR|A0A2Y9LTL6|A0A2Y9LTL6_DELLE -----------------------------------------------------------A 453
TR|A0A383ZCI9|A0A383ZCI9_BALAS AFTDVSE--------KNFNEFFNLHRTAKSKIEDIRLEQEK--EAEKSIRLHFQMEQIVY 531
TR|V9KIB1|V9KIB1_CALMI         SFMAIAL--------HNFNGVPNLLKASKMKMEELKKKQER--HAVSQLRIQFKLEDIVY 497
TR|A0A674K7E1|A0A674K7E1_TERCA TFTEIAM--------QHFENFFYLFREAKTRIEVIKQKQEE--EAETVIANQFKMERIVY 539
TR|A0A4D9DW37|A0A4D9DW37_9SAUR SFTE-------------------------DRIEGISKKQSE--EAETILRIHFKMEQLVY 628
TR|H2DQ54|H2DQ54_XENLA         HFNQIAM--------KHFLAFPNLYRGAKVRIEDICHKQVW--EAERTIRTQFKMEKIIY 496
TR|A0A0F6MUS9|A0A0F6MUS9_ANGJA AFLQLAH--------NNLMGYPNLLKSAKTKIEDIKKEKES--SAESMLRTQFRMELIVY 511
TR|Q98990|Q98990_SALSA         VFLLLAQ--------SSFIGFPNLLKSAKTKIEAIKQVNES--TAESMLRTQFKMEMIVY 496
TR|A0A674EGU5|A0A674EGU5_SALTR VFLLLAQ--------SSFIGFPNLLKSAKTKIEAIKQVNES--TAESMLRTQFKMELIVY 503
SP|Q91192|MX1_ONCMY            VFLLLAQ--------SSFTGFPNLLKSAKTKIEAIKQVNES--TAESMLRTQFKMELIVY 496
TR|A0A286K200|A0A286K200_MYLPI KFIQLAQ--------YSFAGFPNLLKIAKTKIEAIKLDKES--LAESMLRTQFKMELIVY 499
TR|A0A498MHV8|A0A498MHV8_LABRO KFVQLAQ--------CSFVGFPNLLKIAKTKIEAIKQDKES--LAESMLRTQFKMELIVY 500
SP|Q7T2P0|MX1_ICTPU            GFIQLAQ--------NSFLGFPNLLKMAKTKIECIKQVKES--EAETMLRTQFKMELIIY 501
TR|A0A556TK66|A0A556TK66_BAGYA NFIQLAQ--------HSFVGFPNLLKMAKSKIECIKQVQES--EAETMVRTQFKMELIVY 469
TR|A0A3G6IID3|A0A3G6IID3_9TELE KFIQLAQ--------YSFAGFPNLLKIAKTKIEAIKLDKET--LAESMLRTQFKMELIVY 499
SP|Q8JH68|MXA_DANRE            KFIQLAQ--------CSFIGFPNLLKIAKTKIEGIKLNKES--LAESMLKTQFKMELIVY 499
TR|A0A672MEY2|A0A672MEY2_SINGR IFIVFII-------IIIIVIFHVLPCIPQNMVNDIQTKQEA--KAEKRIKEFINMEKLVF 497
TR|Q7T2M3|Q7T2M3_CARAU         KFIQLAQ--------YSFIGFPNLLKIAKTKIEAIKQDKES--QAESMLRTQFKMELIVY 499
TR|A0A673LWX8|A0A673LWX8_9TELE KSIQLAQ--------CSFIGFPNLLKIAKTKIEAIKQDKEC--LAESMLRTQFKMELIVY 499
TR|A0A8B9KF29|A0A8B9KF29_ASTMX -QQSLAQ--------NSFVGFPNLLRTAKTRIDSIKQEKEA--EAESMLRTQFKMELIIY 492


SP|P20591|MX1_HUMAN            CQDQVYRGALQKVR---EKE--LEEEKKKKSWD-FGA--------------------FQS 566
TR|K4JBQ0|K4JBQ0_9PRIM         CQDHVYRGALQKVR---EKE--LEEEKKKKSWD-FGA--------------------FQS 566
TR|H2QL18|H2QL18_PANTR         CQDQVYRGALQKVR---EKE--LEEEKKKKSWD-FGA--------------------FQS 566
SP|A1E2I4|MX1_MACMU            CQDQVYRGALQKVR---EKE--LEEEKKKKSWD-IGT--------------------FQP 566
SP|Q5R5G3|MX1_PONAB            CQDQVYRGALQKVR---EKE--LEEEKKKKSWD-FGA--------------------FQS 566
TR|A0A6P8QPW8|A0A6P8QPW8_GEOSA AQDKIYWRDFLTLQKQNAIT-----SNKTV-----------------------NELTLGV 529
TR|U3EGN4|U3EGN4_CALJA         CQDQVYRSALQKVR---EKE--LEEEKRSKKSC-FAM--------------------VVE 566
SP|Q9N0Y3|MX1_CANLF            CQDHVYQRALQRVR---EKD--SDEEKK-KKTS-S-M-----------------S--HDE 561
TR|M3W8X0|M3W8X0_FELCA         CQDQIYQRALQRVR---ERV--SEEEKRNKKIN-SPI--------------------SEE 564
TR|A0A667GNL8|A0A667GNL8_LYNCA CQDQIYQRALQRVR---ERV--SEEEKRNKKTN-SPI--------------------SEE 564
TR|A0A6P6GZN4|A0A6P6GZN4_PUMCO CQDQIYQRALQRVR---ERV--SEEEKRNKKTN-SPI--------------------SEE 564
TR|A0A6J1XXY6|A0A6J1XXY6_ACIJB CQDQIYQRALQRVR---ERV--SEEEKRNKKTN-SPI--------------------SEE 564
TR|A0A6P4TS13|A0A6P4TS13_PANPR CQDQIYQRALQRVR---ERV--SEDEKRNKKIN-SPI--------------------SEE 602
TR|A0A3Q7SCG7|A0A3Q7SCG7_VULVU CQDQVYQRALQRVR---EKD--SDEEKKNKKTS-S-M-----------------S--HDE 600
TR|I3MJX6|I3MJX6_ICTTR         CQDQVYRGALQGVR---EKE--AEEEKRKSHG----Q--------------------SQP 564
TR|G9KBY2|G9KBY2_MUSPF         CQDQVYQRALQRVR---EKE--SEEGKKNQKIN-SMM-----------------T--TEE 562
TR|A0A2Y9KR88|A0A2Y9KR88_ENHLU CQDQGYQRALQKIR---EKV--LEEGKKNQKIN-SVM-----------------T--TEK 562
SP|P09922|MX1B_MOUSE           CQDQVYKETLKTIR---EKE--AEKEKTKALIN-PAT-----------------FQNNSQ 535
SP|P18588|MX1_RAT              CQDQAYKKALQEIR---EKE--AEKEKSTF-----GA-----------------FQ--HN 556
TR|A0A6P6D2J7|A0A6P6D2J7_PTEVA CQDTVYQTSLGALR---QPA--RSESLLG------------------------------- 591
TR|A0A6J0DKA3|A0A6J0DKA3_PERMB CQEQVYRRALKEVK---EKE--AEEEENNFNF-----------------------QCNSQ 523
TR|G5C5C9|G5C5C9_HETGA         CVGEVGARRPGNSNQQCGVRAGVRDAKGNSRSRARAADSGPLGGKGPTWMLSKGPQGLGR 592
TR|A0A3Q0CWV5|A0A3Q0CWV5_MESAU CQDQVYKGALKKVR---EED--AEREKNKV------T-----------------LQFNSA 549
TR|A0A6I9K5X3|A0A6I9K5X3_CHRAS CQDKIYSGALRDIR---EKE--MEEEKRNSTQH-YSQ--------------------SQK 567
SP|P79135|MX1_BOVIN            CQDQVYRRALQQVR---EKE--AEEEKNKKSNH-YFQ----------------------- 552
TR|A0A6P3H2K4|A0A6P3H2K4_BISBI CQDQVYRRALQQVR---EKE--AEEEKNKKSNH-YFQ----------------------- 558
TR|L8IPR7|L8IPR7_9CETA         CQDQVYRRALQQVR---EKE--AEEEKNKKSNH-YFQ----------------------- 555
SP|P27594|MX1_PIG              CQDQVYRGALQKVR---EKE--AEEEKNRKSNQ-YFL---SSP----------------- 567
TR|A0A1U7U9K3|A0A1U7U9K3_CARSF CQDQVYRGALQKVR---EKE--TEEEKRNKRSH-DS---------------------FFE 530
SP|P33237|MX1_SHEEP            CQDQVYRRALQQVR---EKE--AEEEKKKKSNH-YYQ----------------------- 558
TR|A0A7D4WPX4|A0A7D4WPX4_CAPHI CQDQVYRRALQQVR---EKE--AEEEKKKQSNH-YYQ----------------------- 558
SP|Q28379|MX1_HORSE            CQDHVYRGTLQKVR---ENE--MEEEKKKKTIN---V--------------------WGQ 564
TR|A0A6I9I6Y4|A0A6I9I6Y4_VICPA CQDKVYRGALQKAR---EKE--AEEEKNKKSKH--SF----------------------S 566
TR|A0A6P5BYP3|A0A6P5BYP3_BOSIN CQDQVYRRALQQVR---EKE--AEEEKNKKSNH-YFQ----------------------- 555
TR|S9XPZ9|S9XPZ9_CAMFR         CQDKVYRGALQKAR---EKE--AEEERKKKLQP-WSF--QPSPSVDPSIA--EIFQHLTA 574
TR|A0A384DMX3|A0A384DMX3_URSMA CQDQVYQRALQRVR---EKE--SDEEKKSKKKD-S-M-----------------I--AEE 564
TR|A0A3Q7W4X3|A0A3Q7W4X3_URSAR CQDQVYQRALQRVR---EKE--SDEEKKSKKKD-S-M-----------------I--AEE 564
TR|A0A7E6D288|A0A7E6D288_9CHIR CQDHEYQTWLRKIR---ETE--LQQEKTKSSSM------------------------PSE 563
TR|S7NSW7|S7NSW7_MYOBR         CQDQAYRGALQKIR---EKE--SKEQKGSSREQ---T--------------------SSL 584
TR|A0A1N7THY9|A0A1N7THY9_PIPPI CQDQAYRTALGKIR---GME--AEENKKKKKEH---I--------------------FFE 560
```

```
TR|A0A1N7THZ0|A0A1N7THZ0_ROUAE CQDQVYRKSLQIVR---EKE--KEKVEQNKNKS-RVS-----------------D--LEQ 565
TR|A0A1N7THY7|A0A1N7THY7_HYPMN CQDQVYQRSLQKVR---EKE--N--EEQNKNKS-RVL-----------------D--LVQ 563
TR|A0A1N7THZ1|A0A1N7THZ1_STULI CQDQEYRTWLQKIR---EKE--LEQQKKKLAFA------------------------PSE 561
TR|B2X026|B2X026_CHICK         CQDNIYTDDLKAARAEGISK-----DTKIK-----------------------DLAFGCA 608
TR|A0A172PZ65|A0A172PZ65_ANSCY CQDNIYLDDLNSTKAEILSK-----DS-GK-----------------------EISFGSV 606
TR|A0A0Q3PUG8|A0A0Q3PUG8_AMAAE CQDNLYKNDLESVNREYLNK-----VSNGK-----------------------ESQVGSG 430
SP|Q4ADG6|MX1_PHOVI            CQDQVYQRALQRVR---EKV--ADEE-KNKKIN-S-M-----------------S--SEE 563
SP|Q4ADG7|MX1_OTABY            CQDQVYQCALQRVR---EES--D--KEKDKKIN-S-M-----------------C--SKE 562
TR|A0A2Y9G772|A0A2Y9G772_NEOSC CQDQVYQRALQRVR---EKV--SEEEKKNKKIN-S-M-----------------S--SEE 564
TR|A0A6J2C7I6|A0A6J2C7I6_ZALCA CQDQVYQCALQRVR---EEW--D--KGKDRKIN-S-M-----------------C--SKE 562
TR|A0A2U3WDI8|A0A2U3WDI8_ODORO CQDQVYQCALQRVR---EES--D--KEKNKKVN-S-M-----------------C--SEE 562
TR|A0A1U7SNC8|A0A1U7SNC8_ALLSI CQDNFYKKALRATNVNNVEE-----ALKTL---KFQD------------------VNGIV 637
TR|A0A7M4FFF4|A0A7M4FFF4_CROPO CQDNIYSEDLKSIREETAEQ-----ATDKK---TMTP-------------KSSKFNFGTT 550
TR|A0A340WTU5|A0A340WTU5_LIPVE ------------------------------------------------------------
TR|A0A6J3RAX5|A0A6J3RAX5_TURTR EQGQVRQAAEGAQRHQRQEE--APEGAAAAADP-GSA--RQVPRLNPPSAPPRGLQGAGD 513
TR|A0A341CI24|A0A341CI24_NEOAA TFGQLRKGMLQLLKERRDT----SDRRKPLREP-LQR--LTLARLAK-------FPG--- 572
TR|A0A2Y9LTL6|A0A2Y9LTL6_DELLE EQGQVRQAAEGAQRHQRQEE--APEGAAAAADP-GSA--CQVPRLNPPSAPPRGLQGAGD 508
TR|A0A383ZCI9|A0A383ZCI9_BALAS CQDQVYRRALQKVR---EKE--AAEEKSKKPNS-HFQ-----P----------------- 563
TR|V9KIB1|V9KIB1_CALMI         SQDSNYSDNLELKATEEYRE-----QTPDQ--N-------------------SLTSKMST 531
TR|A0A674K7E1|A0A674K7E1_TERCA CQDSLYHEDLQEIRKTTGVL-----PLGVA---V--------------------THVKSI 571
TR|A0A4D9DW37|A0A4D9DW37_9SAUR CQDNVYRQDLKVIREETPKE-----AANSL---N--P-------------VTLSFTFG-- 663
TR|H2DQ54|H2DQ54_XENLA         CQDRLYSGRLNEVRSADAKP-----TNTTF---L--------------------SIQTPA 528
TR|A0A0F6MUS9|A0A0F6MUS9_ANGJA TQDSTYSHNLDEIRREDEED-----EE-DRTLTAVKQ-------------HKF-KRSIVF 551
TR|Q98990|Q98990_SALSA         TQDSTYSHSLSERKREEEDD-----RP-LPTI--------------------K-IRSTIF 529
TR|A0A674EGU5|A0A674EGU5_SALTR TQDSTYSHSLSERKREEEDD-----QP-LPTI--------------------K-IRSTIF 536
SP|Q91192|MX1_ONCMY            TQDSTYSHSLCERKREEDED-----QP-LT----------------------E-IRSTIF 527
TR|A0A286K200|A0A286K200_MYLPI SQDGTYSQSLQHAKNKLEED-----ENNEDA-----K-------------KSVNCMSVGI 536
TR|A0A498MHV8|A0A498MHV8_LABRO SQDGTYSQSLKDAKDKLEEE-----EERK---------------------PKFHFGSVDN 534
SP|Q7T2P0|MX1_ICTPU            TQDSMYSDTLSTLKVKEEEG-----ER-QKVGILPNS-------------YSIS-CSLYN 541
TR|A0A556TK66|A0A556TK66_BAGYA TQDSMYSSTLSELKLEKERE-----EE-ELRLNGFRP-------------SSTS-CSLYD 509
TR|A0A3G6IID3|A0A3G6IID3_9TELE SQDGTYSQSLQHAKNKLEED-----ESDEDT-----K-------------KSVNCLSIGI 536
SP|Q8JH68|MXA_DANRE            SQDGTYSQSLKHAKDKLEEM-----EKERPQ---PKI-------------KLPLLSSFDL 538
TR|A0A672MEY2|A0A672MEY2_SINGR TQDKVLQQKLNDSDIPQKTG--IETSDKNH----YDDD---------------DTIFNSK 536
TR|Q7T2M3|Q7T2M3_CARAU         SQDGTYSQSLQHAKDKLEEI-----ENDKQQLPQFNA-------------KKLNFVSVDV 541
TR|A0A673LWX8|A0A673LWX8_9TELE SQDGTYSQSLKHAKDKLEKA-----ENEKL---AFKP-------------KRSSIVSVDV 538
TR|A0A8B9KF29|A0A8B9KF29_ASTMX TQDSTYSNTLISLKREEEEG-----EE-QKGGNSYVV-------------SGFGFSSLLS 533


SP|P20591|MX1_HUMAN            SSATDSSMEEIFQHLMAYHQEASKRISSHIPLIIQFFMLQTYGQQ--LQKAMLQLLQDKD 624
TR|K4JBQ0|K4JBQ0_9PRIM         SSATDSSMEEIFQHLMAYHQEASKRISSHIPLIIQFFMLQTYGQQ--LQKAMLQLLQDKD 624
TR|H2QL18|H2QL18_PANTR         SSATDPSMEEIFQHLMAYHQEASKRISSHIPLIIQFFMLQTYGQQ--LQKAMLQLLQDKD 624
SP|A1E2I4|MX1_MACMU            SS-TESSMEEIFQHLMAYHQEASKRISSHIPLVIQFFMLQMYGQQ--LQKAMLQLLQDKD 623
SP|Q5R5G3|MX1_PONAB            SSATDSSMEEIFQHLMAYHQEASKRISSHIPLIIQFFMLQTYGQQ--LQKAMLQLLQDKD 624
TR|A0A6P8QPW8|A0A6P8QPW8_GEOSA VTPTLPQFTEITRHVKAYCSGAFLRLSSQIPMIIYYYALHMYGDM--VERKMLQILQDAN 587
TR|U3EGN4|U3EGN4_CALJA         PSSSESSMEEIFQHLMAYHQECSKRISSHIPLIIQFFVLQTYGQQ--LQKAMLQLLQDKD 624
SP|Q9N0Y3|MX1_CANLF            VSSVNISLSEILEHLLAYRQEATNRISSHIPLIIQYFILQVYGQK--LQNGMLQLLQDKD 619
TR|M3W8X0|M3W8X0_FELCA         VSSGSVSLCEIFEHLLAYRQEAANRISSHIPLIIQYFILQVYGQR--LQKNMLQLLQDKE 622
TR|A0A667GNL8|A0A667GNL8_LYNCA VSSGSVSLCEIFEHLLAYRQEAANRISSHIPLIIQYFILQVYGQR--LQKNMLQLLQDKE 622
TR|A0A6P6GZN4|A0A6P6GZN4_PUMCO VSSGSVSLCEIFEHLLAYRQSGISSPGSREYSGS-------------------------- 598
TR|A0A6J1XXY6|A0A6J1XXY6_ACIJB VSSGSVSLCEIFEHLLAYRQEAANRISSHIPLIIQYFILQVYGQR--LQKNMLQLLQDKE 622
TR|A0A6P4TS13|A0A6P4TS13_PANPR VSSGNVSLCEILEHLLAYRQEAANRIASHIPLIIQYFILQVYGQR--LQKSMLQLLQDKE 660
TR|A0A3Q7SCG7|A0A3Q7SCG7_VULVU VSSVNISLSEILEHLLAYRQEATNRISSHIPLIIQYFILQVYGQK--LQNGMLQLLQDKD 658
TR|I3MJX6|I3MJX6_ICTTR         VSAEESSMAEIFQHLDAYRKEAHNRISSLVPLIIQFFVLKTFGQK--LQTAMLLLLQDKD 622
TR|G9KBY2|G9KBY2_MUSPF         FSLVNISLSEIFEHLMAYHQEA-TRISSHIPLIIQYFILQVYGQK--LQKAMLQLLQDKD 619
TR|A0A2Y9KR88|A0A2Y9KR88_ENHLU LSPVNISLSEISEHVVAYCQEATTRISSHIPLIIQYFVLQVYEQK--LQKAMLQLLQDKD 620
SP|P09922|MX1B_MOUSE           FPQKGLTTTEMTQHLKAYYQECRRNIGRQIPLIIQYFILKTFGEE--IEKMMLQLLQDTS 593
SP|P18588|MX1_RAT              SPRKELTTTEMTQHLNAYYQECGRNIGRQIPLIIQYSILQTFGQE--MEKAMLQLLQDTS 614
TR|A0A6P6D2J7|A0A6P6D2J7_PTEVA ----VSTSQCPNPFVNCSFQETGNRLSTHIPLMIQFFVLQSFGQQ--LQKAMLQLLQDKE 645
TR|A0A6J0DKA3|A0A6J0DKA3_PERMB APQEDLSTTEMSQHPNAYYEECRRSIGRQVSLIIQYFLLPTFGED--MEKAMLQLLQDKT 581
TR|G5C5C9|G5C5C9_HETGA         GRR----------NKTAVAQVCALRVPAGSTGLSAWQGPSSAGLQPQPEGGHLAGLGPER 642
TR|A0A3Q0CWV5|A0A3Q0CWV5_MESAU SSQKDLSITEMTQHLNAYYEECRRSIGRQVPLIIQYFILQRFGEE--MEKAMLQLLQDKV 607
TR|A0A6I9K5X3|A0A6I9K5X3_CHRAS VSSLDSFKEELSQHLFAYHKEVSNRMSNHIPLVIQFFILQLYSQQ--LQKGMLQLLQNKD 625
SP|P79135|MX1_BOVIN            SQVSEPSTDEIFQHLTAYQQEVSTRISGHIPLIIQFFVLRTYGEQ--LKKSMLQLLQDKD 610
TR|A0A6P3H2K4|A0A6P3H2K4_BISBI SQVSEPSTDEIFQHLTAYQQEVSTRISGHIPLIIQFFVLRTYGEQ--LKKSMLQLLQDKD 616
TR|L8IPR7|L8IPR7_9CETA         SQVSEPSTDEIFQHLTAYQQEVSTRISGHIPLIIQFFVLRTYGEQ--LKKSMLQLLQDKD 613
SP|P27594|MX1_PIG              APSSDPSIAEIFQHLIAYHQEVGKRISSHIPLIIQFFILRTFGQQ--LQKSMLQLLQNKD 625
TR|A0A1U7U9K3|A0A1U7U9K3_CARSF QDSENSSMAEIFQHLVAYHKEVSKRLSSHIPLTIQFFILQTFGEQ--LQKGMMQLLQEKD 588
SP|P33237|MX1_SHEEP            SEDSEPSTAEIFQHLMAYHQEVSTRISSHIPLIIQFFVLRTYGEQ--LKKSMLQLLQDKD 616
TR|A0A7D4WPX4|A0A7D4WPX4_CAPHI SEDSEPSTAEIFQHLMAYHQEVSTRISSHIPLIIQFFVLRTYGEQ--LKKSMLQLLQDKD 616
SP|Q28379|MX1_HORSE            NTSTESSMAEILEHLNAYQHEAGNRLSTHIPLIIQFFVLQTFGQQ--LQKSMLQLLQDRD 622
TR|A0A6I9I6Y4|A0A6I9I6Y4_VICPA TLTEDPSIAEVVQHLTAYHEEVSNRISSHIPLIIQYFVLRSFGQQ--LQKGMLQLLQNKD 624
TR|A0A6P5BYP3|A0A6P5BYP3_BOSIN SQVSEPSTDEIFQHLTAYQQEVSTRISGHIPLIIQFFVLRTYGEQ--LKKSMLQLLQDKD 613
TR|S9XPZ9|S9XPZ9_CAMFR         YHEVSPSIAEIFQHLTAYHEEVSNRISSHIPLIIQYFVLRSFGQQ--LQKGMLQLLQNKD 632
TR|A0A384DMX3|A0A384DMX3_URSMA ISSVNISLSEIFEHLLAYRQEANNRISSHIPLIIQYFILQVFGQK--LQKGMLQLLQDKD 622
TR|A0A3Q7W4X3|A0A3Q7W4X3_URSAR ISSVNISLSEIFEHLLAYRQEATNRISSHIPLIIQYFILQVFGQK--LQKGMLQLLQDKD 622
TR|A0A7E6D288|A0A7E6D288_9CHIR KKSSQSFLDEIHQHLWAYHQEASKHLSSQIPLIIQFFMLRSFGQK--LNNAMLQLLQDRE 621
TR|S7NSW7|S7NSW7_MYOBR         EDPLTCSLAEILQHLLAYRLEASNRLSSHIPLIIQFFVLRSYGQQ--LQKAMLQLLQDKE 642
TR|A0A1N7THY9|A0A1N7THY9_PIPPI EDPLSCSLSEILQHLMAYRLEASRRLSSHIPLIIQFFILQSNGQQ--LQKAMLQLLQDKD 618
TR|A0A1N7THZ0|A0A1N7THZ0_ROUAE SFPDVTPLDEILPHLTAYRQDASNRLSTHIPLVIQYFLLQSFGRQ--LEKSMLQLLQDKE 623
TR|A0A1N7THY7|A0A1N7THY7_HYPMN SSPDSSPLGEILPHLTAYRQDARDRLATHIPLVIQFFVLQSFGQQ--LQKSMLQLLQDKE 621
TR|A0A1N7THZ1|A0A1N7THZ1_STULI NSSSQSFLDEIHQHLWAYHQEASKRLSSQIPLIIQFFMLRSYGQK--LSSAMLQLLQDRE 619
TR|B2X026|B2X026_CHICK         SRQCPSFALEMVSHVKAYFTGASKRLSNQIPLIILSTVLHDFGNY--LQTSMLHLLQGKE 666
TR|A0A172PZ65|A0A172PZ65_ANSCY SNQNAIFIQEMISHTKAYFTGASKRLSNQIPLIILSAVLHDFGDN--LQTSMLHLLQERE 664
TR|A0A0Q3PUG8|A0A0Q3PUG8_AMAAE TQQEPYFVQELVSHTKAYFYGASKRLSNQIPLIILSA-FHDFADG--LQTTMLHLLQEKD 487
SP|Q4ADG6|MX1_PHOVI            VSSVNISLSEIFEHLLAYRQEATNRISSHIPLIIQYFILQAYGQK--LQKGMLQLLQDKD 621
SP|Q4ADG7|MX1_OTABY            VSSVNISLSDIFEHLLAYRQEATNRISSHIPLIIQYFILQVYGQK--LQKDMLLLLHDKD 620
TR|A0A2Y9G772|A0A2Y9G772_NEOSC VSSVNISLSEIFEHLLAYRQEATNRISSHIPLIIQYFILQVYGQK--LQKGMLQLLQDKD 622
TR|A0A6J2C7I6|A0A6J2C7I6_ZALCA VSSVNISLSDIFEHLLAYRQEATNRISSHIPLIIQYFILQVYGQK--LQKDMLLLLHDKD 620
```

```
TR|A0A2U3WDI8|A0A2U3WDI8_ODORO VSSVNISLSEIFEHLLAYRQEATNRLSSHIPLIIQYFILQVYGEK--LQKGMLQLLHDKD 620
TR|A0A1U7SNC8|A0A1U7SNC8_ALLSI -PDKSPSTVELACHLEAYFNGAGHRLSSLIPLIIQSFTIQDYGDK--LQNAMLQLLQEKQ 694
TR|A0A7M4FFF4|A0A7M4FFF4_CROPO PLHNQYSVKEMAYHLEAYFTGAGNRLSSQIPLIMQSFILQDYGDN--LQNAMMQLLQEKA 608
TR|A0A340WTU5|A0A340WTU5_LIPVE ---------------YCAHQEVSTRISSHTPLIIQFFVLKTFG-Q--LRKGMLQLLQNKD 482
TR|A0A6J3RAX5|A0A6J3RAX5_TURTR RHLSLPA-------------------SRH------------------------------ 523
TR|A0A341CI24|A0A341CI24_NEOAA ------------------------------------------------------------
TR|A0A2Y9LTL6|A0A2Y9LTL6_DELLE RHLSLPA-------------------SRH------------------------------ 518
TR|A0A383ZCI9|A0A383ZCI9_BALAS QSLSDPSIAEIFQHLIAYHQEVSTRISGHIPLIIQFFMLKTFGGQ--LQKGMLQLLQSKD 621
TR|V9KIB1|V9KIB1_CALMI         SKSKEESIQEMSTHVDTYYKIMSDRLADQVPLIIRYYMMQEFASN--LQVQMLELLQNRQ 589
TR|A0A674K7E1|A0A674K7E1_TERCA ASQKQYSIDEMAYHLNAYFKGAGNRLSTQIPLIIQSYVLQDYTEK--LQNAMLQLLQDKD 629
TR|A0A4D9DW37|A0A4D9DW37_9SAUR TVQNHPAVKDMTYHLEAYFSSASKRLSNQIPLIIQFYILQDFGDK--LQNSILQLLQEKE 721
TR|H2DQ54|H2DQ54_XENLA         PTPMQISCDEMHYHVQAYFKSLKERLSNQIPMIIQYYILHEFSEK--LQNQMMQLIQERE 586
TR|A0A0F6MUS9|A0A0F6MUS9_ANGJA STDSHATLQEMMVHLKSYYRIASQRLADQIPLVIRYLMLQESAAQ--LQREMLQTLQDKE 609
TR|Q98990|Q98990_SALSA         STDNHATLQEMMLHLKSYYRISSQRLADQIPMVIRYLVLQEFASQ--LQREMLQTLQEKD 587
TR|A0A674EGU5|A0A674EGU5_SALTR STDNHATLQEMMLHLKSYYRISSQRLADQIPMVIRYLVLQEFASQ--LQREMLQTLQEKD 594
SP|Q91192|MX1_ONCMY            STDNHATLQEMMLHLKSYYWISSQRLADQIPMVIRYLVLQEFASQ--LQREMLQTLQEKD 585
TR|A0A286K200|A0A286K200_MYLPI STNNNATLREMRLHLESYYSIASKRLSDQIPMVIRYLLLQEAALE--LQRNMLQLLHDKD 594
TR|A0A498MHV8|A0A498MHV8_LABRO GTDSSATLRGMRLHLESYYMIASKRLADQIPMVMRYLLLQEAALE--LQRNMLQLLQNKD 592
SP|Q7T2P0|MX1_ICTPU            HSNNRATLEELMRHLKSYYSIASKRLADQLPLVIRYLLLQESAAQ--LQREMLQLMQDKN 599
TR|A0A556TK66|A0A556TK66_BAGYA YSDNKATLEELTHHLKSYYSIASKRLADQLPLVIRYLLLQESAAQ--MQREMLQLLQDKS 567
TR|A0A3G6IID3|A0A3G6IID3_9TELE STDNNATLREMRLHLESYYSIASKRLSDQIPMVIRYLLLQEAALE--LQRNMLQLLHDKD 594
SP|Q8JH68|MXA_DANRE            GTDNHATLREMRLHLKSYYTIASKRLADQIPMVIRYMLLQEAALE--LQRNMLQLLQDKD 596
TR|A0A672MEY2|A0A672MEY2_SINGR GCALLDTRNLVPEKLVLYYEIIYQRLTDYVPMLIILFMLKDAAKT--LRHQMMELRNGAD 594
TR|Q7T2M3|Q7T2M3_CARAU         STGTHATLREMRLHLESYYTIASKRLADQIPMVIRYLLLQEAALE--LQRNMLQLLQDKD 599
TR|A0A673LWX8|A0A673LWX8_9TELE GTGNHATLREMRLHLESYYMIASKRLADQIPMVIRYLLLQEAALE--LQRNMLQLLQDKD 596
TR|A0A8B9KF29|A0A8B9KF29_ASTMX NTNNHATLQELMRHLKSYYGIASKRLADQVPLMIRYLLLQEAALQ--LQRDMLQLIQDKD 591


SP|P20591|MX1_HUMAN            TYSWLLKERSDTSDK------------RKFLKERL-------ARLTQARRRLAQF----- 660
TR|K4JBQ0|K4JBQ0_9PRIM         TYSWLLKERSDTSDK------------RKFLKERL-------ARLTQARRRLAQF----- 660
TR|H2QL18|H2QL18_PANTR         TYSWLLKERSDTSDK------------RKFLKERL-------ARLTQARRRLAQF----- 660
SP|A1E2I4|MX1_MACMU            TYSWLLKERSDTSDK------------RKFLKERL-------ARLTQARRRLAQF----- 659
SP|Q5R5G3|MX1_PONAB            TYSWLLKERGDTSDK------------RKFLKERL-------ARLTQARRRLAQF----- 660
TR|A0A6P8QPW8|A0A6P8QPW8_GEOSA EFEKLLEQQPDLVNQ------------RKRVKHRI-------QELRQARTLHQPFEFNFM 628
TR|U3EGN4|U3EGN4_CALJA         TYSWLLKERSDTSDK------------RKFLKERL-------ARLTQARRRLAQF----- 660
SP|Q9N0Y3|MX1_CANLF            TYSWLLKERSDTSDK------------RKFLKERL-------ARLAQARRRLAKF----- 655
TR|M3W8X0|M3W8X0_FELCA         TYSWLLKERSDTSDK------------RKFLKERL-------ARLAQARRRLAKF----- 658
TR|A0A667GNL8|A0A667GNL8_LYNCA TYSWLLKERSDTSDK------------RKFLKERL-------ARLAQARRRLAKF----- 658
TR|A0A6P6GZN4|A0A6P6GZN4_PUMCO ------------------------------------------------------------
TR|A0A6J1XXY6|A0A6J1XXY6_ACIJB TYSWLLKERSDTSDK------------RKFLKERL-------ARLAQARRRLAKF----- 658
TR|A0A6P4TS13|A0A6P4TS13_PANPR TYSWLLKERSDTSDK------------RKFLKERL-------ARLAQARRRLAKF----- 696
TR|A0A3Q7SCG7|A0A3Q7SCG7_VULVU TYSWLLKERSDTSDK------------RKFLKERL-------ARLAQARRRLAKF----- 694
TR|I3MJX6|I3MJX6_ICTTR         ACGWLLKERSDTSDK------------RKFLKERL-------SRLTQARHRLAKF----- 658
TR|G9KBY2|G9KBY2_MUSPF         TYNWLLKERSDTSDK------------RKFLKERL-------ARLGQARRRLAKF----- 655
TR|A0A2Y9KR88|A0A2Y9KR88_ENHLU TYNWLLKERSDTSDK------------RKFLKERL-------ARLAQARRHLAKF----- 656
SP|P09922|MX1B_MOUSE           KCSWFLEEQSDTREK------------KKFLKRRL-------LRLDEARQKLAKF----- 629
SP|P18588|MX1_RAT              KCNWFLTEQSDSREK------------KKFLKRRL-------LRLDEAQRKLAKF----- 650
TR|A0A6P6D2J7|A0A6P6D2J7_PTEVA QYDWLLKERSDTCDK------------RKFLKEQH-------RRLVQARHQLAKF----- 681
TR|A0A6J0DKA3|A0A6J0DKA3_PERMB NCSQLLAERSDITEK------------RKFLKERL-------SRLDQARHKLAIF----- 617
TR|G5C5C9|G5C5C9_HETGA         ASA-LSLRGSDHRDSVGTHPAPSLTGSGCFLGQASVACKARAGPLWGGTDPAVPFLSPGT 701
TR|A0A3Q0CWV5|A0A3Q0CWV5_MESAU NCNWLLAERKDTTEK------------RKFLKRRL-------SRLDQARHKLAKF----- 643
TR|A0A6I9K5X3|A0A6I9K5X3_CHRAS GYDWLLKERSDTSDK------------RKFLKERL-------WRLTQAKRRLAKF----- 661
SP|P79135|MX1_BOVIN            QYDWLLKERTDTRDK------------RKFLKERL-------ERLTRARQRLAKF----- 646
TR|A0A6P3H2K4|A0A6P3H2K4_BISBI QYDWLLKERTDTRDK------------RKFLKERL-------ERLTRARQRLAKF----- 652
TR|L8IPR7|L8IPR7_9CETA         QYDWLLKERTDTRDK------------RKFLKERL-------ERLTRARQRLAKF----- 649
SP|P27594|MX1_PIG              QYDWLLRERSDTSDK------------RKFLKERL-------MRLTQARRRLAKF----- 661
TR|A0A1U7U9K3|A0A1U7U9K3_CARSF TYSWLLKERSDTSEK------------RRFLKERL-------ARLTQARRRLAKF----- 624
SP|P33237|MX1_SHEEP            QYDWLLKERTDTRDK------------RKFLKERL-------ERLSRARQRLAKF----- 652
TR|A0A7D4WPX4|A0A7D4WPX4_CAPHI QYDWLLKERTDTRDK------------RKFLKERL-------ERLSRARQRLAKF----- 652
SP|Q28379|MX1_HORSE            TYDWLLKERNDTCDK------------RKFLKERL-------ARLAQARRRLAKF----- 658
TR|A0A6I9I6Y4|A0A6I9I6Y4_VICPA EYDWLLKERSHTSDK------------RKFLKERL-------SRLTQARRRLAKF----- 660
TR|A0A6P5BYP3|A0A6P5BYP3_BOSIN QYDWLLKERTDTRDK------------RKFLKERL-------ERLTRARQRLAKF----- 649
TR|S9XPZ9|S9XPZ9_CAMFR         EYDWLLKERSHTSDK------------RKFLKERL-------SRLTQARRRLAKF----- 668
TR|A0A384DMX3|A0A384DMX3_URSMA TYNWLLKERSDTSDK------------RKFLKERL-------ARLSQARRRLAKF----- 658
TR|A0A3Q7W4X3|A0A3Q7W4X3_URSAR TYNWLLKERSDTSDK------------RKFLKERL-------ARLTQARRRLAKF----- 658
TR|A0A7E6D288|A0A7E6D288_9CHIR NYNLLLTERSDTRDK------------RKFLKEQL-------VRLTKARQELSKF----- 657
TR|S7NSW7|S7NSW7_MYOBR         KYDMLLKEHSDTSDK------------RKFLKERL-------ARLTKARRRLAQF----- 678
TR|A0A1N7THY9|A0A1N7THY9_PIPPI KYSWLLKERGDTSDK------------RRFLKERL-------ARLTKARRRLAQF----- 654
TR|A0A1N7THZ0|A0A1N7THZ0_ROUAE QYGWLLRERSDTCEK------------RKLLKEQH-------RRLVAARHRLAQF----- 659
TR|A0A1N7THY7|A0A1N7THY7_HYPMN QYGWLLRERSDTCEK------------RKRLKEQH-------RRLVQARHRLAQF----- 657
TR|A0A1N7THZ1|A0A1N7THZ1_STULI SYNSLLAERSDTSDK------------RKFLKEQL-------ARLTKARRQLSKF----- 655
TR|B2X026|B2X026_CHICK         EINYLLQEDHEANQ-------------QKLLTSRI-------SHLNKAYQYLVDFKSL-- 705
TR|A0A172PZ65|A0A172PZ65_ANSCY QLNILLQEDSEAARI------------RNYLSGRV-------NRLSKAYQCLKTFSCL-- 703
TR|A0A0Q3PUG8|A0A0Q3PUG8_AMAAE KLSHLLQEDSEAAKQ------------KIYLIERA-------DRLTKARQELRNFTSL-- 526
SP|Q4ADG6|MX1_PHOVI            TYNWLLKERSDTSDK------------RKFLKERL-------SRLAQARRRLAKF----- 657
SP|Q4ADG7|MX1_OTABY            THNWLLKERTDTRDK------------RKLLKERL-------ARLAQARRRLAKF----- 656
TR|A0A2Y9G772|A0A2Y9G772_NEOSC TYNWLLKEHSDTSDK------------RKFLKERL-------LRLAQARRRLAKF----- 658
TR|A0A6J2C7I6|A0A6J2C7I6_ZALCA THNWLLKERSDTSDK------------RKLLKERL-------ARLAQARRRLAKF----- 656
TR|A0A2U3WDI8|A0A2U3WDI8_ODORO TYNWLLKERSDTSDK------------RKFLKERL-------ARLAQARRRLAKF----- 656
TR|A0A1U7SNC8|A0A1U7SNC8_ALLSI LLDCLLMEHKDVAAE------------RNFLTQQI-------NRLRKAQLYLLKVSV--- 732
TR|A0A7M4FFF4|A0A7M4FFF4_CROPO QLGFFLEERKEVADQ------------RNFLTERI-------HRLTKARQHLLYLLHRLR 649
TR|A0A340WTU5|A0A340WTU5_LIPVE KYDRLLKERSDTSNR------------RKLLREPL-------QQLTRA--RLAKF----- 516
TR|A0A6J3RAX5|A0A6J3RAX5_TURTR ------------------------------------------------------------
TR|A0A341CI24|A0A341CI24_NEOAA ------------------------------------------------------------
TR|A0A2Y9LTL6|A0A2Y9LTL6_DELLE ------------------------------------------------------------
TR|A0A383ZCI9|A0A383ZCI9_BALAS EYDWLLKERSDTSDR------------RKFLRERL-------ERLTRARRRLAKF----- 657
TR|V9KIB1|V9KIB1_CALMI         QIEEYLEEEKTIAEK------------RRTLKNKL-------DRLSKAQVHLLDLA---- 626
TR|A0A674K7E1|A0A674K7E1_TERCA KFSILLLEKKDAATE------------RKNVKERI-------KRLTQARQRLAKFPG--- 667
```

```
TR|A0A4D9DW37|A0A4D9DW37_9SAUR KLSFFLQERKDAADQ------------RHFLSGRI-------HRLTQARLRLMKSSVL-- 760
TR|H2DQ54|H2DQ54_XENLA         KLNTLLAEKTDITRV------------RDNLSNRI-------ERLTAASHKLAKFHC--- 624
TR|A0A0F6MUS9|A0A0F6MUS9_ANGJA NVELLLKEDYDIGSK------------RAALQSRL-------KRLTEAHNYLMKF----- 645
TR|Q98990|Q98990_SALSA         NIEQLLKEDFDIGSK------------RAALQNKL-------KRLMKARSYLVEF----- 623
TR|A0A674EGU5|A0A674EGU5_SALTR NIEQLLKEDFDIGSK------------RAALQNKL-------KRLMKARSYLVEF----- 630
SP|Q91192|MX1_ONCMY           NIEQLLKEDIDIGSK------------RAALQSKL-------KRLMKARSYLVEF----- 621
TR|A0A286K200|A0A286K200_MYLPI GVDYLLKEDCDIGQK------------RESLLSRQ-------KRLMKARSLLVTY----- 630
TR|A0A498MHV8|A0A498MHV8_LABRO NVDDLLKEDFDIGQK------------RESLLSRQ-------KRLMKARGLLVTF----- 628
SP|Q7T2P0|MX1_ICTPU           AIDHLLKEDHDIGNK------------RNNLQSRQ-------KRLMEARNYLVKF----- 635
TR|A0A556TK66|A0A556TK66_BAGYA STDQLLKEDHDIGNK------------RSSLQSRQ-------KRLTEARNYLVKF----- 603
TR|A0A3G6IID3|A0A3G6IID3_9TELE CVDFLLKEDFDIGQK------------RESLLSRQ-------NRLMKARSLLVTY----- 630
SP|Q8JH68|MXA_DANRE           GVDNLLKEDCDIGQK------------RENLLSRQ-------TRLIEGTQPLGHLLEVTF 637
TR|A0A672MEY2|A0A672MEY2_SINGR VV-KLLSEDSEHGRM------------RADLKQRL-------ERLTQAQDLISNRL---- 630
TR|Q7T2M3|Q7T2M3_CARAU         GVDILLKEDFDIGQK------------RESLLSRQ-------KRLMKARSLLVTF----- 635
TR|A0A673LWX8|A0A673LWX8_9TELE GIDNLLKEDSDIGQK------------RENLLKRQ-------KRLMKARSLLVTF----- 632
TR|A0A8B9KF29|A0A8B9KF29_ASTMX CTETLLKEDHDIGNK------------RANLLSRQ-------KRLMKAHSYLFLVNVTSD 632


SP|P20591|MX1_HUMAN           PG---------------------------------------------------------- 662
TR|K4JBQ0|K4JBQ0_9PRIM         PG---------------------------------------------------------- 662
TR|H2QL18|H2QL18_PANTR         PG---------------------------------------------------------- 662
SP|A1E2I4|MX1_MACMU           PG---------------------------------------------------------- 661
SP|Q5R5G3|MX1_PONAB           PG---------------------------------------------------------- 662
TR|A0A6P8QPW8|A0A6P8QPW8_GEOSA -DAALANGIYEIPIPSSTADLPFSFCSDNLANGAEELPLTQPTAKQCTKSRKKDATGATN 687
TR|U3EGN4|U3EGN4_CALJA         PG---------------------------------------------------------- 662
SP|Q9N0Y3|MX1_CANLF           PG---------------------------------------------------------- 657
TR|M3W8X0|M3W8X0_FELCA         PG---------------------------------------------------------- 660
TR|A0A667GNL8|A0A667GNL8_LYNCA PG---------------------------------------------------------- 660
TR|A0A6P6GZN4|A0A6P6GZN4_PUMCO ------------------------------------------------------------
TR|A0A6J1XXY6|A0A6J1XXY6_ACIJB PG---------------------------------------------------------- 660
TR|A0A6P4TS13|A0A6P4TS13_PANPR PG---------------------------------------------------------- 698
TR|A0A3Q7SCG7|A0A3Q7SCG7_VULVU PG---------------------------------------------------------- 696
TR|I3MJX6|I3MJX6_ICTTR         PG---------------------------------------------------------- 660
TR|G9KBY2|G9KBY2_MUSPF         PG---------------------------------------------------------- 657
TR|A0A2Y9KR88|A0A2Y9KR88_ENHLU PG---------------------------------------------------------- 658
SP|P09922|MX1B_MOUSE          SD---------------------------------------------------------- 631
SP|P18588|MX1_RAT             SN---------------------------------------------------------- 652
TR|A0A6P6D2J7|A0A6P6D2J7_PTEVA PG---------------------------------------------------------- 683
TR|A0A6J0DKA3|A0A6J0DKA3_PERMB SY---------------------------------------------------------- 619
TR|G5C5C9|G5C5C9_HETGA         PGSPWAP------------------FRQQTRKGAEG------------------------ 719
TR|A0A3Q0CWV5|A0A3Q0CWV5_MESAU SY---------------------------------------------------------- 645
TR|A0A6I9K5X3|A0A6I9K5X3_CHRAS PG---------------------------------------------------------- 663
SP|P79135|MX1_BOVIN           PG---------------------------------------------------------- 648
TR|A0A6P3H2K4|A0A6P3H2K4_BISBI PG---------------------------------------------------------- 654
TR|L8IPR7|L8IPR7_9CETA         PG---------------------------------------------------------- 651
SP|P27594|MX1_PIG             PG---------------------------------------------------------- 663
TR|A0A1U7U9K3|A0A1U7U9K3_CARSF PG---------------------------------------------------------- 626
SP|P33237|MX1_SHEEP           PG---------------------------------------------------------- 654
TR|A0A7D4WPX4|A0A7D4WPX4_CAPHI PG---------------------------------------------------------- 654
SP|Q28379|MX1_HORSE           PG---------------------------------------------------------- 660
TR|A0A6I9I6Y4|A0A6I9I6Y4_VICPA PG---------------------------------------------------------- 662
TR|A0A6P5BYP3|A0A6P5BYP3_BOSIN PG---------------------------------------------------------- 651
TR|S9XPZ9|S9XPZ9_CAMFR         PG---------------------------------------------------------- 670
TR|A0A384DMX3|A0A384DMX3_URSMA PG---------------------------------------------------------- 660
TR|A0A3Q7W4X3|A0A3Q7W4X3_URSAR PG---------------------------------------------------------- 660
TR|A0A7E6D288|A0A7E6D288_9CHIR PS---------------------------------------------------------- 659
TR|S7NSW7|S7NSW7_MYOBR         PG---------------------------------------------------------- 680
TR|A0A1N7THY9|A0A1N7THY9_PIPPI PG---------------------------------------------------------- 656
TR|A0A1N7THZ0|A0A1N7THZ0_ROUAE PG---------------------------------------------------------- 661
TR|A0A1N7THY7|A0A1N7THY7_HYPMN PG---------------------------------------------------------- 659
TR|A0A1N7THZ1|A0A1N7THZ1_STULI PG---------------------------------------------------------- 657
TR|B2X026|B2X026_CHICK         ------------------------------------------------------------
TR|A0A172PZ65|A0A172PZ65_ANSCY ------------------------------------------------------------
TR|A0A0Q3PUG8|A0A0Q3PUG8_AMAAE ------------------------------------------------------------
SP|Q4ADG6|MX1_PHOVI           PG---------------------------------------------------------- 659
SP|Q4ADG7|MX1_OTABY           PG---------------------------------------------------------- 658
TR|A0A2Y9G772|A0A2Y9G772_NEOSC PS---------------------------------------------------------- 660
TR|A0A6J2C7I6|A0A6J2C7I6_ZALCA PG---------------------------------------------------------- 658
TR|A0A2U3WDI8|A0A2U3WDI8_ODORO PG---------------------------------------------------------- 658
TR|A0A1U7SNC8|A0A1U7SNC8_ALLSI ------------------------------------------------------------
TR|A0A7M4FFF4|A0A7M4FFF4_CROPO VMSTTLPK-----------------LSK-------EL---PAWSHKCMPSPEVWK----- 677
TR|A0A340WTU5|A0A340WTU5_LIPVE PG---------------------------------------------------------- 518
TR|A0A6J3RAX5|A0A6J3RAX5_TURTR ------------------------------------------------------------
TR|A0A341CI24|A0A341CI24_NEOAA ------------------------------------------------------------
TR|A0A2Y9LTL6|A0A2Y9LTL6_DELLE ------------------------------------------------------------
TR|A0A383ZCI9|A0A383ZCI9_BALAS PG---------------------------------------------------------- 659
TR|V9KIB1|V9KIB1_CALMI         ------------------------------------------------------------
TR|A0A674K7E1|A0A674K7E1_TERCA ------------------------------------------------------------
TR|A0A4D9DW37|A0A4D9DW37_9SAUR ------------------------------------------------------------
TR|H2DQ54|H2DQ54_XENLA         ------------------------------------------------------------
TR|A0A0F6MUS9|A0A0F6MUS9_ANGJA ------------------------------------------------------------
TR|Q98990|Q98990_SALSA         ------------------------------------------------------------
TR|A0A674EGU5|A0A674EGU5_SALTR ------------------------------------------------------------
SP|Q91192|MX1_ONCMY           ------------------------------------------------------------
TR|A0A286K200|A0A286K200_MYLPI ------------------------------------------------------------
TR|A0A498MHV8|A0A498MHV8_LABRO ------------------------------------------------------------
SP|Q7T2P0|MX1_ICTPU           ------------------------------------------------------------
TR|A0A556TK66|A0A556TK66_BAGYA ------------------------------------------------------------
```

```
TR|A0A3G6IID3|A0A3G6IID3_9TELE ------------------------------------------------------------
SP|Q8JH68|MXA_DANRE            IDY----------------------CNILMQ----------------------------- 646
TR|A0A672MEY2|A0A672MEY2_SINGR ------------------------------------------------------------
TR|Q7T2M3|Q7T2M3_CARAU         ------------------------------------------------------------
TR|A0A673LWX8|A0A673LWX8_9TELE ------------------------------------------------------------
TR|A0A8B9KF29|A0A8B9KF29_ASTMX RRTGLKNN-----------------VQVNLKQS--QK---EDKTTDCLKTKH-------- 662


SP|P20591|MX1_HUMAN            -------------------------------------------------
TR|K4JBQ0|K4JBQ0_9PRIM         -------------------------------------------------
TR|H2QL18|H2QL18_PANTR         -------------------------------------------------
SP|A1E2I4|MX1_MACMU            -------------------------------------------------
SP|Q5R5G3|MX1_PONAB            -------------------------------------------------
TR|A0A6P8QPW8|A0A6P8QPW8_GEOSA TSINQKQQGPSNSSSSGPTGKVNASSIRPPTLIFHSNNQPFNSELPSSP 736
TR|U3EGN4|U3EGN4_CALJA         -------------------------------------------------
SP|Q9N0Y3|MX1_CANLF            -------------------------------------------------
TR|M3W8X0|M3W8X0_FELCA         -------------------------------------------------
TR|A0A667GNL8|A0A667GNL8_LYNCA -------------------------------------------------
TR|A0A6P6GZN4|A0A6P6GZN4_PUMCO -------------------------------------------------
TR|A0A6J1XXY6|A0A6J1XXY6_ACIJB -------------------------------------------------
TR|A0A6P4TS13|A0A6P4TS13_PANPR -------------------------------------------------
TR|A0A3Q7SCG7|A0A3Q7SCG7_VULVU -------------------------------------------------
TR|I3MJX6|I3MJX6_ICTTR         -------------------------------------------------
TR|G9KBY2|G9KBY2_MUSPF         -------------------------------------------------
TR|A0A2Y9KR88|A0A2Y9KR88_ENHLU -------------------------------------------------
SP|P09922|MX1B_MOUSE           -------------------------------------------------
SP|P18588|MX1_RAT              -------------------------------------------------
TR|A0A6P6D2J7|A0A6P6D2J7_PTEVA -------------------------------------------------
TR|A0A6J0DKA3|A0A6J0DKA3_PERMB -------------------------------------------------
TR|G5C5C9|G5C5C9_HETGA         -------------------------------------------------
TR|A0A3Q0CWV5|A0A3Q0CWV5_MESAU -------------------------------------------------
TR|A0A6I9K5X3|A0A6I9K5X3_CHRAS -------------------------------------------------
SP|P79135|MX1_BOVIN            -------------------------------------------------
TR|A0A6P3H2K4|A0A6P3H2K4_BISBI -------------------------------------------------
TR|L8IPR7|L8IPR7_9CETA         -------------------------------------------------
SP|P27594|MX1_PIG              -------------------------------------------------
TR|A0A1U7U9K3|A0A1U7U9K3_CARSF -------------------------------------------------
SP|P33237|MX1_SHEEP            -------------------------------------------------
TR|A0A7D4WPX4|A0A7D4WPX4_CAPHI -------------------------------------------------
SP|Q28379|MX1_HORSE            -------------------------------------------------
TR|A0A6I9I6Y4|A0A6I9I6Y4_VICPA -------------------------------------------------
TR|A0A6P5BYP3|A0A6P5BYP3_BOSIN -------------------------------------------------
TR|S9XPZ9|S9XPZ9_CAMFR         -------------------------------------------------
TR|A0A384DMX3|A0A384DMX3_URSMA -------------------------------------------------
TR|A0A3Q7W4X3|A0A3Q7W4X3_URSAR -------------------------------------------------
TR|A0A7E6D288|A0A7E6D288_9CHIR -------------------------------------------------
TR|S7NSW7|S7NSW7_MYOBR         -------------------------------------------------
TR|A0A1N7THY9|A0A1N7THY9_PIPPI -------------------------------------------------
TR|A0A1N7THZ0|A0A1N7THZ0_ROUAE -------------------------------------------------
TR|A0A1N7THY7|A0A1N7THY7_HYPMN -------------------------------------------------
TR|A0A1N7THZ1|A0A1N7THZ1_STULI -------------------------------------------------
TR|B2X026|B2X026_CHICK         -------------------------------------------------
TR|A0A172PZ65|A0A172PZ65_ANSCY -------------------------------------------------
TR|A0A0Q3PUG8|A0A0Q3PUG8_AMAAE -------------------------------------------------
SP|Q4ADG6|MX1_PHOVI            -------------------------------------------------
SP|Q4ADG7|MX1_OTABY            -------------------------------------------------
TR|A0A2Y9G772|A0A2Y9G772_NEOSC -------------------------------------------------
TR|A0A6J2C7I6|A0A6J2C7I6_ZALCA -------------------------------------------------
TR|A0A2U3WDI8|A0A2U3WDI8_ODORO -------------------------------------------------
TR|A0A1U7SNC8|A0A1U7SNC8_ALLSI -------------------------------------------------
TR|A0A7M4FFF4|A0A7M4FFF4_CROPO -------------------------------------------------
TR|A0A340WTU5|A0A340WTU5_LIPVE -------------------------------------------------
TR|A0A6J3RAX5|A0A6J3RAX5_TURTR -------------------------------------------------
TR|A0A341CI24|A0A341CI24_NEOAA -------------------------------------------------
TR|A0A2Y9LTL6|A0A2Y9LTL6_DELLE -------------------------------------------------
TR|A0A383ZCI9|A0A383ZCI9_BALAS -------------------------------------------------
TR|V9KIB1|V9KIB1_CALMI         -------------------------------------------------
TR|A0A674K7E1|A0A674K7E1_TERCA -------------------------------------------------
TR|A0A4D9DW37|A0A4D9DW37_9SAUR -------------------------------------------------
TR|H2DQ54|H2DQ54_XENLA         -------------------------------------------------
TR|A0A0F6MUS9|A0A0F6MUS9_ANGJA -------------------------------------------------
TR|Q98990|Q98990_SALSA         -------------------------------------------------
TR|A0A674EGU5|A0A674EGU5_SALTR -------------------------------------------------
SP|Q91192|MX1_ONCMY            -------------------------------------------------
TR|A0A286K200|A0A286K200_MYLPI -------------------------------------------------
TR|A0A498MHV8|A0A498MHV8_LABRO -------------------------------------------------
SP|Q7T2P0|MX1_ICTPU            -------------------------------------------------
TR|A0A556TK66|A0A556TK66_BAGYA -------------------------------------------------
TR|A0A3G6IID3|A0A3G6IID3_9TELE -------------------------------------------------
SP|Q8JH68|MXA_DANRE            -------------------------------------------------
TR|A0A672MEY2|A0A672MEY2_SINGR -------------------------------------------------
TR|Q7T2M3|Q7T2M3_CARAU         -------------------------------------------------
TR|A0A673LWX8|A0A673LWX8_9TELE -------------------------------------------------
TR|A0A8B9KF29|A0A8B9KF29_ASTMX -------------------------------------------------
```
