## Supplementary Figure 2 for "The N-terminal domain of MX1 proteins is essential for their antiviral activity against different families of RNA viruses"

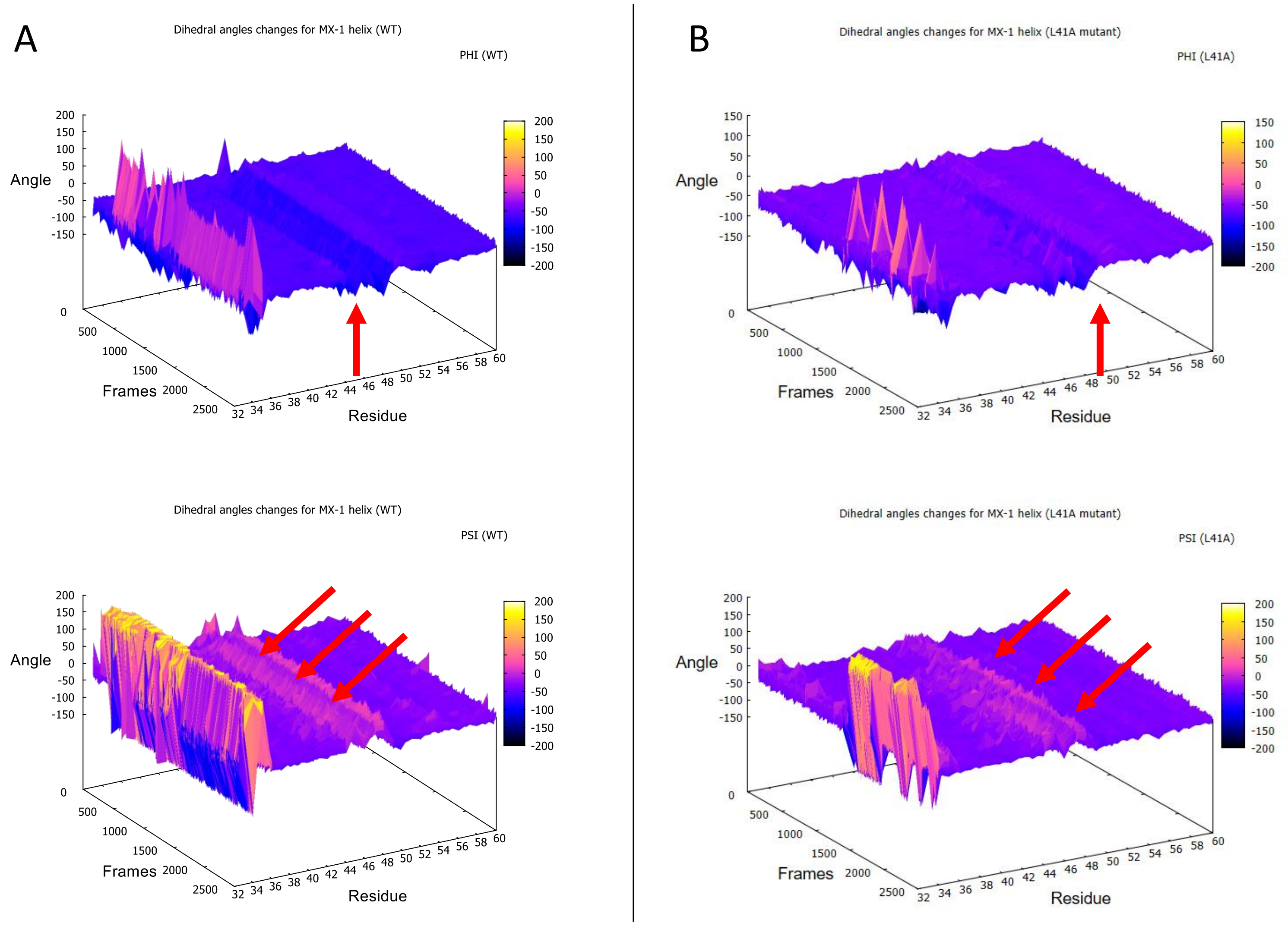


***<u>Sup. Fig. 2</u>***: **Secondary structure of the N-terminal BSE α-helix behave identically between WT and L41A mutant.** Phi and Psi dihedral angles from HsMX1 residues (spanning from 33 to 60) were monitored all along the simulation for WT (**A**) and L41A mutant (**B**). For clarity, only one frame over ten was used for plotting (0.2 ns / frame) and red arrows indicate the location of the α-helix brake (or turn) encountered in both constructs.
